# Shared brain basis of aggression in clinical, forensic, and healthy samples: A meta-analysis

**DOI:** 10.1101/2025.02.17.638609

**Authors:** Harri Harju, Jouni Tuisku, Lauri Nummenmaa

**Affiliations:** Turku PET Centre; Department of Psychology, University of Turku; Turku University Hospital, University of Turku

## Abstract

**Background:** Aggression, violence, and antisocial behaviour constitute a large-scale societal problem. Aggression is common in incarcerated offenders and psychiatric conditions, but also healthy and noninstitutionalized populations vary in violent and aggressive behaviour. The brain basis of aggression has been studied extensively in the past, but the similarities between criminal, pathological and everyday aggression in the brain remain elusive.

**Methods:** We conducted an activation likelihood estimation (ALE) meta-analysis of 406 neuroimaging studies with 28 968 subjects using structural magnetic resonance imaging (MRI), functional magnetic resonance imaging (fMRI), positron emission tomography (PET) and single photon emission tomography (SPECT). The included studies had either i) measured haemodynamic responses during aggression-related functional tasks, ii) compared the brain structure, molecular organization, or function between aggressive forensic and psychiatric populations and control groups, or iii) addressed the effects of trait aggression on brain structure or function.

**Results:** Aggression was consistently associated with altered function and structure in the amygdala, hippocampus, basal ganglia, anterior cingulate cortex, and the dorsolateral and orbitofrontal cortices. Functional coactivation analysis suggested that these regions are most consistently associated with emotional and reward function as well as their regulation. The results were comparable in healthy subjects as well as forensic and psychiatric populations.

**Conclusions:** Aggression is linked with alterations in multiple neurocognitive systems forming a common network for aggressiveness. Particularly the neural systems implicated in reward, emotions and regulation were commonly associated with aggression. The established network is involved in the whole continuum of aggression from benign variations in healthy volunteers to forensic subjects and violent clinical populations, suggesting a common aggression network whose severe perturbations may be linked with criminal behaviour or pathological aggression.

## Introduction

Humans have specialized neural circuits for aggression because we need them. Aggression is defined as behaviour with the intent of harming another person who does not wish to be harmed (Baron & Richardson, 1994). Although human societies flourish due to our ability to cooperate and establish complex social alliances, aggression can also be used for gaining a lot: territory, food, mating partners and other resources (Gómez et al., 2016). Consequently, aggression is highly heritable and about half of its variance can be accredited to genetic factors (Ferguson, 2010; Miles & Carey, 1997; Rhee & Waldman, 2002). Aggression is a major societal problem throughout the world. Violent acts excluding armed conflict account for 1.5 million deaths worldwide annually (WHO, 2007). Hostility and aggression are also common symptomatic manifestations of numerous psychiatric conditions. Aggression is particularly common in individuals with antisocial personality disorders (ASPDs), and consequently ASPDs are prevalent in incarcerated populations (Black et al., 2010; Fridell et al., 2008). Aggression and violence also manifest in other psychiatric conditions ranging from psychoses to substance use disorders aggravating the psychosocial problems and complicating treatment (Fazel et al., 2018; Witt et al., 2013). Clinical and forensic samples aside, aggression is also common in otherwise well-functioning community samples (Jones & Paulhus, 2014). For example, bullying is a global phenomenon both in school and in workplaces, with almost one in three students having experienced bullying by their peers in the past month (UNESCO, 2018) and more than one in five persons in employment reporting experiences of violence and harassment in their working lives (ILO, 2022). Bullying leads to severe consequences for the bullied in terms of physical and psychological wellbeing (Geoffroy et al., 2016) and ability to learn (Rueger & Jenkins, 2014). Hence, aggression is not only problematic in the most severe violent cases, but throughout the continuum from everyday aggression to pathological.

On neural level, aggression is linked with dysfunction in the brain circuits involved in threat detection, emotion regulation, and inhibition resulting in an imbalance of top-down control and bottom-up affective drive (Siever, 2008). The candidate systems underlying aggression thus localize in the amygdala, orbitofrontal cortex, ventromedial and dorsolateral prefrontal cortices, and anterior cingulum (Davidson et al., 2000). Accordingly, meta-analyses have revealed decreased grey matter volume in the insula, hippocampus, putamen, pallidum, and caudate nucleus across aggressive samples versus controls (e.g. Johanson et al., 2020; Rogers & De Brito, 2016). In terms of neurotransmitters and neuromodulators, the most important pathways for aggression involve the serotonin, dopamine, opioid, and possibly also the endocannabinoid system (Duke et al., 2013; Kolla & Mishra, 2018; Lukkarinen et al., 2024; Nummenmaa & Tuominen, 2018; Seo et al., 2008). Finally, some studies suggest that aberrant social cognitive processes may also link with aggression, as hypoactivation linked to social processes has also been found in the fusiform gyrus, amygdala, inferior prefrontal gyrus, superior temporal sulcus (Mier et al., 2014) and the frontal pole (Aoki et al., 2014) as well as in components of the putative mirror neuron system with antisocial groups (Johanson et al., 2020).

An important yet unresolved question regarding the brain basis of aggression is the functioning and perturbations of the human aggression circuit across populations. Given that aggression occurs across healthy populations as well as in psychiatric and forensic populations, it would be imperative to understand whether i) there exists a common aggression network supporting everyday aggression in healthy subjects and its perturbations lead to pathological or criminal aggression, or whether ii) pathological and criminal aggression are linked with alterations distinct from the common aggression network. Moreover, the evidence regarding the role of functional (i.e. evoked responses), structural (i.e. grey matter concentrations) and molecular (i.e. neuroreceptor densities) changes in aggression remains elusive. Current studies on the neural basis of aggression cannot however answer these questions, since most studies including meta-analyses focus on specific subject group (such as on children with conduct disorder (Rogers & De Brito, 2016), imaging method (such as voxel-based morphometry (VBM); (Aoki et al., 2014)), or specific brain regions (such as the prefrontal cortex (Yang & Raine, 2009)).

Here we used an integrative meta-analytic approach for mapping the brain basis of aggression across multiple levels. We combined data from functional, structural, and molecular imaging studies investigating the neurobiological basis of aggression in i) aggressive and violent psychiatric patients, ii) incarcerated violent offenders, and iii) healthy non-institutionalized populations. We found that a common set of limbic and paralimbic regions was involved in the continuum of aggression from benign variations in healthy volunteers to forensic subjects and clinical populations, and that the same network could be identified with functional, structural and molecular imaging techniques. Our results thus reveal a common aggression network whose perturbations are linked with criminal behaviour or pathological aggression.

## Methods

### Data Selection

We used a structured search protocol for establishing the study database (**Figure 1**). We searched the Scopus database for articles published between 1990–2021 that had used structural magnetic resonance imaging (MRI), functional magnetic resonance imaging (fMRI), positron emission tomography (PET) or single photon emission computed tomography (SPECT) for addressing the brain basis of aggression. The studies were included if they had i) measured brain responses to aggression, ii) compared brain structure or function between control subjects and an aggressive group (such as violent prisoners or violent psychotic patients) or iii) addressed the effects of trait aggression on brain structure or function. Studies referred to by the identified papers and review articles were also considered. The studies were sorted into three main categories based on the subjects: i) those with only healthy volunteers, ii) those with psychiatric patients and iii) those with forensic samples.

**Figure 1.**
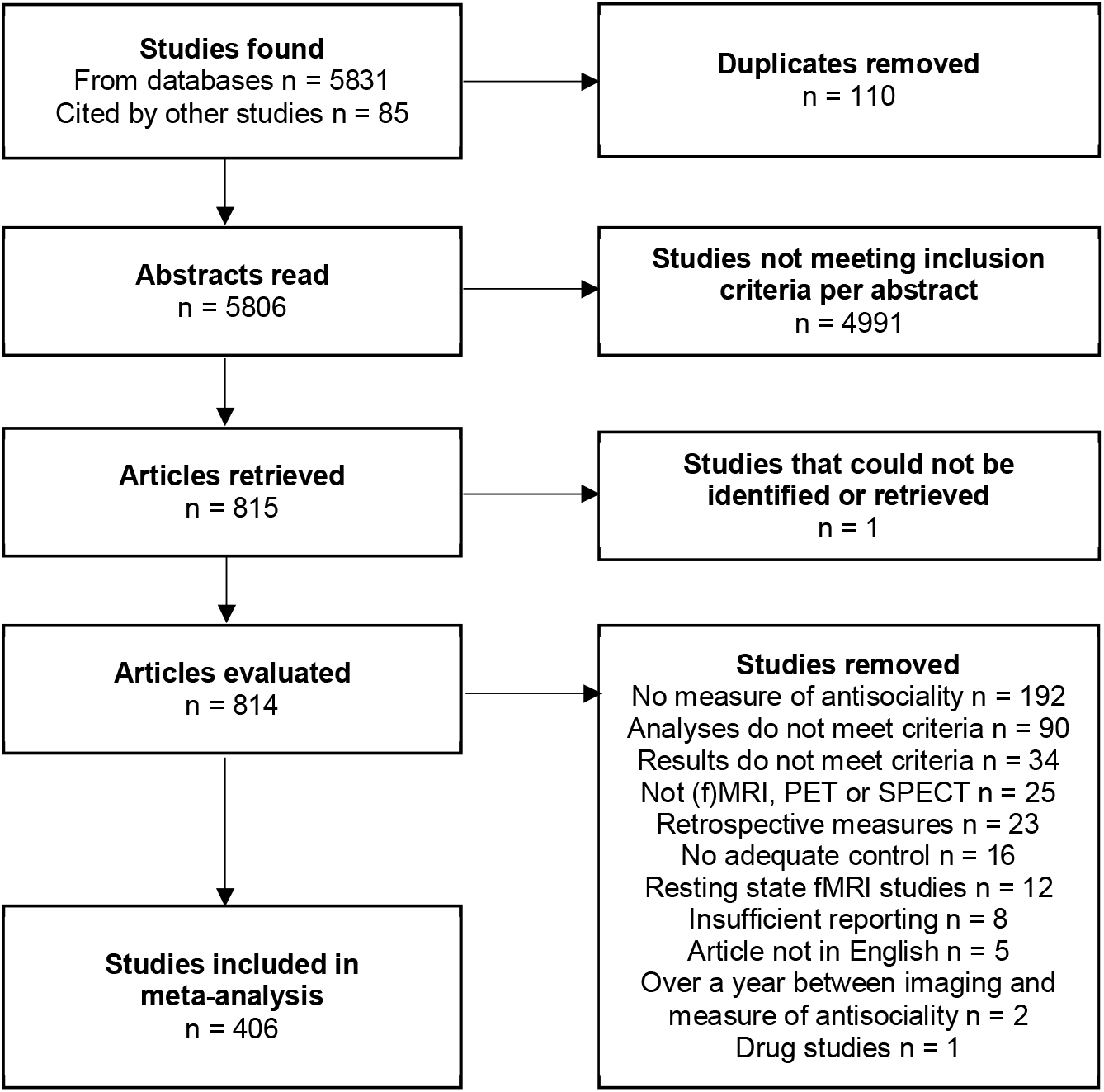
Flowchart of the study selection for the meta-analysis

Only studies reporting the results with stereotaxic coordinates in a standard reference space (Talairach-Tournoux or Montreal Neurological Institute [MNI]) or anatomical regions of interest were included. Studies measuring only white matter properties or resting state brain activity were excluded. Electromagnetic (MEG / EEG) experiments, diffusion tensor imaging (DTI) and arterial spin labelling studies were also not considered.

The search (**Figure 1**) returned 5831 articles. After exclusions, a total of 406 articles were found to be eligible for inclusion (**Table S1**). The following details were extracted from each article: imaging method and details (fMRI-task, such as the Taylor aggression paradigm, or MRI-technique, such as VBM, or the tracer), the number of subjects and their mean age, sex, and reported medical conditions and medications as a whole and grouped into experimental and control groups. If coordinates for results foci for the contrasts were available these were extracted. For studies reporting effects in anatomical regions of interest rather than coordinates (typically PET and SPECT studies), MNI coordinates were assigned with in-house MATLAB functions by first selecting the closest AAL3 atlas label for each ROI, as measured by Levenshtein distance, and then calculating the center of mass for the chosen AAL3 label. The selections were subsequently validated manually. Because this approach would otherwise have yielded the same coordinate for each study analysing a given ROI (such as the amygdala), we added random Gaussian noise (sigma 2 mm) to the resulting coordinates to avoid spatial bias in the ROI-based results. Before analysis, foci reported in the Talairach space were converted to MNI space using the mni2tal transform.

Results were considered duplicate if the subject group was a subgroup of another study’s subjects and the grouping was not done based on a new or relevant factor. These results were removed from the database. Correlational results were considered duplicate if the main effect had already been reported via a group comparison of the same subjects (e.g. correlation with PCL-R scores in addition to a comparison between high and low scores) or with another, higher order correlation (e.g. correlation with PCL-R subscale scores in addition to the PCL-R total score). Some studies had potentially overlapping samples, but these results were not removed from the database if the overlap could not be explicitly resolved from the reports. Altogether this process yielded 5047 foci from 1076 contrasts (**Table 1**).

**Table 1.** Summary of the studies included in the ALE analyses.

| Analysis | Studies | Subjects | Contrasts | Foci |
| --- | --- | --- | --- | --- |
| **Primary analyses** |  |  |  |  |
| Omnibus effect | 406 | 28968 | 1076 | 5047 |
| Offenders vs. controls | 106 | 8029 | 321 | 1684 |
| Patients vs. controls | 186 | 10934 | 479 | 2038 |
| Healthy volunteers | 123 | 10266 | 276 | 1325 |
| **Modality-specific analyses** |  |  |  |  |
| fMRI studies | 229 | 13538 | 682 | 3239 |
| Structural MRI studies | 124 | 13758 | 275 | 1326 |
| PET and SPECT studies | 58 | 1919 | 119 | 482 |
| **fMRI subanalyses** |  |  |  |  |
| Aggression | 39 | 1556 | 93 | 680 |
| Other emotions | 115 | 6377 | 376 | 1700 |
| Executive functions | 43 | 4040 | 118 | 433 |
| Other tasks | 33 | 1668 | 95 | 426 |
| **Structural MRI subanalyses** |  |  |  |  |
| Antisocial patients | 53 | 3181 | 111 | 725 |
| Psychotic patients | 16 | 786 | 29 | 93 |
| Other patients | 22 | 3351 | 52 | 206 |

### Activation Likelihood Estimation

Summary maps were generated using the Activation Likelihood Estimation based meta-analysis of neuroimaging data (Eickhoff et al., 2012) as implemented in NiMARE v0.0.7 (Salo et al., 2021, 2023) .Significance level was estimated using Monte Carlo method with 10 000 iterations and the results were thresholded at *p* < 0.01, FWE corrected at cluster level. **Table 1** shows the number of studies, subjects, contrasts, and foci for the conducted ALE analyses. The following four primary analyses were performed:

1. Omnibus effect across all imaging modalities and groups
2. Main effect in violent offender groups
3. Main effect in violent patient groups
4. Main effect in healthy controls only

The dataset contained studies from multiple modalities. Accordingly, we also performed validation analyses where we assessed the main effect of aggression separately for each imaging modality (fMRI, structural MRI, PET and SPECT) across all subject groups. Because the fMRI studies were run with variable paradigms, we also analysed the fMRI studies separately for aggression-related tasks (such as Taylor’s Aggression Paradigm or videos of violence), other emotion-related tasks (such as emotion identification or emotion inducing pictures), executive function tasks (such as the Stroop task or a Go/No-Go task) as well as all other tasks. Finally, because the structural MRI dataset contained a substantial number of studies on different patient groups, we also ran separate ALE analysis for the structural MRI studies comparing primarily psychotic patients, antisocial patients, and all other patients against controls.

Concordance between the results across analyses was assessed by calculating spatial Spearman correlation between the ALE maps resulting from each analysis. To assess functional coactivation, correlations between the analyses and neural networks associated with 12 common brain functions (emotions, regulation, reward, memory, social, learning, pain, executive, inhibition, language, attention, motor) on Neurosynth (Yarkoni et al., 2011) were calculated with the Neurovault cognitive decoder tool (Gorgolewski et al., 2015). The results were visualized with the software *Surf Ice* (https://www.nitrc.org/projects/surfice/) and *MRIcroGL* (https://www.nitrc.org/projects/mricrogl/) (Rorden & Brett, 2000). To aid in visualization, regional (ROI-level) results were extracted from the activation maps for 14 regions involved in emotion, inhibition and aggression. These ROIs included the amygdala, pallidum, hippocampus, putamen, nucleus accumbens, caudate, insula, thalamus, anterior, middle and posterior cingulum cortex, orbitofrontal cortex and Broca’s areas 44 and 45 as defined per the AAL3 atlas. The mean z-scores were transformed to Pearson’s r and visualized as a heatmap with the R Superheat package (Barter & Yu, 2018; R Core Team, 2021).

## Results

### Included studies

A total of 406 studies and 28 968 subjects were included in the meta-analysis. 229 of the studies were fMRI, 124 structural MRI and the rest 58 PET or SPECT studies. Most (115) of the fMRI studies dealt with tasks focusing on emotions, 39 involved an aggression task, 43 an executive task and 33 involved a task not belonging in any of the categories. 186 of the studies included patients, 123 only healthy subjects and 106 had forensic samples. Altogether 10 934 participants were in patient group studies, 8029 participants in studies focusing on offenders and 10 266 participants in studies done only on healthy control subjects. Most (53) of the structural MRI-studies with patient groups had antisocial subjects (diagnoses included e.g. ASPD for adults and conduct disorder for children). Structural MRI studies with psychotic, mostly schizophrenic subjects were in the minority with 16 studies. There were 22 structural MRI patient group studies with other than psychotic or primarily antisocial patients. Diagnoses in this category varied and included e.g. autism spectrum disorders, substance use disorders, dementia, and paraphilic disorders.

### Omnibus analysis

The omnibus analysis across all studies (**Figure 2A**) revealed that aggression was consistently associated with alterations in amygdala, hippocampus, ventral (nAcc) and dorsal (caudate and putamen) striatum, anterior cingulate cortex, and orbital and dorsolateral prefrontal cortices as well as medial temporal cortex. Similar pattern was observed when the forensic subjects, patients, and healthy controls were analysed separately (**Figure 2B-D**), although insular effects were less prominent in the forensic population and healthy populations. Overlap between the omnibus analysis and the primary analyses (offenders vs. controls, patients vs. controls and healthy volunteers) is shown in **Figure S1-A**.

**Figure 2.**
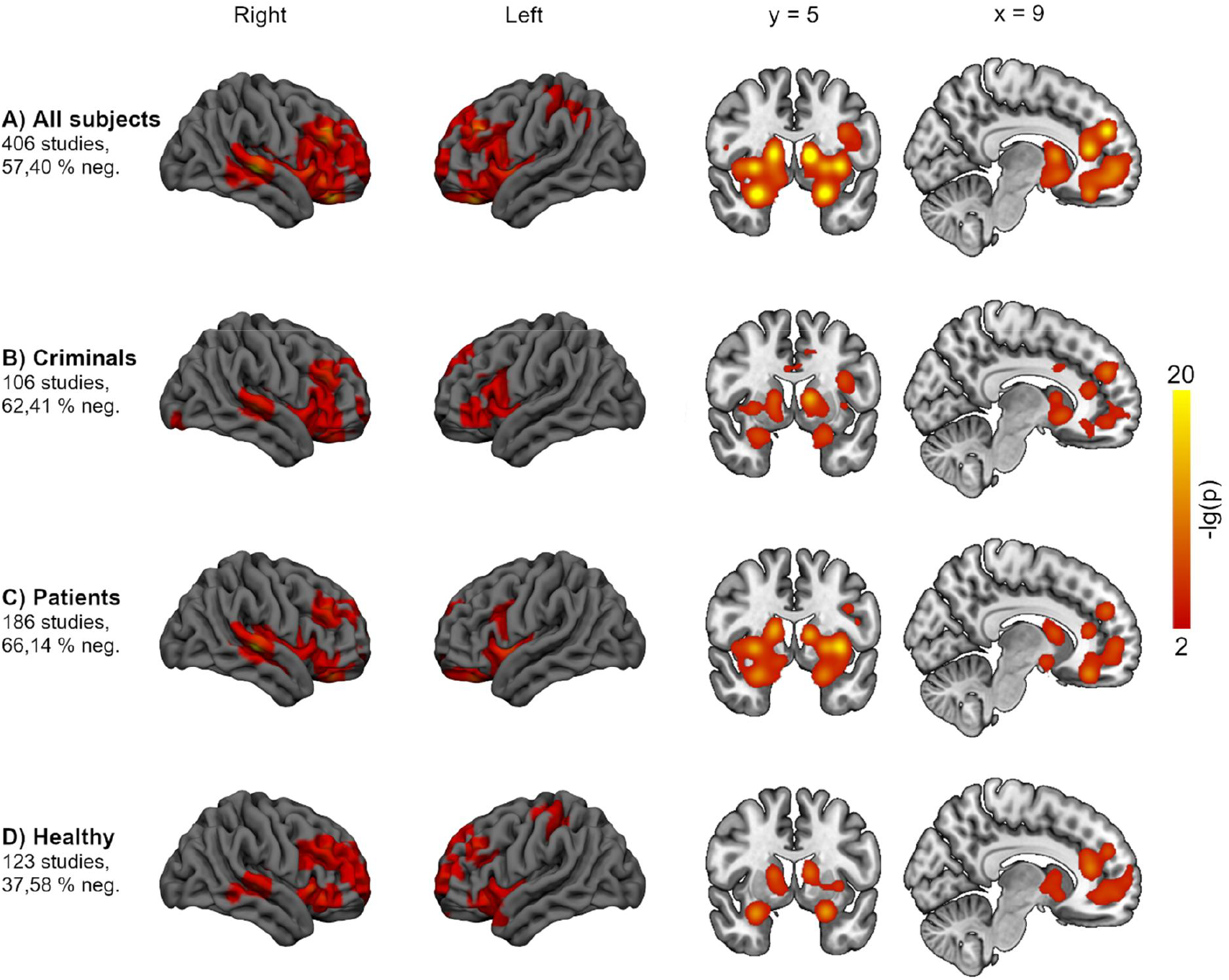
Results from the omnibus analysis (A), and subanalyses of offender group studies (B), psychiatric patient group studies (C), and healthy control group studies only (D). Beside each subanalysis is the percentage of negative effects among those included in the analysis. These subanalyses were highly similar with the omnibus analysis (spatial Spearman correlation coefficients between these subanalyses and the omnibus analysis were in the range of 0.8-0.9).

### Modality-specific analyses

Next, we analysed the effect of aggression separately for different imaging modalities (while pooling the subject groups) to test how consistent the effects were across structural, functional, and molecular studies. The effects were in general agreement with the omnibus analyses, but some subtle differences were observed. Results from the fMRI studies (**Figure 3A**) accorded with the omnibus analysis with significant effects in amygdala, hippocampus, ventral and dorsal striatum, anterior cingulate cortex, and orbital and lateral frontal cortices as well as medial temporal cortex; additional clusters were observed in the temporoparietal and inferior temporal cortices. Structural MRI (**Figure 3B**) studies yielded otherwise similar results, but the temporoparietal effects were not significant. The pattern of the molecular imaging studies (PET and SPECT) was also broadly similar (**Figure 3C**), with most notable differences being the absence of temporal and parietal effects and the large cluster in thalamus. Overlap between the omnibus analysis and the modality-specific analyses (fMRI, structural MRI, PET / SPECT is shown in **Figure S1-B**.

**Figure 3.**
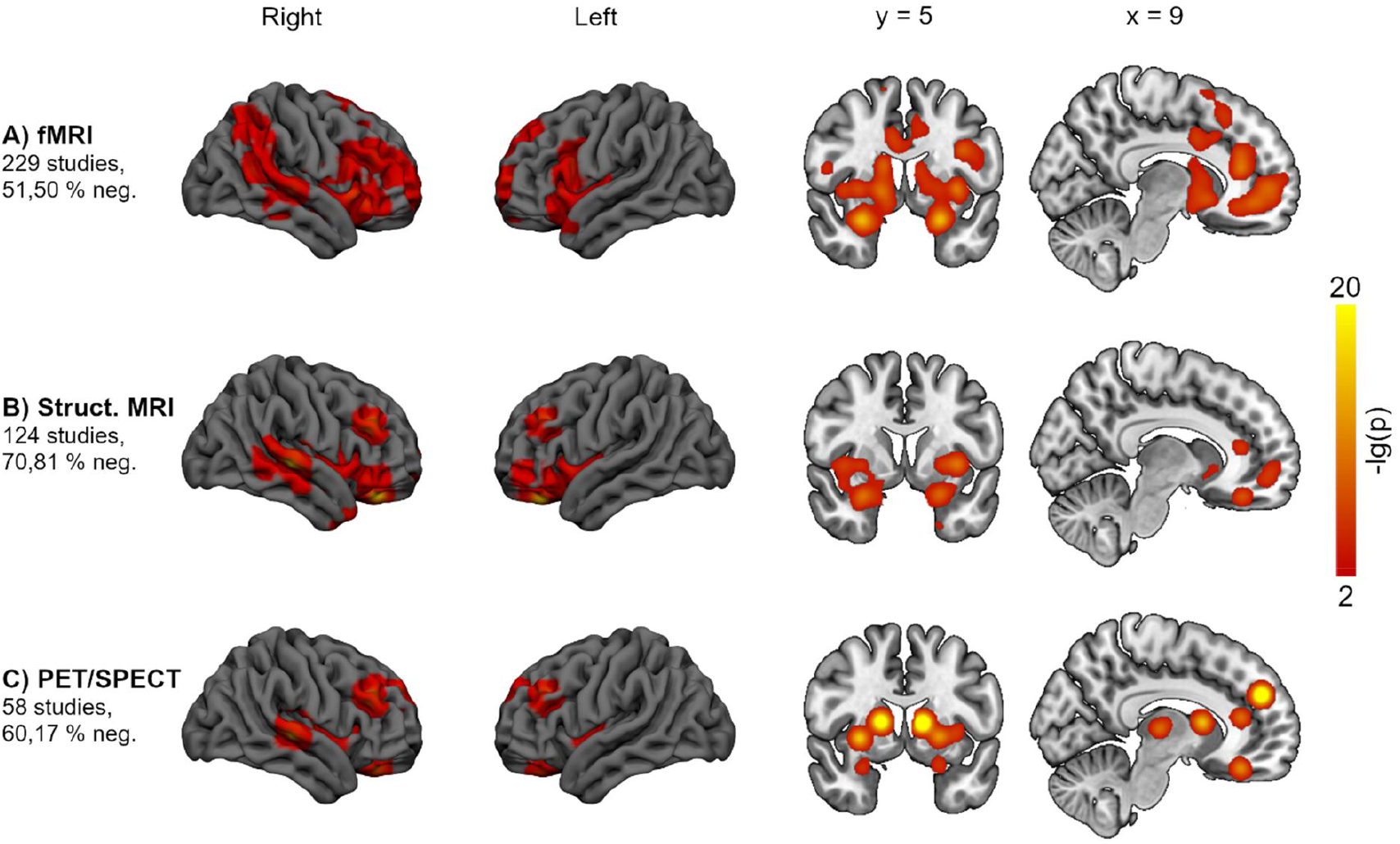
Modality-specific analyses. Results for the subsample of fMRI (A), structural MRI (B) and PET and SPECT (C) studies.

### Subanalyses for fMRI tasks

Next, we analysed the fMRI studies separately across the tasks. The aggression-related tasks yielded activations in inferior parietal and posterior temporal cortices, ACC, and pallidum. Tasks using other emotion-related protocols yielded an activation pattern that was more similar to the omnibus analysis, in that significant effects were found in amygdala, hippocampus, dorsal and ventral striatum, anterior cingulate, and lateral frontal cortices. For tasks tapping executive functions, the effects were localized in ACC and for the heterogeneous category of “other” protocols effects were observed primarily in medial frontal cortex and insula (**Figure S2**). Due to the heterogeneity of the tasks, no single region was found to be significantly associated with responses across all fMRI tasks.

### Subanalyses for patient groups in structural MRI studies

Because there were numerous structural MRI studies conducted in different aggressive patient groups (a total of 7318 subjects), we had sufficient statistical power for computing ALE separately for patients with antisocial disorders (n = 3181), psychotic disorders (n = 786) as well as other disorders pooled together (n = 3351). For the antisocial group, significant effects were found in amygdala, hippocampus, caudate, insula, and lateral prefrontal and orbitofrontal cortices. For psychotic patients the effects were focused on hippocampus, amygdala, caudate, and OFC. Finally, the heterogeneous category “other disorders” yielded effects in hippocampus, amygdala, and posterior superior temporal cortex (**Figure S3**).

### Concordance analyses and region-of-interest level results

Next, we computed the spatial similarity (Spearman correlation) of the ALE maps for the different analysis. This (**Figure 4**) revealed two broad clusters of studies. First, effects for the main omnibus analysis, fMRI studies (and specifically with emotional tasks), violent patients, healthy participants, offenders, and structural MRI studies were similar (r:s > 0.73). Second, results for structural MRI studies in antisocial, psychotic as well as other patient populations, PET and SPECT studies, and fMRI studies with aggression, executive, and mixed tasks differed more from the studies in the first cluster as well as from each other. Nevertheless, even the patterns of results from these studies were similar to the omnibus effect (r:s > 0.52) and with the other main study groups (r:s > 0.40).

**Figure 4.**
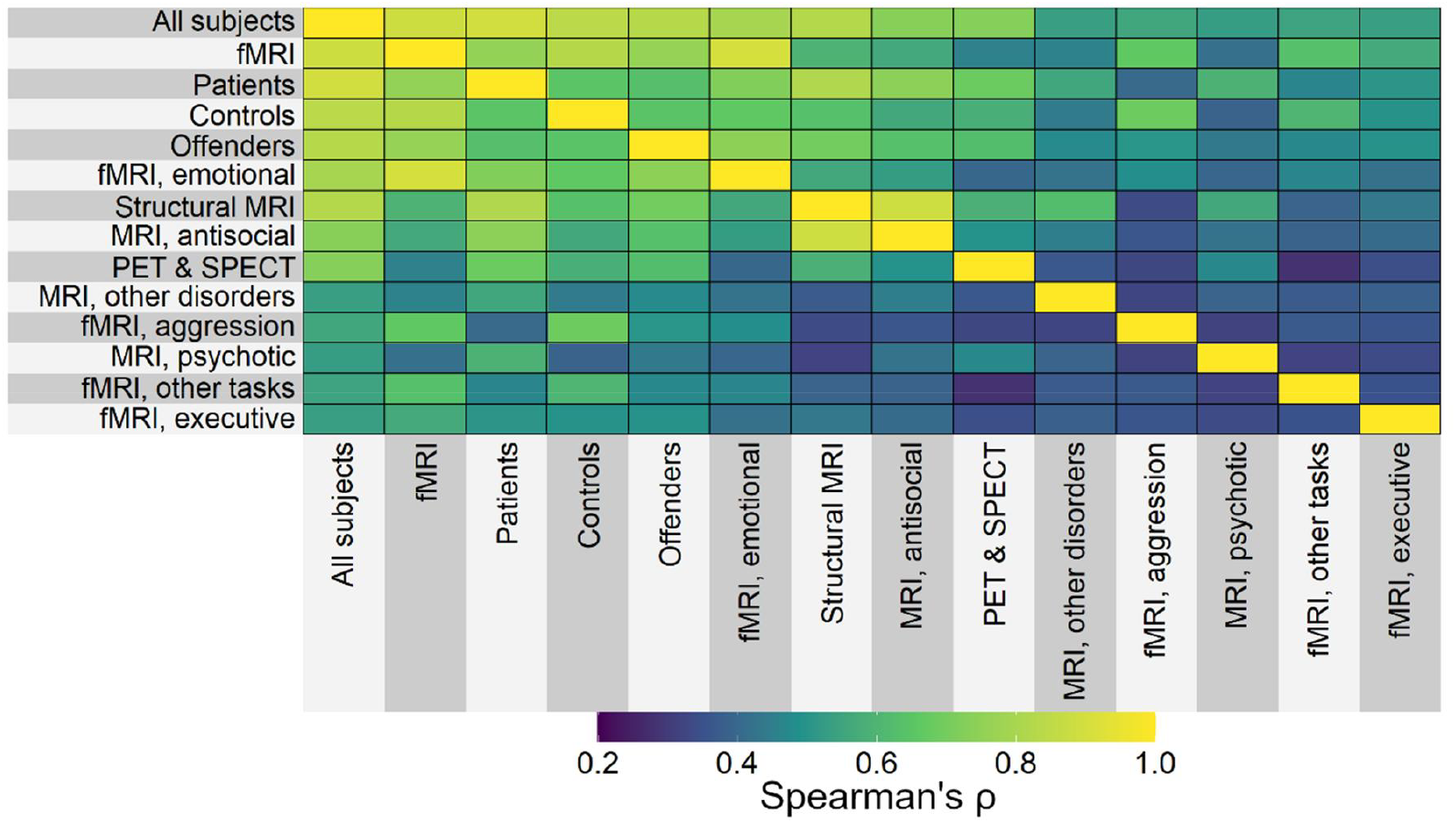
Similarity (spatial Spearman correlation) of the ALE results across analyses.

To summarize the regional results across the different analysis, we plotted the regional Z values (transformed into Pearson’s r) from the ALE maps for each ROI (**Figure 5**). This revealed that effects in amygdala, hippocampus, and striatum (particularly nACC and pallidum) were most consistently implicated in aggression across the analyses. Effects in anterior cingulum were also consistent but less strongly so, while effects were not in general found in BA44, OFC, thalamus, and middle and posterior cingulate cortices. The most notable exceptions to this pattern were the less intense hippocampal and ventral striatal activations in the fMRI studies as well as that consistent thalamic effects were only observed in the molecular imaging studies.

**Figure 5.**
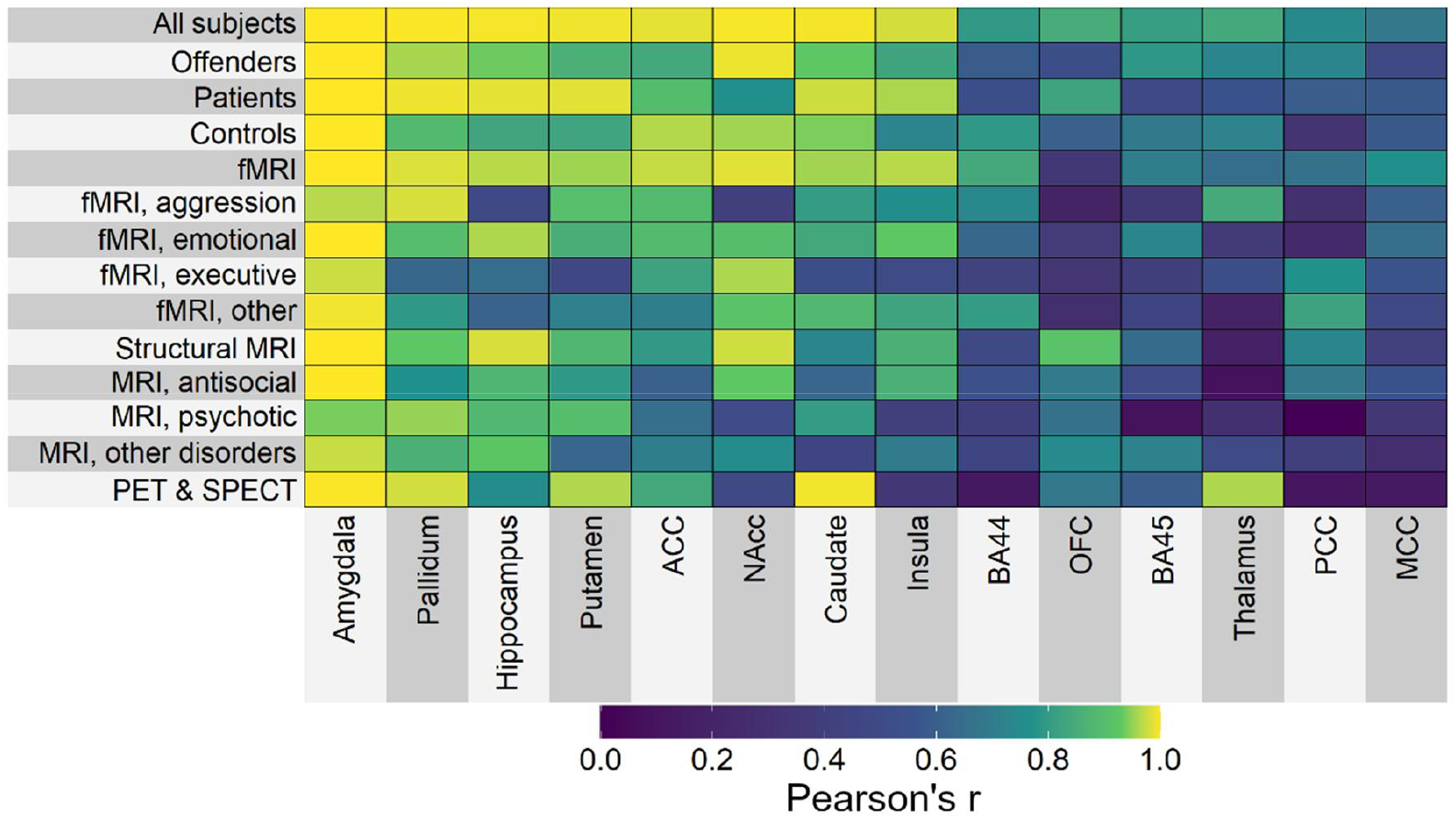
Region-of interest analysis. Note: the data are shown for visualization purposes only, statistical inference is based on the ALE maps.

### Functional coactivation analysis

To test which functional brain networks are most consistently associated with aggression, we ran a functional coactivation analyses by computing the spatial correlations between the meta-analytic ALE maps generated in this study and functional meta-analytic maps retrieved from NeuroSynth database for different cognitive and emotional functions (Yarkoni et al., 2011). The NeuroSynth maps were retrieved with keywords emotions, regulation, reward, memory, social, learning, pain, executive, inhibition, language, attention, and motor. This analysis (**Figure 6**) revealed that functionally the aggression-related effects in the ALE maps were most consistently associated with emotion, regulation, and reward related functions. Moderate associations were observed with memory, social functions, learning and pain, while weakest associations were found for executive, inhibitory and language and motor functions. Importantly, this pattern of functional coactivations was consistent across the results from the different ALE analyses.

**Figure 6.**
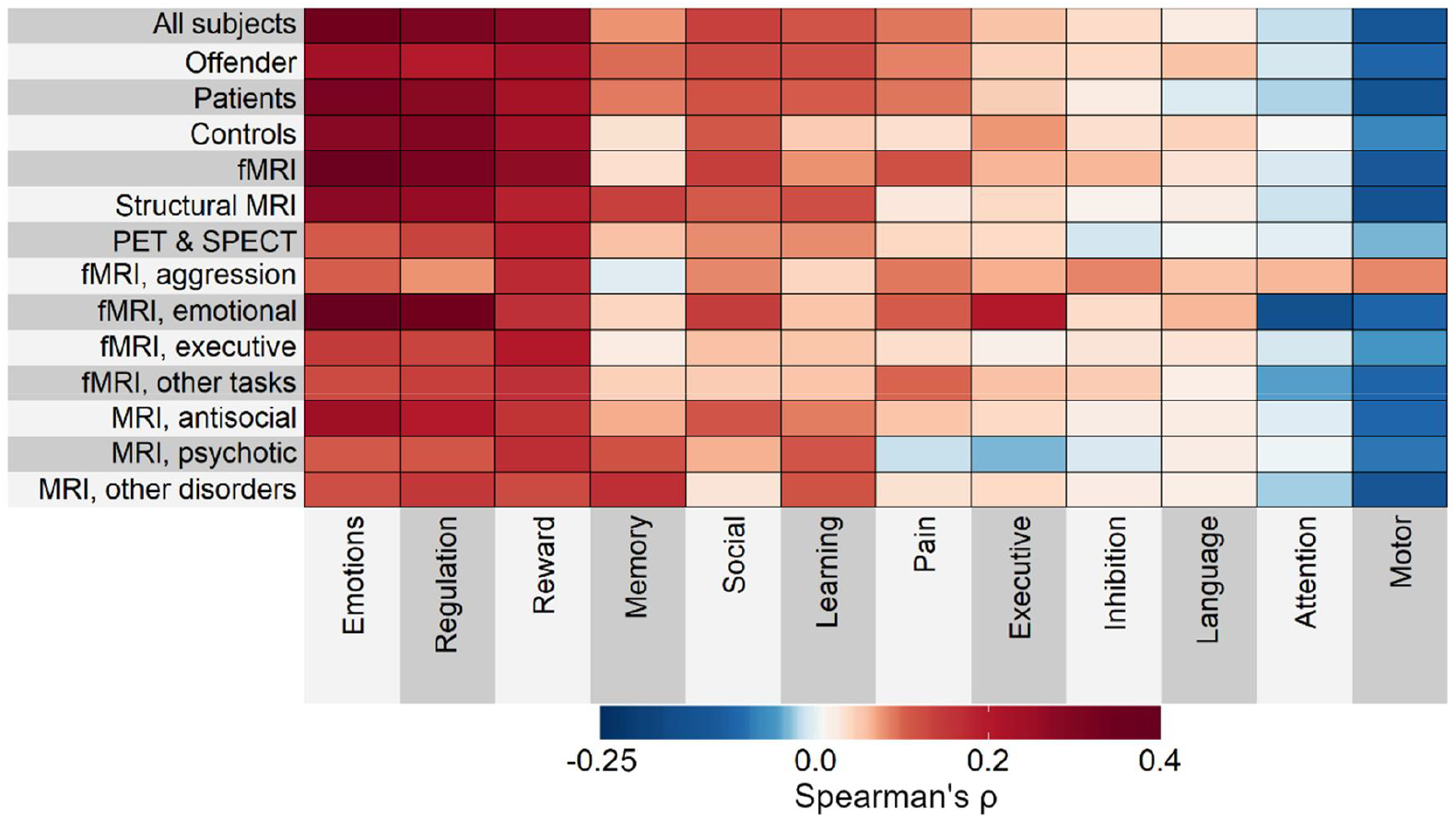
Functional coactivation analysis for task-specific brain networks across analyses.

Finally, we conducted the coactivation analysis at regional level to address the functional significance of the regions associated with aggression in the ALE. Clustering of the omnibus ALE map identified five main clusters in the data. The striatolimbic cluster comprised of amygdala, hippocampus, thalamus, dorsal and ventral striatum, anterior insula and lateral inferior frontal cortices. The medial frontal cluster comprised of anterior cingulate cortex, medial prefrontal and orbitofrontal cortices. The dorsolateral prefrontal cluster contained bilateral foci in the middle frontal gyrus. The temporal cluster was focused on the posterior and middle parts of superior and middle temporal cortex, and the parietal cluster comprised S1 in the postcentral gyrus extending also to inferior parietal cortex. We then computed the proportional overlap between each cluster (C) and functional networks (F) described above as N_(C∩F)_/N_c_ thus the proportions reflect the proportion of each cluster belonging to each functional network (**Figure 7**). The striatolimbic cluster had over 40% overlap with all the tested networks except language network, while most salient association were found with reward (74%), learning (63%), memory (60%), social (55%), and regulatory (52%) functions. The medial frontal cluster was associated with primarily with reward (44%) and social (42%) functions, with all other functions showing < 25% overlap. The dorsolateral cluster in turn showed broad associations memory (79%), executive (77%), attentional (65%), inhibition (55%) and motor (54%) functions. The parietal cluster was highly overlapping with motor (99%) and attentional (82%) functions but also had large overlap with memory (76%), learning (75%) and inhibitory (68%) functions. Finally, the temporal cluster had large overlap with language functions (92%) and more modest overlap with attentional (67%), motor (60%) and social (38%) functions.

**Figure 7.**
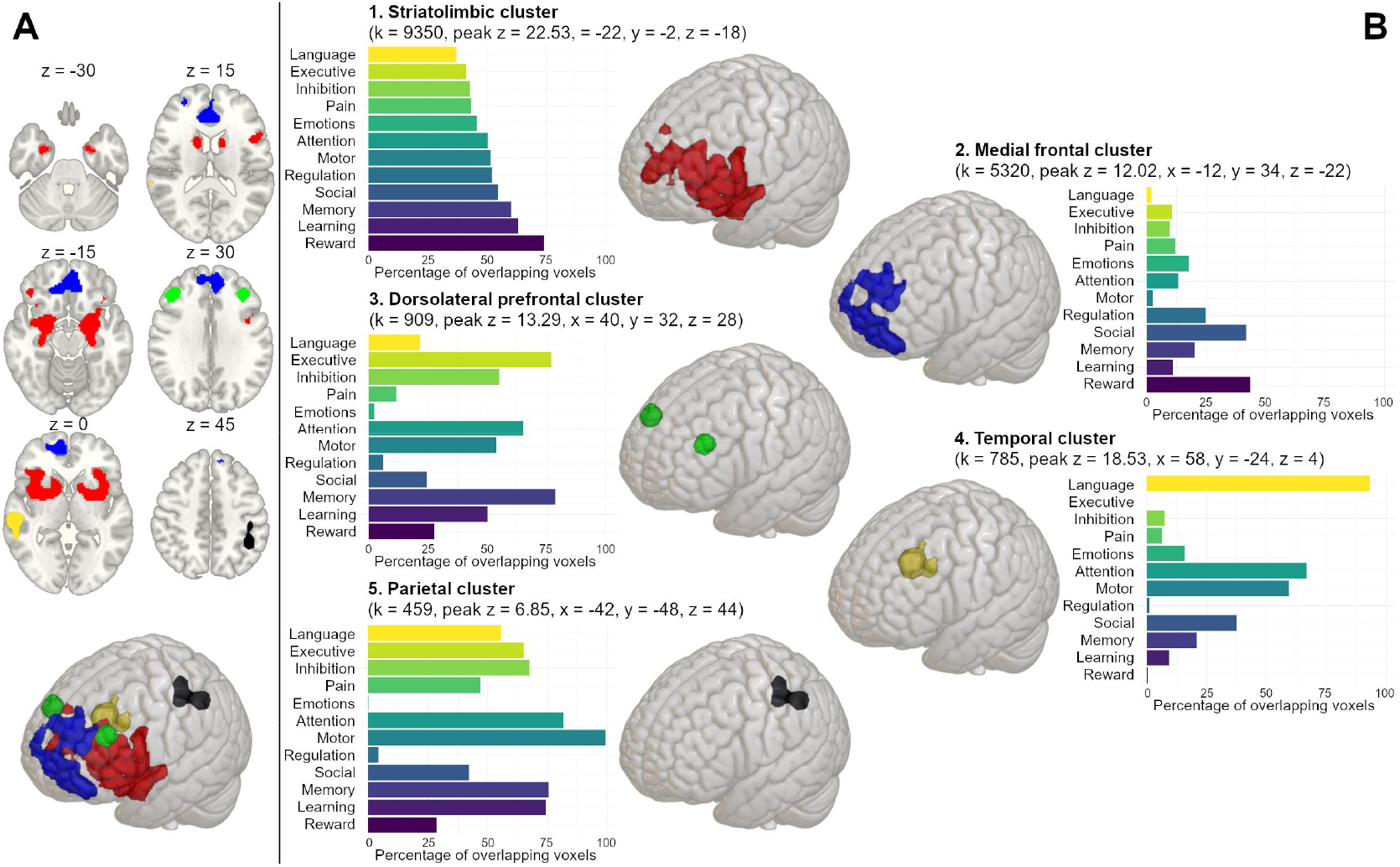
Regional functional coactivation analysis (B) for the clusters (A) observed in the omnibus analysis. The bar charts show the proportion (%) of overlap between each cluster and each functional meta-analytic map retrieved from NeuroSynth. Above each bar chart is presented the size of the cluster, peak z value of the cluster, and the MNI coordinates of the peak.

## Discussion

Our main finding was that the structure, function, and molecular organisation of a common set of limbic, paralimbic, frontal, and temporoparietal brain regions is consistently implicated in aggression in forensic, clinical, and healthy populations. Most reliable effects were found in the amygdala, hippocampus, striatum, and anterior cingulate and lateral prefrontal cortices. These results were observed consistently, independent of whether cerebral macrostructure, molecular organization, or functional profiles were considered. Functional coactivation analyses revealed that aggression was particularly associated with brain networks subserving reward and emotion processing as well as their regulation. Alterations in the structure and functioning of this general aggression network (**GAN**) thus underlies all studied types of aggression and thus the neural basis of aggression in clinical and forensic groups is not distinctly different from healthy populations. All in all, our findings reveal a unified neural network underlying the continuum from benign to pathological aggression and underscore the role of aberrant functioning of reward, emotion, and self-regulation circuits in aggressive behaviour.

### Consistent aggression network across imaging modalities and groups

Aggression is a complex and heterogeneous construct (Rosell & Siever, 2015). Although we used a broad definition of aggression encompassing reactive and proactive aggression, pathological and criminal violence as well benign acute laboratory-induced aggression, the same aggression network emerged from the ALE analyses despite the heterogeneity of study designs and populations. We analysed the data from structural, functional and molecular studies both separately and independently, and in found that the results were concordant across imaging modalities. The only exception was the subanalysis with PET and SPECT studies, whose results were more narrowly focused on striatum and thalamus. This is not unexpected, as PET and SPECT radiotracer binding can be regionally selective, and for example most dopaminergic receptor tracers bind selectively to striatothalamic areas (Malén et al., 2022, 2024). The overall concordance likely reflects multiple underlying biological mechanisms. Specific neurotransmitter and neurotransporter levels are reflected in regional grey matter densities derived from MR images. The grey matter density signal derived from MR images for VBM analysis reflects gross density of neurons, as a large bulk of cell bodies and neuropils are in grey matter (Purves et al., 2017). Consequently, binding of PET receptor and transporter radioligands is associated with the underlying mesoscopic differences in grey matter densities across subjects, as measured with VBM (Kraus et al., 2012; Manninen et al., 2021; Woodward et al., 2009). Similarly fusion imaging studies with PET and fMRI have shown the availability of regional neuroreceptors are linked with regional neural responses to e.g. affective stimulation in fMRI (Karjalainen et al., 2017, 2019). Although brain structure and function are significantly correlated, the correspondence is not perfect because as function measured by fMRI reflects complex multisynaptic interactions across large-scale structural brain networks (Suárez et al., 2020; Sui et al., 2014). Nevertheless, our data suggest that the same brain network whose structural and molecular organization is altered in aggressive individuals also coordinates transient responses during aggressive episodes as demonstrated by the fMRI data.

In addition to consistency across modalities, the GAN was also consistently observed in the separate analysis focusing on pathological aggression in psychiatric patients, criminal aggression in incarcerated populations as well as aggression in healthy controls. These findings underline that that a common aggression network is involved in both everyday benign aggression as well as pathological or criminal variants of aggressive behaviour and confirm that pathological and criminal aggression are manifestations of the aberrant structure and function in the same system that governs aggressive responses in typical healthy individuals. However, the subanalyses across different aggressive patient populations revealed some differences. Results from ASPD corresponded well with the results from the omnibus analysis and were focused on the GAN, highlighting alterations in the amygdala, striatum and orbitofrontal cortex previously implicated in ASPD (Tully et al., 2023) but also in temporocortical regions involved in social perception (Santavirta et al., 2023). This accords with the behavioural phenotype of ASPD which is characterised by both aggression as well as deviance from social norms and disregarding others’ well-being. The ALE map from ASPD also accorded with that from the violent offenders, likely reflecting the fact that antisocial personality disorder and concomitant alterations in brain structure and function constitute a risk factor for criminal offending as almost one third of incarcerated offenders have diagnosed ASPD (Black et al., 2010; Fridell et al., 2008). In turn, for patients with psychosis, the results were restricted primarily to striatum and parietal cortices. Although the sample size for psychotic patients was limited (16 studies), these results suggest that aggressive outbursts in psychosis stem from different neural perturbations than those in ASPD. Indeed, studies have found that violent criminality in psychosis is driven by external trigger such as stressful life events, injuries or substance abuse, rather than interpersonal conflicts or strategical acquisition of resources through violence (Sariaslan et al., 2016).

### Emotional, motivational, and regulatory components of the general aggression network

Regionally, aggression was most consistently associated with structure and function in the amygdala, hippocampus, ventral and dorsal striatum, and anterior cingulate cortex (**Fig 5**). Functional coactivation analysis (**Fig 6-7**) confirmed that this constellation of regions shows largest overlap with emotional, regulatory and reward-related processes. Amygdala has consistently been linked with aggression in previous studies and is involved in processing of emotional information and the identification of threats in one’s environment (Sergerie et al., 2008; Šimić et al., 2021). Heightened amygdala activity may reflect hypersensitivity to perceived threats which can lead to hostile attribution bias, a common finding in most aggressive samples. The amygdala is well connected with the hippocampus, and we also observed consistent aggression-dependent functional, structural, and molecular changes in this area. Hippocampus is critical for memory and emotional learning (Bird & Burgess, 2008; Lacagnina et al., 2019) and it also contributes to fear extinction – learning which previously threatening stimuli no longer predict threat (Shin & Liberzon, 2010). Thus, hippocampal changes could lead to dysfunction learning from threats, which in turn could lead to increased aggression. Striatal areas, especially the pallidum, have been consistently linked with motivational processes (e.g. Smith et al., 2009).

Striatum in turn integrates widespread cortical inputs and modulates thalamocortical activity indirectly. Particularly the ventral and dorsomedial striata have been implicated in aggression due to their role in response selection as well as in governing motor and emotional response sequences (Rosell & Siever, 2015). Striatal abnormalities could thus lead to dysregulated reward-seeking behaviour, where aggressive actions are orchestrated by heightened sensitivity to rewards (material gain, social dominance) or reduced sensitivity to punishment cues. Regions associated with emotional regulation, namely the anterior cingulate and orbitofrontal cortex, were also consistently implicated in aggression. Anterior cingulate cortex is well known for its role in affective (ventral part) and cognitive (dorsal part) control. The affective subdivision is connected to amygdala, periaqueductal grey, nucleus accumbens, hypothalamus, anterior insula, hippocampus, and orbitofrontal cortex while the cognitive subdivision connects with lateral prefrontal cortex, parietal cortex, and premotor and supplementary motor cortices (Bush et al., 2000). Although our results highlight the role of both affective and cognitive subdivisions in aggression, it is notable that majority of the other regions highlighted by the various ALE analyses are known to connect with the affective rather than the cognitive subdivision.

While the GAN showed consistent overlap with meta-analytic functional activation patters for emotions, regulation and reward, the associations with inhibitory, executive, attentional as well as langue networks were weaker and correlations with attention and motor networks were negative. The prominence of affective processes in aggression was further supported by the analysis for different tasks in the fMRI studies. Tasks involving direct proxies of aggressive behaviour and those involving emotional stimuli consistently activated the same cortical-subcortical aggression network as was observed in the omnibus analysis, suggesting a central role of emotional perturbances in aggressive behaviour (Davidson et al., 2000). In turn, tasks tapping executive functions only yielded small frontotemporal and cingulate clusters (similarly as the emotion and aggression tasks), while effects in the central executive brain network (Niendam et al., 2012) whose perturbations are also involved in various psychiatric conditions (Goodkind et al., 2015) were not salient. The heterogeneous category of fMRI tasks only yielded limited number of ALE clusters primarily in the temporal cortex and striatum. However, these results reflect only the overall functional overlap of the GAN and when the meta-analytic map was parcellated into five main clusters, we found that the dorsolateral prefrontal cluster and parietal clusters showed consistent overlap with inhibitory, attentional and (working) memory functions. Altogether the data show that aggression is most consistently associated with large-scale alterations in the stritolimbic reward, emotion and emotion regulation networks, aberrant structure and function of the dorsolateral prefrontal and parietal regions that subserve inhibition and working memory related functions.

Our findings stress the importance of emotion and its regulation to aggression accords with previous studies and theoretical models of aggression. The general aggression model (Allen et al., 2018) states that the present internal sate (affect, cognition and arousal) of an individual affects their appraisal of their environment and context and their decision-making processes, which in turn affects their behaviour and thus aggression. The chosen behaviour changes the current environment, which in turn affects the present internal state of the individual. Our finding of the centrality of emotion and emotion regulation networks in aggression fit well with this model, since with disrupted emotional regulation the affectional state of an individual can spiral a mildly irritating or stressful situation into full-blown aggressiveness. Amygdala and ACC function might explain the early emotional responses to the situation while late cognitive appraisals and decision whether to act aggressively could involve PFC and OFC. General aggression model and the importance of emotion regulation also link with the developmental track of aggressiveness, since aggression in youth is related to the aberrant maturation of the PFC (Hostetler et al., 2024). This can lead to harmful choices of behaviour, leading to further escalation and hostility of the environment and problems with social interactions. This with time can reduce the possibility to learn emotion skills such as emotion regulation and processing of emotional information of the environment in relation to peers. Thus, the relationship between neural changes and aggression can be moderated by the social context, as seen in how the relationship between amygdala-ventrolateral PFC connectivity and aggression is moderated by social impairment in children (Ibrahim et al., 2022). Furthermore, as repeated engagement with aggressive behaviour can reinforce an aggressive model of behaviour leading to neural changes, it becomes clear how large of an impact disrupted emotion regulation can have on the development of aggression. This suggests differences in the way aggressive individuals assess and react to their emotional environments, leading to violence, rather than being unable of inhibition. This is interesting when considering how consistently attention-deficit disorders have been linked with aggression and criminal behaviour (Mohr-Jensen & Steinhausen, 2016). All in all, our results thus underscore the role of excessive bottom-up drive rather than lacking top-down control in aggression.

### Limitations

ALE does not allow differentiating between positive and negative effects in the original studies, which complicates the interpretation of our results and might also dismiss important information on the nature of changes found in the reported areas. We nevertheless addressed this by reporting the percentage of negative and positive effects included in each analysis, although determining the direction of the relative signal in fMRI studies can be artificial and even subjective due to the way the subtraction contrasts are set up. We however stress that the goal of this study was to map cerebral areas involved in aggression independently of the direction of the association. The strength of PET and SPECT studies included in this meta-analysis is their high molecular specificity allowing in vivo quantification of specific tissues with different radioligands. However, due to limited statistical power the analysis could not be divided into subanalyses between radioligands to capitalize in the uniqueness of these imaging modalities. Converting reported regional results into coordinate space using the centre of mass of atlas ROIs involves uncertainties, but similar uncertainty is also involved in the cluster peak-based analysis because the shape of the clusters is not accounted for in the analysis. Finally, this meta-analysis of cross-sectional data cannot address the exact role of genetic causes and experience-dependent plasticity in the observed alterations in the composition of the general aggression network in the clinical and forensic samples, which needs to be addressed in future longitudinal studies.

## Conclusions

Our large-scale multimodal meta-analysis of the brain basis of aggression revealed a common aggression network spanning multiple neurocognitive systems involved particularly in emotional and reward processing, memory and learning, and emotional regulation. Contribution of the general executive networks to aggression were limited. The found common aggression network was consistently associated with the whole continuum of aggression from benign to criminal and pathological. Further clinical studies are needed to assess the development of aggression and especially the prevention and treatment of it.

## Acknowlegements

This study was supported by the Academy of Finland (grants #294897 and #332225), Sigrid Juselius Stiftelse, Signe och Ane Gyllenbergs Stiftelse, and the TYKS foundation.

