## Supplementary information for "Shared brain basis of aggression in clinical, forensic, and healthy samples: A meta-analysis"

Harri Harju[1], Jouni Tuisku[1] and Lauri Nummenmaa[1,2,3]

1 Turku PET Centre
2 Department of Psychology, University of Turku
3 Turku University Hospital, University of Turku

Address Correspondence to:
Harri Harju
Turku PET Centre
Kiinamyllynkatu 4-6
FI-20540 Turku, Finland


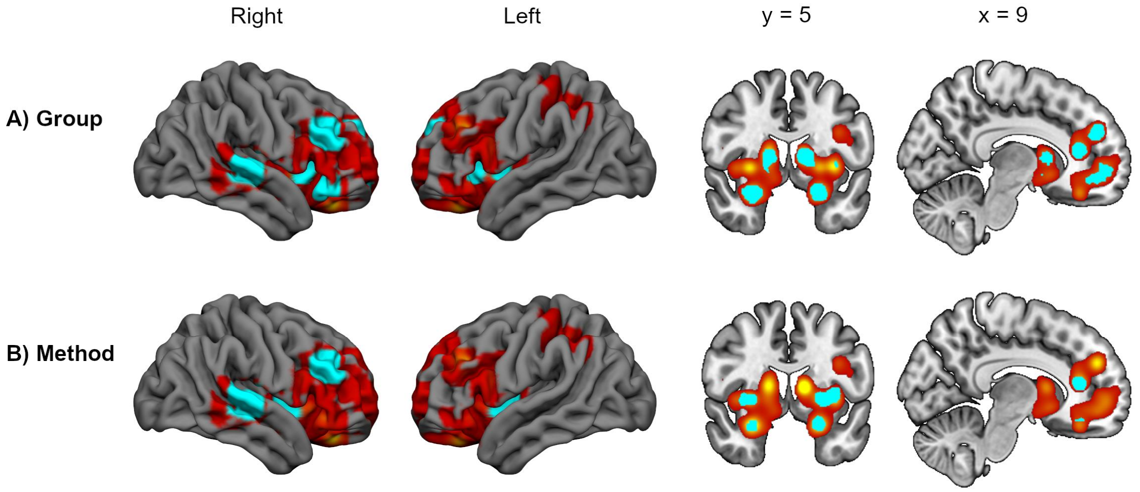


Figure S1. Overlapping voxels (shown in cyan) between the omnibus analysis and all the primary analyses (offenders vs. controls, patients vs. controls and healthy volunteers) (A) and between the omnibus analysis and all the modality-specific analyses (fMRI, structural MRI, PET / SPECT) (B). The overlapping voxels are shown on top of the omnibus analysis (red-yellow).

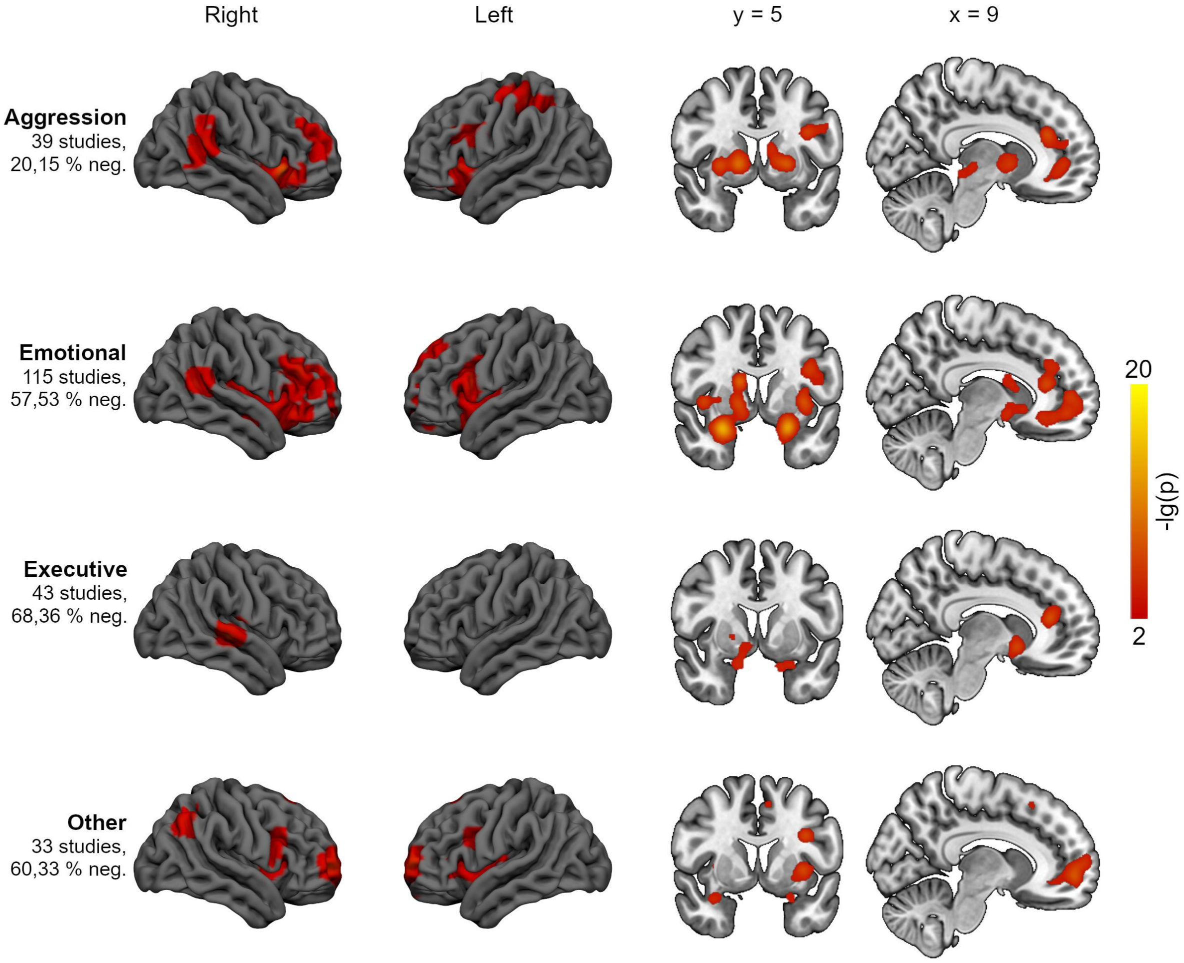


Figure S2. Results from the subanalyses for each fMRI task type. Aggression-related tasks (n = 1556) included e.g. Taylor's Aggression Paradigm and videos of violence, other emotion-related tasks (n = 6377) included e.g. emotion identification tasks or emotion inducing pictures, and executive function tasks (n = 4040) included e.g. Stroop tasks and Go/No-Go tasks. All other tasks (n = 1668), such as gambling tasks and neutral pictures, were assigned to the heterogeneous other-category. Beside each subanalysis is the percentage of negative effects among those included in the analysis.

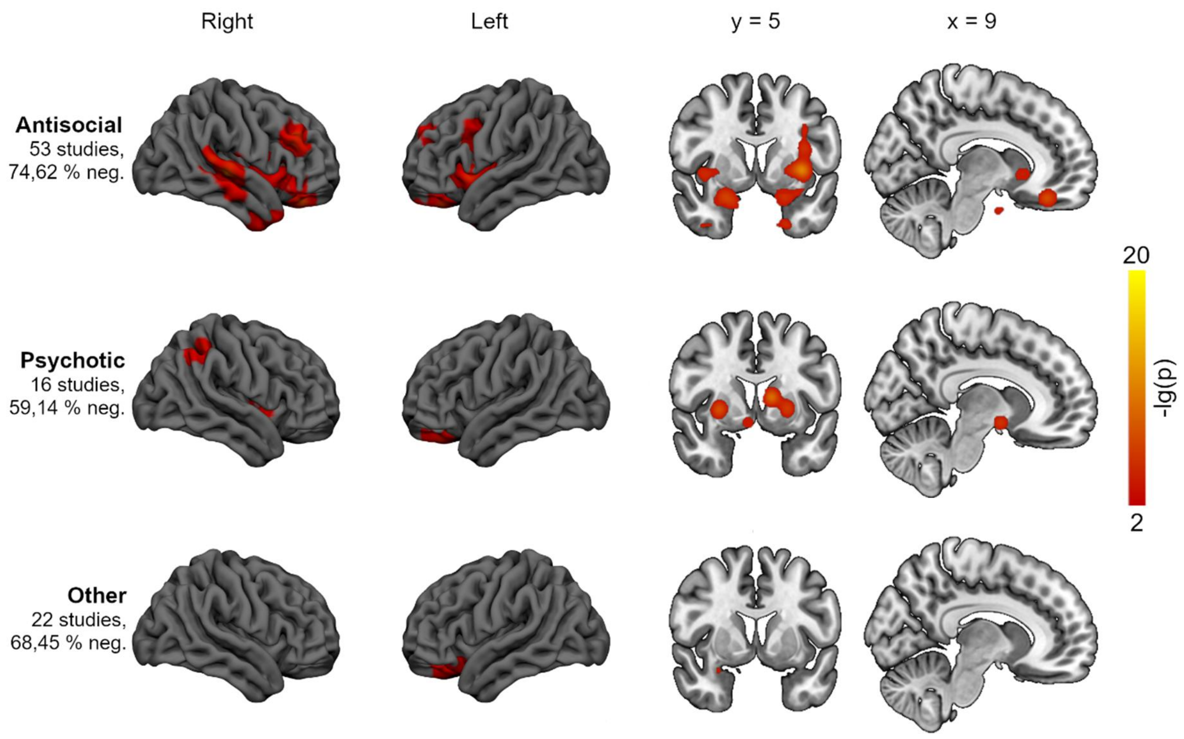


Figure S3. Results from the subanalyses for patient groups in structural MRI studies. Primarily antisocial patients (n = 3181) had diagnoses such as antisocial personality disorder or conduct disorder, and psychotic patients (n = 786) were mostly diagnosed with schizophrenia. The heterogeneous category of "other patients" (n = 3351) included diagnoses such as autism spectrum disorders, substance use disorders, dementia, and paraphilic disorders. Beside each subanalysis is the percentage of negative effects among those included in the analysis.

### Table S1. Included studies

| Num. | Study | Modality | Modality detail (fMRI: task/stimulus, MRI: method, PET/SPECT: tracer) | Design | Measure of antisociality | Patient group | n | Men | Women | Relevant contrasts |
|---|---|---|---|---|---|---|---|---|---|---|
| 1 | Abe et al. (2018) | fMRI | Coin-flip prediction task | Correlational | PCL-R | Dishonest incarcerated individuals | 43 | 43 | 0 | 1 |
| 2 | Achterberg et al. (2016) | fMRI | Social Network Aggression Task (SNAT) | Correlational | na | na | 30 | 15 | 15 | 1 |
| 3 | Achterberg et al. (2018) | fMRI | SNAT | Correlational | na | na | 385 | 181 | 204 | 1 |
| 4 | Achterberg et al. (2020) | fMRI | SNAT | Correlational | na | na | 360 | 173 | 187 | 1 |
| 5a | Aggensteiner et al. (2020) | fMRI | Emotional face matching task | Group comparison | CBCL, ICU | ODD and/or CD or individuals with CBCL T value of over 79 in aggression or rule-breaking behaviour | 177 | 129 | 48 | 2 |
| 5b | Aggensteiner et al. (2020) | fMRI | Emotional face matching task | Correlational | CBCL, ICU | ODD and/or CD or individuals with CBCL T value of over 79 in aggression or rule-breaking behaviour | 166 | 122 | 44 | 1 |
| 6 | Aghajani et al. (2021) | fMRI | Emotional face recognition/resonance task | Group comparison | na | Antisocial male juvenile offenders with CD | 81 | 81 | 0 | 3 |
| 7 | Aharoni et al. (2013) | fMRI | Go/NoGo | Correlational | na | Adult male offenders | 86 | 86 | 0 | 1 |
| 8 | Alia-Klein et al. (2008) | PET | [ 11C]clorgyline | Correlational | MPQ aggression scale | na | 27 | 27 | 0 | 1 |
| 9 | Alia-Klein et al. (2009) | fMRI | Reaction time and (sub)vocalization task focusing on words Yes and No | Correlational | revised State-Trait Anger Expression Inventory (STAXI-2) | na | 27 | 27 | 0 | 2 |
| 10 | Alia-Klein et al. (2014) | PET | [18F]fluoro-deoxyglucose | Group comparison | BPAQ Physical aggression scale | Aggressive males | 25 | 25 | 0 | 5 |
| 11 | Alvarenga et al. (2012) | MRI | VBM | Correlational | DY-BOCS | OCD | 38 | 15 | 23 | 2 |
| 12 | Amen et al. (2007) | SPECT | [99mTc]-d, I-HMPAO | Group comparison | na | Murderers | 22 | 22 | 0 | 1 |
| 13 | Amen et al. (1996) | SPECT | [99mTc]-d, I-HMPAO | Group comparison | na | Psychiatric patients with aggressive behaviour | 80 | 60 | 20 | 2 |
| 14 | Amen & Carmichael (1997) | SPECT | Ceretec (99m TC hexamethylpropylene amine oxime) | Group comparison | na | Children and adolescents with ODD | 84 | 63 | 21 | 1 |
| 15 | Anderson et al. (2018) | fMRI | Auditory oddball task | Correlational | PCL-R | Adult inmates | 168 | 168 | 0 | 3 |
| 16 | Anderson et al. (2017) | fMRI | Emotion attention task | Correlational | PCL-R | Adult male incarcerated offenders | 120 | 120 | 0 | 4 |
| 17 | Baird et al. (2010) | fMRI | Affect recognition from facial stimuli mixed with fixation trials | Correlational | BPAQ hostility scale | na | 14 | 0 | 14 | 1 |
| 18a | Barkataki et al. (2008) | fMRI | Visual go/nogo -task | Group comparison | na | ASPD and history of violence | 28 | 28 | 0 | 1 |
| 18b | Barkataki et al. (2008) | fMRI | Visual go/nogo -task | Group comparison | na | SCZ and a history of violence | 26 | 26 | 0 | 2 |
| 19a | Barkataki et al. (2006) | MRI | MEASURE program and the Cavalieri method | Group comparison | na | ASPD | 28 | 28 | 0 | 2 |
| 19b | Barkataki et al. (2006) | MRI | MEASURE program and the Cavalieri method | Group comparison | na | SCZ and history of violence | 28 | 28 | 0 | 2 |
| 19c | Barkataki et al. (2006) | MRI | MEASURE program and the Cavalieri method | Group comparison | na | SCZ and history of violence | 28 | 28 | 0 | 1 |
| 20 | Baskin-Sommers et al. (2012) | fMRI | Instructed fear-conditioning task | Group comparison | ESI | Externalizing inmates | 37 | 37 | 0 | 2 |
| 21a | Beames et al. (2020) | fMRI | Solving anagrams while being insulted/provocated | Correlational | BPAQ | na | 21 | 12 | 9 | 2 |
| 21b | Beames et al. (2020) | fMRI | Solving anagrams while being insulted/provocated | Correlational | BPAQ | na | 24 | 8 | 16 | 1 |
| 22 | Beaver et al. (2008) | fMRI | Pictures of emotional faces | Activation study | na | na | 22 | 9 | 13 | 2 |
| 23 | Beckwith et al. (2018) | MRI | VBM | Correlational | PPI | na | 155 | 65 | 90 | 1 |
| 24 | Beckwith et al. (2021) | MRI | VBM | Correlational | Number of criminal arrests (total and violent separately) | na | 123 | 52 | 71 | 1 |
| 25 | Benegal et al. (2007) | MRI | VBM | Correlational | SSAGA-II | Alcohol-naive offspring of early onset alcohol-dependent individuals with two or more dependent first-degree relatives | 41 | 41 | 0 | 1 |
| 26a | Bertsch et al. (2013) | MRI | VBM | Group comparison | PCL-factor 1 | Offenders with BPD | 27 | 27 | 0 | 1 |
| 26b | Bertsch et al. (2013) | MRI | VBM | Group comparison | PCL-factor 1 | Antisocial offenders | 26 | 26 | 0 | 1 |
| 26c | Bertsch et al. (2013) | MRI | VBM | Group comparison | PCL-factor 1 | Antisocial offenders | 25 | 25 | 0 | 2 |
| 27 | Besteher et al. (2017) | MRI | VBM | Correlational | SCL-90-R | na | 409 | 184 | 225 | 1 |
| 28a | Beyer et al. (2015) | fMRI | TAGG showing either neutral or angry video of opponent before decision phase | Activation study | na | na | 41 | 41 | 0 | 2 |
| 28b | Beyer et al. (2015) | fMRI | TAGG showing either neutral or angry video of opponent before decision phase | Correlational | TAGG punishment selections | na | 41 | 41 | 0 | 1 |
| 29 | Beyer et al. (2014) | fMRI | TAGG and viewing pictures of neutral/emotional, social/single pictures | Correlational | Taylor aggression paradigm punishment selections | na | 34 | 14 | 20 | 1 |
| 30 | Birbaumer et al. (2005) | fMRI | Aversive differential pavlovian delay conditioning with neutral faces serving as conditioned and pain as unconditioned stimuli | Group comparison | PCL-R | Emotionally detached psychopaths with criminal records | 20 | 20 | 0 | 2 |
| 31a | Biundo et al. (2015) | MRI | Freesurfer | Group comparison | MIDI, QUIP-RS | PD and ICDs | 91 | 51 | 40 | 2 |
| 31b | Biundo et al. (2015) | MRI | Freesurfer | Group comparison | MIDI, QUIP-RS | PD and ICDs | 110 | 70 | 40 | 3 |
| 32 | Bjork et al. (2010) | fMRI | MID | Correlational | CBCL externalizing scale | Adolescents with externalizing disorders | 24 | 18 | 6 | 1 |
| 33 | Bjork et al. (2012) | fMRI | Factorial Reward Anticipation task (modified MID) | Correlational | PPI | na | 31 | 18 | 13 | 2 |
| 34a | Bobes et al. (2013) | fMRI | Fearful and neutral faces presented to subjects. Structural with VBM | Group comparison | RPQ | Violent males | 54 | 54 | 0 | 1 |
| 34b | Bobes et al. (2013) | MRI | VBM | Group comparison | RPQ | Violent males | 54 | 54 | 0 | 1 |
| 35 | Boccardi et al. (2013) | MRI | Manual tracing and radial distance mapping | Group comparison | PCL-R | Violent offenders | 51 | 51 | 0 | 1 |
| 36 | Boccardi et al. (2011) | MRI | Cortical pattern matching and radial distance mapping | Group comparison | PCL-R | Violent offenders | 51 | 51 | 0 | 1 |
| 37 | Boccardi et al. (2010) | MRI | Manual tracing and radial distance mapping | Group comparison | PCL-R | Violent offenders | 51 | 51 | 0 | 2 |
| 38 | Boes et al. (2009) | MRI | Freesurfer | Correlational | PBS | na | 61 | 61 | 0 | 1 |
| 39 | Boes et al. (2008) | MRI | Freesurfer | Group comparison | PBS conduct scale | High PBS score | 40 | 40 | 0 | 1 |

| | | | | | | | | | | |
|---|---|---|---|---|---|---|---|---|---|---|
| 40 | Breitschuh et al. (2018) | MRI | VBM | Correlational | FAF | na | 26 | 26 | 0 | 1 |
| 41 | Brunnlieb et al. (2013) | fMRI | Modified TAGG | Activation study | AQ, IRI | na | 15 | 15 | 0 | 2 |
| 42 | Buades-Rotger et al. (2017) | fMRI | Modified TAGG | Activation study | na | na | 36 | 0 | 36 | 3 |
| 43 | Buckholtz et al. (2010) | fMRI | MID | Correlational | PPI IA-score | na | 24 | 8 | 16 | 1 |
| 44 | Budhiraja et al. (2019) | MRI | VBM | Group comparison | na | CD | 56 | 0 | 56 | 2 |
| 45 | Budhiraja et al. (2017) | MRI | Freesurfer | Group comparison | PCL-YV, PCL-SV | CD | 56 | 0 | 56 | 2 |
| 46a | Byrd et al. (2018) | fMRI | Card guessing task with chance of high or low monetary reward or loss | Group comparison | CBCL, APSD | Conduct problems | 51 | 51 | 0 | 1 |
| 46b | Byrd et al. (2018) | fMRI | Card guessing task with chance of high or low monetary reward or loss | Correlational | CBCL, APSD | Boys with conduct problems | 64 | 64 | 0 | 4 |
| 47 | Caldwell et al. (2019) | MRI | VBM | Correlational | ICU | Incarcerated male adolescents | 269 | 269 | 0 | 1 |
| 48 | Caldwell et al. (2015) | fMRI | Seeing pictures with and without moral violations | Correlational | PCL-R | Incarcerated male cocaine users | 87 | 87 | 0 | 1 |
| 48 | Caldwell et al. (2015) | fMRI | Seeing pictures with and without moral violations | Group comparison | PCL-R | Incarcerated male volunteers | 342 | 342 | 0 | 2 |
| 49a | Cardinale et al. (2018) | fMRI | Reading emotionally evocative statements and judging whether they would be acceptable to say to another person | Group comparison | SDQ, CBCL, ICU | Youths with clinical range scores in both SDQ and CBCL (conduct problems & externalizing) and high callous-unemotional traits | 26 | 18 | 8 | 3 |
| 49b | Cardinale et al. (2018) | fMRI | Reading emotionally evocative statements and judging whether they would be acceptable to say to another person | Group comparison | SDQ, CBCL, ICU | Youths with clinical range scores in both SDQ and CBCL (conduct problems & externalizing) and low callous-unemotional traits | 18 | 13 | 5 | 2 |
| 50 | Carlson et al. (2010) | fMRI | Neutral or fearful faces quickly masked with another neutral or fearful face | Correlational | STAXI-2, State-Trait Anxiety Inventory | na | 15 | 8 | 7 | 2 |
| 51a | Carré et al. (2013) | fMRI | Face-matching task with emotional faces (fearful, angry, surprised and neutral) | Activation study | BPAQ physical aggression scale | na | 64 | 28 | 36 | 1 |
| 51b | Carré et al. (2013) | fMRI | Face-matching task with emotional faces | Correlational | BPAQ physical aggression scale | na | 28 | 28 | 0 | 1 |
| 52 | Castellanos-Ryan et al. (2014) | fMRI | SST and a modified version of the MID | Correlational | Development and Well-Being Assessment interview, SDQ, Alcohol Use Disorders Identification Test | na | 1778 | 866 | 912 | 3 |
| 53 | Chang et al. (2017) | PET | 4-[18F]-ADAM | Group comparison | ASRS, BIS, WHOQOL-BREF | CD | 22 | 22 | 0 | 1 |
| 54 | Charpentier et al. (2016) | MRI | VBM | Correlational | Zuckerman Sensation Seeking Scale, Eysenck I7, in-house computerized modification of Section M of the SSAGA (antisocial acts) | na | 176 | 147 | 29 | 2 |
| 55 | Chen et al. (2021) | fMRI | PSAP | Activation study | 30-item Chinese version of the BPAQ | na | 34 | 18 | 16 | 2 |
| 56 | Chester & DeWall (2016) | fMRI | TAGG | Correlational | AMII Anger Expression-Out scale | na | 69 | 22 | 47 | 1 |
| 57 | Chester et al. (2017) | MRI | VBM | Correlational | BAQ | na | 138 | 47 | 91 | 2 |
| 58 | Chester et al. (2018) | fMRI | Cyberball social rejection task and a TAGG-like task | Correlational | BAQ, AMII | na | 60 | 22 | 38 | 1 |
| 59 | Choe et al. (2015) | fMRI | Emotional face matching and shape-matching | Correlational | BIS, criminal court records of arrests | na | 178 | 178 | 0 | 2 |
| 60a | Chumachenko et al. (2015) | MRI | Freesurfer | Group comparison | Youth Self Report, DISC-IV, Eysenck Junior Impulsiveness Scale, Peak Aggressive Behavior, CBCL | Adolescent males with serious substance use and conduct problems | 44 | 44 | 0 | 1 |
| 60b | Chumachenko et al. (2015) | MRI | Freesurfer | Correlational | Youth Self Report, DISC-IV, Eysenck Junior Impulsiveness Scale, Peak Aggressive Behavior rating scale, CBCL | Adolescent males with substance use and conduct problems | 25 | 25 | 0 | 1 |
| 61 | Coccaro et al. (2018) | MRI | VBM | Correlational | LHA aggression subscale | na | 287 | 145 | 142 | 1 |
| 62 | Coccaro et al. (2016) | MRI | VBM | Group comparison | LHA aggression scale, BPAQ aggression scale | IED | 168 | 83 | 85 | 1 |
| 63a | Coccaro et al. (2016a) | fMRI | Identifying emotional valence from emotional faces | Group comparison | LHA aggression scale, BPAQ verbal and physical aggression scores | IED | 56 | 18 | 38 | 1 |
| 63b | Coccaro et al. (2016a) | fMRI | Identifying emotional valence from emotional faces | Correlational | LHA aggression scale, BPAQ verbal and physical aggression scores | IED | 56 | 18 | 38 | 1 |
| 64 | Coccaro et al. (2007) | fMRI | Identifying gender from emotional faces | Group comparison | LHA aggression scale, BPAQ, STAXI | IED | 20 | 5 | 5 | 14 |
| 65a | Cohn et al. (2013) | fMRI | Classical differential delay fear conditioning task with two neutral faces as conditioned stimuli | Correlational | DISC-IV , YPI, RPQ | Adolescents who were first arrested by the police before the age of 12 and met the criteria for ODD or CD currently or previously in their history | 76 | 61 | 15 | 3 |
| 65b | Cohn et al. (2013) | fMRI | Classical differential delay fear conditioning task with two neutral faces as conditioned stimuli | Group comparison | DISC-IV, YPI, RPQ | Adolescents who were first arrested by the police before the age of 12 and with ODD or CD | 51 | 41 | 10 | 1 |
| 66a | Cohn et al. (2016) | fMRI | Classical differential delay fear conditioning task with two neutral faces as conditioned stimuli | Group comparison | DISC-IV, YPI | Adolescents who were first arrested by the police before the age of 12 and met the criteria for ODD or CD currently or previously in their history | 75 | 60 | 15 | 1 |
| 66b | Cohn et al. (2016) | fMRI | Classical differential delay fear conditioning task with two neutral faces as conditioned stimuli | Correlational | DISC-IV, YPI | Adolescents who were first arrested by the police before the age of 12 and met the criteria for ODD or CD currently or previously in their history | 75 | 60 | 15 | 1 |
| 67a | Cohn et al. (2015) | fMRI | MID | Correlational | DISC-IV, YPI | Adolescents who were first arrested by the police before the age of 12 and met the criteria for ODD or CD currently or previously in their history | 128 | 108 | 20 | 1 |
| 67b | Cohn et al. (2015) | fMRI | MID | Group comparison | DISC-IV, YPI | Adolescents who were first arrested by the police before the age of 12 and current ODD or CD | 45 | 36 | 9 | 2 |
| 68 | Cohn et al. (2016a) | MRI | VBM | Correlational | DISC-IV, YPI (callous-unemotional subscale), CBCL, Youth Self Report (YSR) | Childhood arrestees | 134 | 114 | 20 | 2 |
| 69a | Contreras-Rodríguez et al. (2014) | fMRI | Emotional face matching task | Group comparison | PCL-R | Psychopatic incarcerated violent armed offenders | 44 | 44 | 0 | 3 |
| 69b | Contreras-Rodríguez et al. (2014) | fMRI | Emotional face matching task | Correlational | PCL-R | Psychopatic incarcerated violent armed offenders | 44 | 44 | 0 | 2 |
| 70a | Contreras-Rodríguez et al. (2015) | MRI | VBM | Group comparison | PCL-R | Psychopatic incarcerated violent armed offenders | 44 | 44 | 0 | 2 |
| 70b | Contreras-Rodríguez et al. (2015) | MRI | VBM | Correlational | PCL-R | Psychopatic incarcerated violent armed offenders | 44 | 44 | 0 | 4 |
| 71 | Cope et al. (2014) | MRI | VBM | Group comparison | PCL-YV | Homicide offenders | 155 | 155 | 0 | 1 |
| 72 | Cope et al. (2014a) | MRI | VBM | Correlational | PCL-YV | Incarcerated adolescents | 39 | 0 | 39 | 1 |
| 73 | Cope et al. (2012) | MRI | VBM | Correlational | PCL-R | Individuals from probation, parole and drug-treatment centers, most with some psychiatric diagnosis | 66 | 36 | 30 | 2 |
| 74 | Cope et al. (2014b) | fMRI | Rating drug craving elicited by viewing neutral or drug-related pictures | Correlational | PCL-R | Drug dependent prisoners | 137 | 44 | 93 | 1 |
| 75 | Cropley et al. (2021) | MRI | FreeSurfer | Correlational | GOASSESS | Various medical conditions | 1313 | 654 | 659 | 4 |
| 76 | Crowley et al. (2010) | fMRI | CBG | Group comparison | DISC-IV | Abstinent adolescent male patients in treatment for antisocial substance disorder | 40 | 40 | 0 | 3 |
| 77 | Crowley et al. (2015) | fMRI | CBG | Group comparison | DISC-IV | Substance use disorder, CD | 81 | 40 | 41 | 2 |
| 78 | da Cunha-Bang et al. (2019) | fMRI | Emotional face matching task | Correlational | BPAQ, STAXI, NEO-PI-R , BIS, PPI-R | Incarcerated violent offenders | 47 | 47 | 0 | 1 |

| | | | | | | | | | | |
|---|---|---|---|---|---|---|---|---|---|---|
| 79a | da Cunha-Bang et al. (2017) | fMRI | PSAP | Group comparison | BPAQ, STAXI-2 | Incarcerated violent offenders with one or multiple personality disorders, most commonly antisocial | 44 | 44 | 0 | 2 |
| 79b | da Cunha-Bang et al. (2017) | fMRI | PSAP | Correlational | BPAQ, STAXI-2 | Incarcerated violent offenders with one or multiple personality disorders, most commonly antisocial | 44 | 44 | 0 | 2 |
| 80 | da Cunha-Bang et al. (2016) | PET | [11C]SB207145 | Correlational | BPAQ | na | 36 | 36 | 0 | 1 |
| 81a | Dalwani et al. (2011) | MRI | VBM | Group comparison | DISC, CIDI-SAM, Peak Aggression Scale, EIS | CD and at least one non-nicotine substance use disorder | 44 | 44 | 0 | 2 |
| 81b | Dalwani et al. (2011) | MRI | VBM | Correlational | DISC, CIDI-SAM, Peak Aggression Scale, EIS | Adolescents referred to treatment with CD and at least one non-nicotine substance use disorder | 19 | 19 | 0 | 1 |
| 82a | Dalwani et al. (2014) | fMRI | CBG | Group comparison | na | Abstinent adolescent male patients in treatment for antisocial substance disorder | 40 | 40 | 0 | 2 |
| 82b | Dalwani et al. (2014) | fMRI | CBG | Correlational | na | Abstinent adolescent male patients in treatment for antisocial substance disorder | 40 | 40 | 0 | 1 |
| 83 | Dambacher et al. (2014) | fMRI | TAGG | Activation study | na | na | 15 | 15 | 0 | 1 |
| 84 | De Brito et al. (2009) | MRI | VBM | Group comparison | SDQ conduct problem scale, ASPD callous-unemotional trait scale | Conduct problems and callous-unemotional traits | 48 | 48 | 0 | 2 |
| 85a | de Oliveira-Souza et al. (2008) | MRI | VBM | Group comparison | PCL:SV | ASPD | 30 | 16 | 14 | 2 |
| 85b | de Oliveira-Souza et al. (2008) | MRI | VBM | Correlational | PCL:SV | ASPD | 30 | 16 | 14 | 1 |
| 86 | Decety & Porges (2011) | fMRI | Imagining being a part of situations of either interpersonal help or harm (or neutral baseline) presented via animations | Activation study | na | na | 22 | 22 | 0 | 4 |
| 87a | Decety et al. (2013) | fMRI | Taking either a self-perspective or an other-perspective while viewing visual stimuli of painful and non-painful situations | Group comparison | PCL-R | Psychopathic criminals | 77 | 77 | 0 | 4 |
| 87b | Decety et al. (2013) | fMRI | Taking either a self-perspective or an other-perspective while viewing visual stimuli of painful and non-painful situations | Correlational | PCL-R | Psychopathic criminals | 121 | 121 | 0 | 8 |
| 88 | Decety et al. (2015) | fMRI | Matching emotional faces to probable reactions of perpetrator/agent or victim/beneficiary in morally laden social interactions | Group comparison | PCL-R | Psychopathic criminals | 88 | 88 | 0 | 8 |
| 89a | Decety et al. (2009) | fMRI | Seeing painful situations (pain/no-pain, self-inflicted/inflicted by other person) | Group comparison | DBD checklist, DISC, Child and Adolescent Disposition Scale | Adolescents with aggressive CD | 16 | | | 2 |
| 89b | Decety et al. (2009) | fMRI | Seeing painful situations (pain/no-pain, self-inflicted/inflicted by other person) | Correlational | DBD checklist, DISC, Child and Adolescent Disposition Scale | Adolescents with aggressive CD | 8 | | | 2 |
| 90 | Decety et al. (2014) | fMRI | Emotional faces | Group comparison | PCL-R | Psychopathic criminals | 46 | 46 | 0 | 8 |
| 91a | Decety et al. (2013a) | fMRI | Videos of persons harming one another and viewing dynamic facial expressions of pain | Group comparison | PCL-R | Psychopathic criminals | 46 | 46 | 0 | 4 |
| 91b | Decety et al. (2013a) | fMRI | Videos of persons harming one another and viewing dynamic facial expressions of pain | Correlational | PCL-R | Psychopathic criminals | 70 | 70 | 0 | 8 |
| 92 | Deeley et al. (2006) | fMRI | Identifying gender from emotional faces | Group comparison | PCL-R | Repeat offenders with PCL-R score of 25 or above | 15 | 15 | 0 | 2 |
| 93 | Delfin et al. (2019) | SPECT | 99mTc-exametazime | Group comparison | na | Recidivist forensic psychiatric patients | 44 | 39 | 5 | 1 |
| 94a | Deming et al. (2020) | fMRI | Matching either emotional faces or shapes to obscured faces in images portraying two individuals interacting | Correlational | PCL-R | Adult inmates | 94 | 94 | 0 | 2 |
| 94b | Deming et al. (2020) | fMRI | Matching either emotional faces or shapes to obscured faces in images portraying two individuals interacting | Group comparison | PCL-R | Inmates | 59 | 59 | 0 | 1 |
| 95 | Deming et al. (2018) | fMRI | Making yes/no judgements about trait adjectives about self, others, or case | Correlational | PCL-R | Incarcerated offenders | 57 | 57 | 0 | 3 |
| 96 | Denson et al. (2009) | fMRI | Ruminating on provocation, self or thinking about affectively neutral statements after provocation/insult | Correlational | BPAQ, Displaced Aggression Questionnaire, Profile of Mood States (POMS) anger/hostility scale, Positive and Negative Affect Schedule (PANAS-X) hostility scale | na | 20 | 8 | 12 | 3 |
| 97 | Dolan et al. (2002) | MRI | Manual tracing method | Group comparison | Special Hospital Assessment of Personality and Socialisation (SHAPS) (now known as the Antisocial Personality Questionnaire) | Psychopathic violent offenders | 29 | 29 | 0 | 3 |
| 98 | Dolan & Fullam (2009) | fMRI | Identifying gender from pictures of emotional faces | Group comparison | PCL:SV | Violent offenders with SCZ and high psychopathy scores | 24 | 24 | 0 | 2 |
| 99 | Dong et al. (2017) | fMRI | Judging whether or not actors felt pain when they either depicted a neutral expression while receiving nonpainful stimulation or a painful expression while receiving painful stimulation | Group comparison | IRI scale, SDQ, BPAQ, APSD | Adolescents with CD | 66 | 66 | 0 | 1 |
| 100 | Dotterer et al. (2017) | fMRI | Matching fearful or angry faces | Correlational | Self-Report of Delinquency Questionnaire (SRD), five item sum from APSD callous-unemotional scale | na | 220 | 111 | 109 | 1 |
| 101 | Dougherty et al. (2006) | PET | [11C]SCH 23,390 | Group comparison | na | Patients with MDD and anger attacks | 20 | 10 | 10 | 1 |
| 102 | Emmerling et al. (2016) | fMRI | TAGG | Activation study | na | na | 15 | 15 | 0 | 5 |
| 103 | Ermer et al. (2013) | MRI | VBM | Correlational | PCL-YV | Incarcerated adolescents | 191 | 191 | 0 | 2 |
| 104 | Ermer et al. (2012) | MRI | VBM | Correlational | PCL-R | Incarcerated males | 296 | 296 | 0 | 2 |
| 105 | Ewbank et al. (2018) | fMRI | Identifying gender from emotional faces | Group comparison | YPI | Adolescents with CD | 71 | 71 | 0 | 2 |
| 106 | Fahim et al. (2012) | MRI | VBM | Group comparison | The Dominic-R interactive | ODD | 38 | 20 | 18 | 2 |
| 107 | Fahim et al. (2011) | MRI | VBM | Group comparison | The Dominic-R interactive | ODD or CD | 47 | 23 | 24 | 2 |
| 108a | Fairchild et al. (2014) | fMRI | Identifying gender from emotional faces | Group comparison | YPI, ICU | Adolescents with CD | 36 | 0 | 36 | 2 |
| 108b | Fairchild et al. (2014) | fMRI | Identifying gender from emotional faces | Correlational | YPI, ICU | Adolescents with CD | 18 | 0 | 18 | 1 |
| 109a | Fairchild et al. (2013) | MRI | VBM | Group comparison | YPI | Adolescents with CD | 42 | 0 | 42 | 2 |
| 109b | Fairchild et al. (2013) | MRI | VBM | Correlational | YPI | Adolescents with CD | 42 | 0 | 42 | 2 |
| 110 | Fairchild et al. (2011) | MRI | VBM | Group comparison | YPI, ICU | CD | 90 | 90 | 0 | 1 |
| 111a | Fairchild et al. (2015) | MRI | Freesurfer | patiens/control | YPI callous-unemotional scale | CD | 56 | 56 | 0 | 3 |
| 111b | Fairchild et al. (2015) | MRI | Freesurfer | Correlational | YPI callous-unemotional scale | Adolescents with CD | 36 | 36 | 0 | 2 |
| 112 | Fede et al. (2016) | fMRI | Judging the morality of morally wrong, not wrong or controversial acts or concepts | Correlational | PCL-R | Incarcerated adults | 245 | 245 | 0 | 2 |
| 113 | Fehlbaum et al. (2018) | fMRI | AST | Group comparison | YPI | Youths with CD | 78 | 58 | 20 | 2 |
| 114 | Fehr et al. (2014) | fMRI | First person videos presenting neutral, social-positive or reactive-aggressive interactions | Activation study | na | na | 20 | 20 | 0 | 2 |
| 115 | Finger et al. (2011) | fMRI | PAT | Group comparison | APSD, PCL:YV | Youths with psychopathic traits and either CD or ODD | 30 | 18 | 12 | 1 |

| | | | | | | | | | | |
|---|---|---|---|---|---|---|---|---|---|---|
| 116 | Finger et al. (2008) | fMRI | Probabilistic reversal learning task | Group comparison | APSD, PCL:YV | Youths with psychopathic traits and either CD or ODD | 28 | 18 | 10 | 1 |
| 117 | Foell et al. (2016) | fMRI | Passive viewing or suppressing emotions while viewing pleasant, unpleasant and neutral pictures | Correlational | 30 item trait disinhibition scale comprised of Exterlanizing Spectrum Inventory items | na | 45 | 0 | 45 | 2 |
| 118 | Frankle et al. (2005) | PET | [11C]McN 5652 | Group comparison | na | IED | 20 | 10 | 10 | 1 |
| 119 | Freeman et al. (2015) | fMRI | visual Go/NoGo task | Group comparison | PCL-R | Psychopathic criminals | 44 | 44 | 0 | 1 |
| 120 | Fullam et al. (2009) | fMRI | Either lying or telling the truth about everyday acts | Correlational | PPI | na | 24 | 24 | 0 | 5 |
| 121 | Gan et al. (2016) | fMRI | PSAP | Group comparison | structured clinical interview, STAXI-2 | Full or subclinical IED | 18 | 18 | 0 | 2 |
| 122 | Gansler et al. (2011) | MRI | Manual tracing with Medx 3.4.2 software's manual tracing tool | Correlational | LHA - Revised Aggression subscale (LHA-R-Agg), BIS-11 | Various psychiatric patients | 36 | 36 | 0 | 3 |
| 123 | Gansler et al. (2009) | MRI | Manual tracing | Correlational | LHA - Revised (LHA-R), BIS | Various psychiatric patients | 41 | 36 | 5 | 1 |
| 124a | Gao et al. (2021) | MRI | VBM | Group comparison | SDQ, BPAQ | CD | 182 | 182 | 0 | 3 |
| 124b | Gao et al. (2021) | MRI | VBM | Correlational | SDQ, BPAQ | CD | 96 | 96 | 0 | 1 |
| 124c | Gao et al. (2021) | MRI | VBM | Correlational | SDQ, BPAQ | na | 86 | 86 | 0 | 2 |
| 125a | Gard et al. (2017) | fMRI | Emotional face matching | Correlational | Structured Clinial Interview for DSM-IV Axis II Personality Disorders | na | 310 | 310 | 0 | 1 |
| 125b | Gard et al. (2017) | fMRI | Emotional face matching task | Activation study | Structured Clinial Interview for DSM-IV Axis II Personality Disorders | na | 167 | 167 | 0 | 1 |
| 126a | George et al. (2004) | PET | 18FDG | Group comparison | Brown-Goodwin Lifetime Aggression Scale | Alcohol dependent male perpetrators of domestic violence | 18 | 18 | 0 | 1 |
| 126b | George et al. (2004) | PET | 18FDG | Group comparison | Brown-Goodwin Lifetime Aggression Scale | Alcohol dependent male perpetrators of domestic violence | 19 | 19 | 0 | 1 |
| 126c | George et al. (2004) | PET | 18FDG | Correlational | Brown-Goodwin Lifetime Aggression Scale | Alcohol dependent male perpetrators of domestic violence | 8 | 8 | 0 | 1 |
| 126d | George et al. (2004) | PET | 18FDG | Correlational | Brown-Goodwin Lifetime Aggression Scale | na | 10 | 10 | 0 | 1 |
| 127a | Gerra et al. (1998) | SPECT | 99m Tchexamethyl-propylene-amine oxime (HMPAO) | Group comparison | personality disorders questionnaire -revised (PDQ-R) | Abstinent heroin addicts with lifetime ASPD symptoms | 18 | 13 | 5 | 1 |
| 127b | Gerra et al. (1998) | SPECT | 99m Tchexamethyl-propylene-amine oxime (HMPAO) | Group comparison | personality disorders questionnaire -revised (PDQ-R) | Abstinent heroin addicts with lifetime ASPD symptoms | 18 | 13 | 5 | 1 |
| 128 | Geurts et al. (2016) | fMRI | MID | Group comparison | PCL-R, PPI | Psychopathic criminals and healthy subjects with PPI impulsive/antisocial traits subscale score over 146 | 34 | 34 | 0 | 1 |
| 129 | Glenn et al. (2017) | fMRI | Either lying or answering truthfully to either autobiographical or non-autobiographical questions concerning either criminal or noncriminal matters | Correlational | PCL-R: 2nd edition | na | 16 | 14 | 2 | 16 |
| 130a | Goldstein et al. (2005) | PET | 18FDG | Correlational | MMPI-2 anger content scale | Recently abstinent cocaine-dependent subjects | 17 | 17 | 0 | 1 |
| 130b | Goldstein et al. (2005) | PET | 18FDG | Correlational | MMPI-2 anger content scale | na | 16 | 16 | 0 | 1 |
| 131 | Gopal et al. (2013) | MRI | Manual tracing | Correlational | LHA - Revised (LHA-R), BIS | Various psychiatric patients | 41 | 36 | 5 | 2 |
| 132 | Gordon et al. (2004) | fMRI | Matching emotional pictures either by identity or emotion | Group comparison | PPI | Healthy males who scored high on the PPI emotional-interpersonal subscale | 18 | 18 | 0 | 2 |
| 133 | Gorka et al. (2015) | MRI | VBM | Correlational | BPAQ | na | 253 | 0 | 253 | 1 |
| 134 | Goyer et al. (1994) | PET | [18F]FDG | Correlational | modified LHA | Various personality disorders | 60 | 33 | 27 | 1 |
| 135a | Gregory et al. (2015) | fMRI | Probabilistic response reversal task | Group comparison | PCL-R | Psychopathic offenders with ASPD | 30 | 30 | 0 | 2 |
| 135b | Gregory et al. (2015) | fMRI | Probabilistic response reversal task | Group comparison | PCL-R | Psychopathic offenders with ASPD | 32 | 32 | 0 | 2 |
| 136a | Gregory et al. (2012) | MRI | VBM | Group comparison | PCL-R, RPQ | Psychopathic offenders with ASPD | 39 | 39 | 0 | 1 |
| 136b | Gregory et al. (2012) | MRI | VBM | Group comparison | PCL-R, RPQ | Psychopathic offenders with convictions for violent crimes and ASPD | 44 | 44 | 0 | 1 |
| 137 | Han et al. (2012) | fMRI | Partial face encoding task | Group comparison | PPI -revised (PPI-R) | Healthy participants in the top tertile for PPI coldheartedness score | 36 | 13 | 19 | 2 |
| 138 | Harenski et al. (2014) | fMRI | Rating the severity of moral transgression in moral, non-moral and neutral pictures | Correlational | PCL-R | Incarcerated volunteers | 157 | 0 | 157 | 2 |
| 139 | Harenski et al. (2014a) | fMRI | Rating the severity of moral transgression in moral, non-moral and neutral pictures | Correlational | PCL-YV, Inventory of Callous and Unemotional Traits - Youth Version (ICU-Y), KSADS-PL | Incarcerated adolescents | 111 | 111 | 0 | 6 |
| 140 | Harenski et al. (2010) | fMRI | Rating the severity of moral transgression in moral, non-moral and neutral pictures | Correlational | PCL-R | Incarcerated individuals | 72 | 72 | 0 | 2 |
| 141 | Harenski et al. (2012) | fMRI | Rating the severity of pain caused in painful and non painful pictures | Group comparison | Severe Sexual Sadism Scale | Sadistic male sexual offenders | 15 | 15 | 0 | 1 |
| 142 | Heesink et al. (2017) | fMRI | Fear-and-Escape Task (virtual-predator task) | Group comparison | STAXI-2, BPAQ | Military veterans with anger/impulsive aggression problems | 57 | 57 | 0 | 2 |
| 143 | Heesink et al. (2018) | fMRI | Identifying emotional valence from pictures that elicit emotions | Group comparison | STAXI-2, BPAQ | Military veterans with anger/impulsive aggression problems | 56 | 56 | 0 | 1 |
| 144a | Herpertz et al. (2008) | fMRI | Emotional pictures | Group comparison | KIDDIE-SADS, CBCL, German Parental and Teacher Report on ADHD and CD symptoms (FBB-HKS and FBB-SSV) | CD | 44 | 44 | 0 | 1 |
| 144b | Herpertz et al. (2008) | fMRI | Passive viewing of emotional pictures | Correlational | K-SADS, CBCL, German Parental and Teacher Report on ADHD and CD symptoms (FBB-HKS and FBB-SSV) | CD | 22 | 22 | 0 | 2 |
| 145 | Hirono et al. (2000) | SPECT | Technetium Tc 99m-labeled hexamethylpropylene amineoxime | Group comparison | neuropsychiatry inventory (NPI) | Aggressive dementia patients | 20 | 8 | 12 | 1 |
| 146 | Hofhansel et al. (2020) | MRI | VBM | Correlational | PCL-R, BPAQ, RPQ | Violent offenders | 54 | 54 | 0 | 6 |
| 147 | Holz et al. (2015) | MRI | VBM | Correlational | Mannheim Parent Interview, the Schedule for Affective Disorders and Schizophrenia for School-Age Children | na | 167 | 67 | 100 | 1 |
| 148 | Hoptman et al. (2006) | MRI | segmented with EMS (Expectation Maximization Segmentation) and IRIS software | Correlational | OAS log-transformed total aggression severity score, PANSS Hostility & Poor Impulse Control items and aggressive incidents during study period | Hospitalized treatment-resistant patients with SCZ or schizoaffective disorder | 49 | 43 | 6 | 3 |
| 149 | Hoptman et al. (2005) | MRI | segmented with EMS (Expectation Maximization Segmentation) and IRIS software, manual tracing | Correlational | OAS log-transformed total aggression severity score and PANSS Hostility item | Hospitalized treatment-resistant patients with SCZ or schizoaffective disorder | 49 | 43 | 6 | 1 |
| 150 | Hosking et al. (2017) | fMRI | Choosing between larger but later or smaller but sooner monetary rewards | Correlational | PCL-R | Incarcerated offenders | 45 | 45 | 0 | 1 |
| 151a | Howner et al. (2012) | MRI | the FACE program | Group comparison | PCL-SV | Psychopathic offenders | 19 | 19 | 0 | 1 |
| 151b | Howner et al. (2012) | MRI | the FACE program | Correlational | PCL-SV | Psychopathic offenders and offenders with autism spectrum disorders | 26 | 26 | 0 | 1 |
| 152 | Huang et al. (2019) | fMRI | MID | Correlational | CBCL, ICU | na | 29 | 14 | 15 | 2 |
| 153 | Huber et al. (2018) | MRI | VBM | Group comparison | Brief psychiatric rating scale - excitement component (BPRS-EC) | At-risk mental state or first-episode psychosis patients with agitated-aggressive syndrome | 111 | 81 | 30 | 1 |

| | | | | | | | | | | |
|---|---|---|---|---|---|---|---|---|---|---|
| 154 | Huebner et al. (2008) | MRI | VBM | Group comparison | K-SADS | CD | 46 | 46 | 0 | 2 |
| 155 | Hwang et al. (2018) | fMRI | Social and non-social reinforcement-learning task | Group comparison | K-SADS | Disruptive behavior disorders | 48 | 26 | 22 | 2 |
| 156a | Hwang et al. (2016) | fMRI | AST | Group comparison | ICU - Parent Version (ICU-P), K-SADS | Disruptive behavior disorders and high callous-unemotional traits | 46 | 25 | 21 | 3 |
| 156b | Hwang et al. (2016) | fMRI | AST | Group comparison | ICU - Parent Version (ICU-P), K-SADS | Youth with disruptive behavior disorders and low callous-unemotional traits | 45 | 27 | 18 | 3 |
| 156c | Hwang et al. (2016) | fMRI | AST | Group comparison | ICU- Parent Version (ICU-P), K-SADS | Youth with disruptive behavior disorders and high callous-unemotional traits | 35 | 22 | 13 | 3 |
| 157 | Hyatt et al. (2012) | MRI | Freesurfer | Group comparison | K-SADS-PL | Adolescents with CD | 43 | 24 | 19 | 2 |
| 158 | Hyde et al. (2014) | fMRI | Emotional face matching task | Correlational | NEO-PI-R (expert consensus prototype matching approach), multidimensional personality questionnaire - brief form (MPQ-BF) | na | 103 | 46 | 57 | 4 |
| 159 | Ibrahim et al. (2019) | fMRI | Identifying gender from emotional faces | Correlational | K-SADS-PL, ICU, CBCL | Autism spectrum disorder and disruptive behavior | 38 | 32 | 6 | 2 |
| 160a | Intrator et al. (1997) | SPECT | 99mTc-hexamethylpropyleneamine oxime (99mTc-HMPAO) | Group comparison | PCL-R | Male patients in an inpatient substance abuse program with PCL-R scores of at least 25 | 17 | 17 | 0 | 1 |
| 160b | Intrator et al. (1997) | SPECT | 99mTc-hexamethylpropyleneamine oxime (99mTc-HMPAO) | Group comparison | PCL-R | Male patients in an inpatient substance abuse program with PCL-R scores of at least 25 | 17 | 17 | 0 | 1 |
| 161 | Jiang et al. (2016) | MRI | Freesurfer | Group comparison | na | ASPD | 52 | 52 | 0 | 2 |
| 162 | Jones et al. (2009) | fMRI | Identifying gender from emotional faces | Group comparison | SDQ conduct problems -skaala, APSD | Conduct problems and callous-unemotional traits | 30 | 30 | 0 | 1 |
| 163 | Jones et al. (2017) | fMRI | Reappraisal task | Group comparison | na | Juvenile sexual offenders | 39 | 39 | 0 | 2 |
| 164a | Joyal et al. (2007) | fMRI | Go/no-go task | Group comparison | na | Homicide offenders with primary diagnosis of SCZ | 24 | 24 | 0 | 2 |
| 164b | Joyal et al. (2007) | fMRI | Go/no-go task | Group comparison | na | Homicide offenders with primary diagnosis of SCZ | 24 | 24 | 0 | 2 |
| 164c | Joyal et al. (2007) | fMRI | Go/no-go task | Group comparison | na | Homicide offenders with primary diagnosis of SCZ | 24 | 24 | 0 | 2 |
| 165 | Juhász et al. (2001) | PET | Fluorodexyglucose (FDG) | Group comparison | CBCL | Children with partial epilepsy and significant aggressive behavior that could not be attributed to medication side effects | 13 | 7 | 6 | 2 |
| 166a | Kiehl et al. (2001) | fMRI | Affective memory task | Group comparison | PCL-R | Criminal psychopaths | 16 | | | 4 |
| 166b | Kiehl et al. (2001) | fMRI | Affective memory task | Group comparison | PCL-R | Criminal psychopaths | 16 | | | 5 |
| 167 | Kiehl et al. (2004) | fMRI | Lexical decision task | Group comparison | PCL-R | Criminal psychopaths | 16 | 16 | 0 | 1 |
| 168 | Kim & James (2015) | fMRI | Suppressing emotions or just watching emotional pictures | Correlational | Aggressive Behavior Inventory (completed by significant other) | na | 17 | | | 1 |
| 169 | Kirino et al. (2019) | fMRI | Sound omission mismatch negativity paradigm | Correlational | PANSS | SCZ | 12 | 10 | 2 | 1 |
| 170a | Klapwijk et al. (2016) | fMRI | Explicit empathy task | Group comparison | K-SADS-PL, ICU | CD and callous-unemotional traits | 56 | 56 | 0 | 2 |
| 170b | Klapwijk et al. (2016) | fMRI | Explicit empathy task (judge either own or presented emotional state with emotional faces as stimuli and a perceptual decision as a control) | Correlational | K-SADS-PL, ICU | Adolescents with CD and callous-unemotional traits | 23 | 23 | 0 | 1 |
| 171 | Klapwijk et al. (2016a) | fMRI | Dictator game with info about prior emotional reaction to unfair decision provided to the participant | Group comparison | ICU | Adolescents with CD | 65 | 65 | 0 | 3 |
| 172 | Klasen et al. (2020) | fMRI | Playing Carmaggedon: TDR 2000 in either a violent (killing pedestrians) or nonviolent (collect points) scenario | Activation study | na | na | 15 | 15 | 0 | 1 |
| 173a | Kolla et al. (2021) | PET | [11C]CURB | Group comparison | urgency, premediation, perseverance, sensation seeking, positive urgency impulsive behavior scale (UPPS-P), BDHI, RPQ, PCL:SV, Triarchic psychopathy measure (TriPM) | ASPD and a history of violent offending | 32 | 32 | 0 | 1 |
| 173b | Kolla et al. (2021) | PET | [11C]CURB | Correlational | urgency, premediation, perseverance, sensation seeking, positive urgency impulsive behavior scale (UPPS-P), BDHI, RPQ, PCL:SV, Triarchic psychopathy measure (TriPM) | ASPD and a history of violent offending | 32 | 32 | 0 | 2 |
| 174 | Kolla et al. (2014) | MRI | VBM | Group comparison | PCL-R | Psychopathic violent offenders with ASPD | 24 | 24 | 0 | 2 |
| 175a | Kolla et al. (2015) | PET | [11C] harmine | Group comparison | NEO PI-R, PCL-R, Iowa gambling task | ASPD patients with histories of impulsive violent offending | 36 | 36 | 0 | 1 |
| 175b | Kolla et al. (2015) | PET | [11C] harmine | Correlational | NEO PI-R, PCL-R, Iowa gambling task | ASPD patients with histories of impulsive violent offending | 18 | 18 | 0 | 2 |
| 176 | Korponay et al. (2017) | MRI | VBM | Correlational | PCL-R | Prisoners | 124 | 124 | 0 | 1 |
| 177 | Korponay et al. (2017) | MRI | VBM | Correlational | PCL-R | Prisoners | 124 | 124 | 0 | 2 |
| 178 | Krämer et al. (2007) | fMRI | TAGG | Activation study | na | na | 15 | | | 2 |
| 179 | Kuikka et al. (1998) | SPECT | [123I]β-CIT | Group comparison | na | Alcoholic violent offenders | 42 | | | 1 |
| 180a | Kumari et al. (2006) | fMRI | n-back task | Group comparison | clinical and forensical records of violence, Gunn and Robertson's 1976 violence scale | Violent SCZ patients | 25 | 25 | 0 | 1 |
| 180b | Kumari et al. (2006) | fMRI | n-back task | Group comparison | clinical and forensical records of violence, Gunn and Robertson's 1976 violence scale | ASPD patients with a history of violence | 23 | 23 | 0 | 2 |
| 181a | Kumari et al. (2009) | MRI | MEASURE program and Cavalieri method | Group comparison | Gunn and Robertson Scale for Violence | SCZ patients with histories of serious violence | 24 | 24 | 0 | 1 |
| 181b | Kumari et al. (2009) | MRI | MEASURE program and Cavalieri method | Group comparison | Gunn and Robertson Scale for Violence | SCZ patients with histories of serious violence | 24 | 24 | 0 | 1 |
| 181c | Kumari et al. (2009) | MRI | MEASURE program and Cavalieri method | Correlational | Gunn and Robertson Scale for Violence, IVE-7 | SCZ patients with histories of serious violence | 38 | 38 | 0 | 1 |
| 182a | Kumari et al. (2009a) | fMRI | Alternating between the threat of an electric schock and a safe condition | Group comparison | Gunn and Robertson Scale for Violence | SCZ patients with histories of serious violence | 27 | 27 | 0 | 2 |
| 182b | Kumari et al. (2009a) | fMRI | Alternating between the threat of an electric schock and a safe condition | Group comparison | Gunn and Robertson Scale for Violence | SCZ patients with histories of serious violence | 26 | 26 | 0 | 1 |
| 182c | Kumari et al. (2009a) | fMRI | Alternating between the threat of an electric schock and a safe condition | Group comparison | Gunn and Robertson Scale for Violence | SCZ or ASPD patients with histories of serious violence | 53 | 53 | 0 | 1 |
| 183 | Kumari et al. (2013) | MRI | MEASURE program and Cavalieri method | Group comparison | Gunn and Robertson Scale for Violence | SCZ patients with histories of serious violence | 43 | 43 | 0 | 1 |
| 184a | Kumari et al. (2014) | MRI | MEASURE program and Cavalieri method | Group comparison | Gunn and Robertson Scale for Violence | ASPD | 29 | 29 | 0 | 1 |
| 184b | Kumari et al. (2014) | MRI | MEASURE program and Cavalieri method | Group comparison | Gunn and Robertson Scale for Violence | SCZ patients with histories of serious violence | 28 | 28 | 0 | 1 |
| 185 | Kuroki et al. (2017) | MRI | VBM | Group comparison | na | SCZ patients with histories serious violence | 58 | 58 | 0 | 1 |
| 186 | Kuruoğlu et al. (1996) | SPECT | 99mTc-HMPAO | Group comparison | na | Chronic alcoholics on remission diagnosed with ASPD | 25 | 25 | 0 | 1 |
| 187 | Kärgel et al. (2017) | fMRI | Go/no-go | Group comparison | na | Pedophilic sexual offenders | 77 | 77 | 0 | 1 |

| | | | | | | | | | | |
|---|---|---|---|---|---|---|---|---|---|---|
| 188 | Laakso et al. (2003) | PET | [18F]fluorodopa | Correlational | Karolinska Scales of Personality | na | 33 | 22 | 11 | 1 |
| 189 | Laakso et al. (2002) | MRI | Semi-automatic segmentation technique, which combines tracing and thresholding, manual tracing for subregional measurements | Group comparison | PCL-R | Patients with ASPD, early-onset type 2 alcoholism and history of violent offences | 57 | 57 | 0 | 1 |
| 190 | Laakso et al. (2001) | MRI | Manual tracing | Correlational | PCL-R | Violent offenders with ASPD and type 2 alcoholism | 18 | 18 | 0 | 1 |
| 191 | Lam et al. (2017) | MRI | Freesurfer | Correlational | PCL-R | Accused murderers, some with SCZ | 67 | 56 | 11 | 2 |
| 192 | Larson et al. (2013) | fMRI | Focusing either on a threat (electric shock risk) related or a nonrelevant stimulus | Group comparison | PCL-R | Psychopathic prisoners | 49 | 49 | 0 | 2 |
| 193 | Lasko et al. (2019) | MRI | VBM | Correlational | Levenson's Self-Report Psychopathy Scale (LSRP), the Shork Dark Triad (SD3) | na | 144 | 69 | 75 | 3 |
| 194 | Lawrence & Brooks (2014) | PET | 6-[18F]Fluoro-L-DOPA (FDOPA) | Correlational | Cloniger's Tri-dimensional Personality Questionnaire (TPQ) | na | 12 | 12 | 0 | 1 |
| 195 | Lee et al. (2009) | fMRI | Neutral, positive and violent pictures | Group comparison | STAXI, BIS | Domestic violence offenders | 23 | 23 | 0 | 4 |
| 196a | Leutgeb et al. (2015) | MRI | VBM | Group comparison | PCL-R, Violence Risk Appraisal Guide (VRAG), Violence Risk Scale (VRS), STAXI | High-risk violent offenders | 77 | 77 | 0 | 2 |
| 196b | Leutgeb et al. (2015) | MRI | VBM | Correlational | PCL-R, Violence Risk Appraisal Guide (VRAG), Violence Risk Scale (VRS), STAXI | High-risk violent offenders | 40 | 40 | 0 | 5 |
| 196c | Leutgeb et al. (2015) | MRI | VBM | Correlational | PCL-R, Violence Risk Appraisal Guide (VRAG), Violence Risk Scale (VRS), STAXI | High-risk violent offenders | 37 | 37 | 0 | 2 |
| 197 | Li et al. (2006) | fMRI | Script-guided imagery paradigm to induce emotional distress | Correlational | California Psychological Inventory socialization scale (CPI-so) | Abstinent cocaine abusers | 10 | 0 | 10 | 1 |
| 198a | Liu et al. (2020) | MRI | VBM | Group comparison | modified OAS (MOAS), PANSS | Violent SCZ patients | 75 | 52 | 23 | 1 |
| 198b | Liu et al. (2020) | MRI | VBM | Group comparison | modified OAS (MOAS), PANSS | Violent SCZ patients | 76 | 51 | 25 | 1 |
| 199a | Liu et al. (2012) | fMRI | Go/noGo -task | Correlational | Zuckerman-Kuhlman Personality Questionnaire (ZKPQ), NEO-PI-R | na | 15 | 0 | 15 | 3 |
| 199b | Liu et al. (2012) | fMRI | Go/noGo -task | Correlational | Zuckerman-Kuhlman Personality Questionnaire (ZKPQ), NEO-PI-R | na | 13 | 13 | 0 | 2 |
| 200a | Lockwood et al. (2013) | fMRI | Viewing pictures showing another person's hand or foot in painful or nonpainful situations | Correlational | Child and Adolescent Symptom Inventory (CASI-4R), ICU | Children with CD | 36 | 36 | 0 | 2 |
| 200b | Lockwood et al. (2013) | fMRI | Viewing pictures showing another person's hand or foot in painful or nonpainful situations | Group comparison | Child and Adolescent Symptom Inventory (CASI-4R), ICU | CD | 55 | 55 | 0 | 2 |
| 201a | Lotze et al. (2007) | fMRI | TAGG | Correlational | Self-report psychopathy scale (SRPS) primary psychopathy scale | na | 14 | 14 | 0 | 1 |
| 201b | Lotze et al. (2007) | fMRI | TAGG | Group comparison | Self-report psychopathy scale (SRPS) primary psychopathy scale | Participants with above average scores from SRPS primary psychopathy scale | 14 | 14 | 0 | 2 |
| 202 | Lozier et al. (2014) | fMRI | Emotional faces | Correlational | SDQ, CBCL, ICU, RPQ | Conduct problems | 46 | 26 | 20 | 6 |
| 203a | Lundwall et al. (2017) | MRI | Freesurfer | Correlational | Aberrant Behavior Checklist Irritability scale | Aggressive children with an autism spectrum disorder | 32 | 32 | 0 | 1 |
| 203b | Lundwall et al. (2017) | MRI | Freesurfer | Correlational | Aberrant Behavior Checklist Irritability scale | Aggressive children with autism spectrum disorder | 45 | 45 | 0 | 1 |
| 204 | Ly et al. (2012) | MRI | Freesurfer | Group comparison | PCL-R | Psychopathic inmates | 52 | 52 | 0 | 1 |
| 205 | Mackey et al. (2017) | MRI | VBM | Correlational | Kirby's Monetary Choice Questionnaire (MCQ), Life Events Questionnaire (LEQ) | na | 1741 | 846 | 895 | 2 |
| 206 | Marín-Lahoz et al. (2020) | MRI | VBM | Correlational | BIS-11, Iowa Gambling task (IGT), Balloon analogue risk task (BART), delay discounting task (DDT) | PD | 89 | 55 | 34 | 7 |
| 207a | Marín-Lahoz et al. (2020a) | PET | 18F-FDG | Group comparison | Questionnaire for impulsive-compulsive disorders (QUIP) | Nondemented PD patients with ICDs | 24 | 13 | 11 | 1 |
| 207b | Marín-Lahoz et al. (2020a) | PET | 18F-FDG | Group comparison | Questionnaire for impulsive-compulsive disorders (QUIP) | Nondemented PD patients with ICDs | 24 | 13 | 11 | 1 |
| 208a | Marín-Morales et al. (2021) | fMRI | Emotion regulation task (passively observing, attending to emotions or suppressing emotions while viewing neutral, overall negative or interpersonal violence-related pictures) | Group comparison | Conflict Tactics Scale-2 (CTS2), IRI | Domestic violence offenders | 55 | 55 | 0 | 2 |
| 208b | Marín-Morales et al. (2021) | fMRI | Emotion regulation task (passively observing, attending to emotions or suppressing emotions while viewing neutral, overall negative or interpersonal violence-related pictures) | Group comparison | Conflict Tactics Scale-2 (CTS2), IRI | Domestic violence offenders | 53 | 53 | 0 | 1 |
| 209a | Marsh & Cardinale (2014) | fMRI | Emotionally Evocative Statements Task (EEST) | Group comparison | PPI-Revised (PPI-R) | Healthy adults with high psychopathic traits | 33 | 13 | 20 | 4 |
| 209b | Marsh & Cardinale (2014) | fMRI | Emotionally Evocative Statements Task (EEST) | Correlational | PPI-Revised (PPI-R), RPQ | Healthy adults with high psychopathic traits | 33 | 13 | 20 | 3 |
| 210 | Marsh et al. (2013) | fMRI | Evaluating the severity of pain from pictures of severe, moderate or no pain while imagining either being in the picture or it happening to someone else | Group comparison | K-SADS-PL, PCL:YV | Psychopathic adolescents with CD or ODD | 35 | 23 | 12 | 2 |
| 211 | Marsh et al. (2011) | fMRI | Categorizing illegal and legal behaviors in a moral judgement implicit association task | Group comparison | K-SADS-PL, PCL:YV, APSD (ASPD) | Psychopathic adolescents with CD or ODD | 28 | 19 | 9 | 3 |
| 212 | Martinelli et al. (2021) | fMRI | Judging social intention of laughter (friendly/including laughter or hostile/excluding laughter) | Correlational | BPAQ physical aggression scale | na | 50 | 21 | 29 | 2 |
| 213 | Martinez-Horta et al. (2021) | MRI | VBM | Group comparison | Problem Behaviors Assessment for Huntington's Disease (PBA-s) irritability and aggression scores | Patients with early or mild Huntington's disease and clinically significant scores of aggression and/or irritability | 31 | 11 | 20 | 1 |
| 214 | Mathews et al. (2005) | fMRI | Counting Stroop task | Group comparison | na | Adolescents with disruptive behavior disorder with aggressive features | 38 | 28 | 10 | 2 |
| 215 | Matthies et al. (2012) | MRI | Manual tracing within MRreg program | Group comparison | LHA | Healthy participants with higher than sample median aggression scores (LHA > 5) | 18 | 0 | 18 | 1 |
| 216 | Maurer et al. (2019) | fMRI | Go/NoGo | Correlational | PCL:YV | Incarcerated offenders | 182 | 182 | 0 | 1 |
| 217a | McCloskey et al. (2016) | fMRI | Identifying emotional valence from emotional faces | Group comparison | LHA | IED and various personality disorder and Axis I disorder comorbidities | 40 | 24 | 16 | 2 |
| 217b | McCloskey et al. (2016) | fMRI | Identifying emotional valence from emotional faces | Correlational | LHA | IED and various personality disorder and Axis I disorder comorbidities | 40 | 24 | 16 | 1 |
| 218 | Meffert et al. (2013) | fMRI | Passively observing or empathizing videos showing neutral, loving, painful or excluding hand interactions or actually experiencing them | Group comparison | PCL-R, PPI | Psychopathic offenders | 44 | 44 | 0 | 6 |
| 219 | Meldrum et al. (2018) | fMRI | Go/NoGo | Correlational | CBCL | na | 85 | 59 | 26 | 1 |
| 220 | Menks et al. (2021) | fMRI | Implicit emotional face processing paradigm | Group comparison | K-SADS-PL, YPI | Adolescents with CD | 58 | 19 | 39 | 2 |
| 221 | Michalska et al. (2015) | MRI | VBM | Correlational | DISC-IV | ODD or CD | 111 | 58 | 53 | 1 |
| 222 | Michalska et al. (2016) | fMRI | Viewing pictures of intentional and unintentional harm | Correlational | DISC-IV, ICU | Children with some CD symptoms | 107 | 55 | 52 | 8 |
| 223 | Miedl et al. (2018) | fMRI | Viewing neutral and severely violent film clips | Activation study | na | na | 53 | 0 | 53 | 1 |

| | | | | | | | | | | |
|---|---|---|---|---|---|---|---|---|---|---|
| 224 | Mier et al. (2014) | fMRI | Answering questions associated with theory of mind, emotion recognition or facial processing concerning emotional faces | Group comparison | PCL-R, PPI | Psychopathic inmates | 29 | 29 | 0 | 4 |
| 225 | Miskovich et al. (2018) | MRI | Freesurfer | Correlational | PCL-R | Adult inmates | 716 | 716 | 0 | 1 |
| 226a | Moeller et al. (2014) | fMRI | Stroop task | Group comparison | LHA, STAXI | IED | 28 | 28 | 0 | 3 |
| 226b | Moeller et al. (2014) | fMRI | Stroop task | Correlational | LHA, STAXI | IED | 49 | 49 | 0 | 1 |
| 227 | Mohammadi et al. (2020) | MRI | VBM | Correlational | IRI, Fragebogen zur Erfassung von Agressionsfaktoren (FAF), Cook-Medley Hostility Scale (Ho scale), Die Skalen zum Erleben von Emotionen (SEE) | Young adults with video game addiction | 29 | 29 | 0 | 3 |
| 228a | Molenberghs et al. (2014) | fMRI | Seemingly rewarding (money) or punishing (electric shock) other participants in an in-group or an out-group according to whether they answered trivia questions correctly | Correlational | SRP-III | na | 48 | 24 | 24 | 1 |
| 228b | Molenberghs et al. (2014) | fMRI | Seemingly rewarding (money) or punishing (electric shock) other participants in an in-group or an out-group according to whether they answered trivia questions correctly | Activation study | SRP-III | na | 48 | 24 | 24 | 1 |
| 229 | Molenberghs et al. (2016) | fMRI | Viewing videos of ingroup and outgroup members either interacting or causing intentional harm to each other | Activation study | na | na | 48 | 24 | 24 | 1 |
| 230 | Montag et al. (2012) | fMRI | Viewing generally unpleasant, pleasant or neutral pictures or violent screenshots from Counterstrike | Activation study | ANPS anger scale | na | 40 | 40 | 0 | 1 |
| 231 | Morandotti et al. (2013) | MRI | VBM | Correlational | BDHI | BPD | 11 | | | 1 |
| 232 | Murray et al. (2017) | fMRI | Card-guessing game | Correlational | Self-Report of Antisocial Behavior Questionnaire, APSD, Structured Clinical Interview for DSM-IV for Axis II personality disorders | na | 144 | 144 | 0 | 6 |
| 233 | Müller et al. (2008) | MRI | VBM | Group comparison | PCL-R | Psychopathic patients | 34 | 34 | 0 | 1 |
| 234 | Müller et al. (2008a) | fMRI | Positive, neutral and negative affective pictures and a Simon task | Group comparison | PCL-R | Psychopathic offenders with ASPD | 22 | 22 | 0 | 2 |
| 235 | Müller et al. (2003) | fMRI | Positive, neutral and negative emotional images | Group comparison | PCL-R | Psychopathic criminals | 12 | 12 | 0 | 4 |
| 236a | Naaijen et al. (2020) | MRI | Freesurfer | Group comparison | CBCL, K-SADS, RPQ, ICU | CD and/or ODD and/or scored above the clinical cut-off for aggressive behavior and/or rule-breaking behavior in CBCL | 254 | 185 | 69 | 1 |
| 236b | Naaijen et al. (2020) | MRI | Freesurfer | Correlational | CBCL, K-SADS, RPQ, ICU | CD and/or ODD and/or scored above the clinical cut-off for aggressive behavior and/or rule-breaking behavior in CBCL | 254 | 185 | 69 | 2 |
| 237 | Nakano et al. (2006) | SPECT | 99mTc-ECD | Correlational | Semi-structured interview with family members with a modified version of the Neuropsychiatric Inventory (NPI) | Frontotemporal dementia patients, most with histories of some antisocial behavior | 98 | 51 | 47 | 1 |
| 238a | Narayan et al. (2007) | MRI | Manual tracing and cortical pattern matching | Group comparison | clinical interview for DSM-IV personality disorders, Gunn and Robertson scale | ASPD and a history of violence | 29 | 29 | 0 | 1 |
| 238b | Narayan et al. (2007) | MRI | Manual tracing and cortical pattern matching | Group comparison | clinical interview for DSM-IV axis I disorders, Gunn and Robertson scale for violence | SCZ and a history of violence | 27 | 27 | 0 | 1 |
| 239 | Navalpotro-Gomez et al. (2019) | SPECT | 123I-2β-carbometoxy-3β-(4-iodophenyl)-N-(3-fluoropropyl) nortropane (123I-FP-CIT) | Correlational | QUIP-RS | Nondemented PD patients with ICDs | 16 | 14 | 2 | 2 |
| 240a | New et al. (2007) | MRI | Manual tracing | Correlational | BDHI, BIS-7b | na | 24 | 15 | 9 | 1 |
| 240b | New et al. (2007) | MRI | Manual tracing | Correlational | BDHI, BIS-7b | BPD, IED and impulsive aggression | 26 | 17 | 9 | 1 |
| 241a | New et al. (2009) | PET | 18fluorodeoxyglucose [18FDG] | Group comparison | BIS-II, STAXI, OAS Modified (OAS-M), BDHI, BPAQ | BPD and IED | 74 | 40 | 34 | 2 |
| 241b | New et al. (2009) | PET | 18fluorodeoxyglucose [18FDG] | Correlational | BIS-II, STAXI, OAS Modified (OAS-M), BDHI, BPAQ | na | 36 | 18 | 18 | 2 |
| 241c | New et al. (2009) | PET | 18fluorodeoxyglucose [18FDG] | Correlational | BIS-II, STAXI, OAS Modified (OAS-M), BDHI, BPAQ | BPD and IED | 38 | 22 | 16 | 1 |
| 242 | Oberlin et al. (2012) | fMRI | Smelling either odors of preferred alcoholic drinks, appetitive control odors or non-appetitive odors while saline (or 6 % alcohol) was infused intravenously | Correlational | SSAGA | Heavy drinkers | 30 | | | 4 |
| 243 | Oder et al. (1992) | SPECT | 99mTc-HMPAO | Correlational | the Giessen test | Patients with severe closed head injury coming up for a follow up | 36 | 31 | 5 | 2 |
| 244 | Oquendo et al. (2005) | PET | [18F]FDG | Correlational | BDHI, Brown Goodwin Lifetime Aggression Scale (BG), BIS | Patients diagnosed with MDD and comorbid BPD | 19 | 0 | 19 | 2 |
| 245 | Osumi et al. (2012) | fMRI | Receiving unfair offers of monetary distribution while being either able or unable to punish the offerer | Correlational | primary and secondary psychopathy scales (PSPS) | na | 23 | 23 | 0 | 2 |
| 246 | Overgaauw et al. (2019) | fMRI | Judging trustworthiness of faces with feedback | Group comparison | PPI Short Form (PPI-SF) | Healhty participants with psychopathic traits | 42 | 0 | 42 | 1 |
| 247 | Pagliaccio et al. (2017) | fMRI | Global-local selective attention task | Group comparison | Affective Reactivity Index (ARI) | Youths with disruptive mood dysregulation disorder | 58 | 28 | 30 | 2 |
| 248 | Pardini & Phillips (2010) | fMRI | Faces showing emotions at mild and prototypic intensities | Group comparison | The Self-Report of Psychopathy-III (SRP-III) | Chronically violent men | 42 | 42 | 0 | 3 |
| 249 | Pardini et al. (2014) | MRI | FMRIB's Integrated Registration and Segmentation Tool (FIRST) in the FMRIB Software Library (FSL) version 4.1 | Correlational | Self-Report of Delinquency, Violence History Questionnaire, Self-Report of Psychopathy-III - short form(SRP-II-SF), Adult Self-Report (ASR), Aggression Questionnaire - short form (AQ-SF), Impulsive-Premeditated Aggression Scale, Teacher Report Form (TRF), Child Psychopathy Scale-revised, RPQ | Chronically violent men | 56 | 56 | 0 | 3 |
| 250 | Parsey et al. (2002) | PET | [carbonyl-C-11]WAY-100635 | Correlational | Brown Goodwin Aggression History Scale | na | 25 | 13 | 12 | 1 |
| 251 | Passamonti et al. (2010) | fMRI | Emotional faces | Group comparison | YPI | Adolescents with CD | 62 | 62 | 0 | 2 |
| 252a | Pawliczek et al. (2013) | fMRI | Frustration task; solvable and unsolvable anagrams with monetary rewards | Group comparison | BPAQ, PPI-R, LHA, Freiburger Aggression Inventory (FAI) | Healthy individuals with high trait aggression | 39 | 39 | 0 | 2 |
| 252b | Pawliczek et al. (2013) | fMRI | Frustration task; solvable and unsolvable anagrams with monetary rewards | Correlational | BPAQ, PPI-R, LHA, Freiburger Aggression Inventory (FAI) | Healthy individuals with high trait aggression | 21 | 21 | 0 | 1 |
| 253 | Pawliczek et al. (2013a) | fMRI | SST with emotional faces | Group comparison | BPAQ, BIS, PPI-R | Healthy individuals with high trait aggression | 33 | 33 | 0 | 3 |
| 254 | Payer et al. (2012) | fMRI | Labeling or matching emotional faces | Correlational | BPAQ, STAXI | na | 10 | 4 | 6 | 1 |
| 255 | Payer et al. (2011) | fMRI | Labeling or matching emotional faces | Correlational | BPAQ, STAXI | na | 37 | 19 | 18 | 1 |
| 256 | Paz-Alonso et al. (2020) | fMRI | Simplified Iowa Gambling Task | Group comparison | na | PD with ICDs but without dementia | 35 | 31 | 4 | 1 |
| 257 | Pellicano et al. (2015) | MRI | Freesurfer | Group comparison | QUIP | Non-demented patients with PD with ICDs | 36 | 30 | 6 | 1 |
| 258 | Pera-Guardiola et al. (2016) | MRI | VBM | Group comparison | PCL-R, BIS | Psychopathic criminals | 39 | 39 | 0 | 1 |
| 259 | Perez-Rodriguez et al. (2012) | PET | 18fluoro-deoxyglucose | Group comparison | BIS-II, OAS-M, STAXI, BDHI, BPAQ | BPD and IED | 40 | 40 | 0 | 1 |
| 260 | Perino et al. (2019) | fMRI | CP | Correlational | Illinois Bully Scale (IBS) | Adolescents with histories of conduct problems and recent delinquent behaviour | 24 | 12 | 12 | 2 |

| | | | | | | | | | | |
|---|---|---|---|---|---|---|---|---|---|---|
| 261 | Pietrini et al. (2000) | PET | [15O]H2O | Activation study | na | na | 15 | 8 | 7 | 5 |
| 262 | Porges & Decety (2013) | fMRI | Watching videos of either mixed martial arts or capoeira as control | Activation study | na | na | 49 | 49 | 0 | 2 |
| 263a | Prehn et al. (2013) | fMRI | Choosing between high or low risk monetary rewards | Group comparison | International Personality Disorder Examination (IPDE), PCL-R, Temperament and Character Inventory (TCI), Questionnaire for Factors of Aggressiveness (FAF) | Emotionally hypo-reactive criminals with ASPD | 24 | 24 | 0 | 3 |
| 263b | Prehn et al. (2013) | fMRI | Choosing between high or low risk monetary rewards | Group comparison | International Personality Disorder Examination (IPDE), PCL-R, Temperament and Character Inventory (TCI), Questionnaire for Factors of Aggressiveness (FAF) | Emotionally hyper-reactive criminals with ASPD | 25 | 25 | 0 | 1 |
| 263c | Prehn et al. (2013) | fMRI | Choosing between high or low risk monetary rewards | Correlational | International Personality Disorder Examination (IPDE), PCL-R, Temperament and Character Inventory (TCI), Questionnaire for Factors of Aggressiveness (FAF) | Emotionally hypo-reactive criminals with ASPD and emotionally hyper-reactive criminals with ASPD and BPD | 36 | 36 | 0 | 1 |
| 264a | Prehn et al. (2013a) | fMRI | Verbal n-back task with emotional pictures in the background | Group comparison | PCL-R, International Personality Disorder Examination, Temperament and Character Inventory (TCI), Questionnaire for Factors of Aggressiveness (FAF) | Violent criminal offenders with ASPD and BPD | 32 | 32 | 0 | 4 |
| 264b | Prehn et al. (2013a) | fMRI | Verbal n-back task with emotional pictures in the background | Correlational | PCL-R, International Personality Disorder Examination, Tempoeament and Character Inventory (TCI), Questionnaire for Factors of Aggressiveness (FAF) | Violent criminal offenders with ASPD and BPD | 15 | 15 | 0 | 1 |
| 265 | Premi et al. (2016) | SPECT | 123I-FP-CIT | Group comparison | QUIP-RS | PD and ICD | 84 | 54 | 30 | 1 |
| 266a | Pujara et al. (2013) | MRI | Freesurfer | Correlational | PCL-R | Psychopatic inmates | 18 | 18 | 0 | 1 |
| 266b | Pujara et al. (2013) | fMRI | Passive gain or loss of money (slot machine) | Correlational | PCL-R | Psychopatic inmates | 18 | 18 | 0 | 2 |
| 267 | Pujol et al. (2012) | fMRI | Moral dilemma task and a Stroop task | Group comparison | PCL-R | Psychopathic criminals | 44 | 44 | 0 | 2 |
| 268 | Puri et al. (2008) | MRI | VBM (FSL-VBM) | Group comparison | na | SCZ patients with histories of violent offending | 26 | 22 | 4 | 1 |
| 269 | Qiao et al. (2016) | fMRI | Passively viewing negative or neutral emotional images | Group comparison | modified OAS (MOAS), BIS-11 | Violent adolescents | 39 | 39 | 0 | 2 |
| 270 | Qiao et al. (2012) | fMRI | Go/no-go | Group comparison | modified OAS (MOAS), BIS-11 | Violent adolescents | 39 | 39 | 0 | 4 |
| 271 | Quan et al. (2019) | MRI | VBM | Correlational | Word Sentence Association Paradigm - Hostility (WSAP-H), Attitudes toward Violence Scale (AVS) | na | 176 | 83 | 93 | 3 |
| 272 | Raine et al. (1997) | PET | FDG | Group comparison | na | Murderers pleading not guilty by reason of insanity | 82 | 78 | 4 | 2 |
| 273 | Raine et al. (1994) | PET | FDG | Group comparison | na | Murderers pleading not guilty by reason of insanity | 44 | 40 | 4 | 1 |
| 274a | Raine et al. (2000) | MRI | Semiautomatic segmentation with CAMRA S200 ALLEGRO software | Group comparison | na | ASPD | 47 | 47 | 0 | 1 |
| 274b | Raine et al. (2000) | MRI | Semiautomatic segmentation with CAMRA S200 ALLEGRO software | Group comparison | na | ASPD | 55 | 55 | 0 | 1 |
| 275a | Raine et al. (1998) | PET | FDG | Group comparison | na | Predatory murderers pleading not guilty by reason of insanity | 56 | | | 1 |
| 275b | Raine et al. (1998) | PET | FDG | Group comparison | na | Affective murderers pleading not guilty by reason of insanity | 50 | | | 1 |
| 276a | Raine et al. (2011) | MRI | Automated segmentation algorithm, LONI Pipeline Processing Environment | Group comparison | Structured clinical interview for DSM-IV, Self-Report Crime Checklist | ASPD | 42 | 42 | 0 | 1 |
| 276b | Raine et al. (2011) | MRI | Automated segmentation algorithm, LONI Pipeline Processing Environment | Group comparison | Structured clinical interview for DSM-IV, Self-Report Crime Checklist | ASPD | 48 | 48 | 0 | 1 |
| 276c | Raine et al. (2011) | MRI | Automated segmentation algorithm, LONI Pipeline Processing Environment | Correlational | Structured clinical interview for DSM-IV, Self-Report Crime Checklist | na | 12 | 0 | 12 | 2 |
| 277 | Rao et al. (2010) | fMRI | Balloon Analogue Risk Task (BART) | Group comparison | na | PD patients with ICDs | 18 | 15 | 3 | 1 |
| 278 | Raschle et al. (2019) | fMRI | Passively viewing or regulating emotions via reappraisal while viewing neutral or negative pictures | Group comparison | KSADS-PL, YPI, CBCL | CD and/or ODD | 59 | 0 | 59 | 1 |
| 279 | Raschle et al. (2018) | MRI | VBM | Correlational | CBCL, ICU, YPI | na | 81 | 81 | 0 | 1 |
| 280 | Regenbogen et al. (2010) | fMRI | Watching virtual (from a FPS game) or acted scenes of nonviolent or violent content | Activation study | na | na | 11 | 11 | 0 | 2 |
| 281 | Reniers et al. (2012) | fMRI | Moral and non-moral decision making task | Correlational | Levenson Self-Report Psychopathy Scale (LSRP) | na | 24 | 24 | 0 | 1 |
| 282a | Repple et al. (2018) | fMRI | Modified TAGG (with monetary reward/punishment) | Correlational | BPAQ, STAXI | na | 22 | 22 | 0 | 2 |
| 282b | Repple et al. (2018) | fMRI | Modified TAGG (with monetary reward/punishment) | Correlational | BPAQ, STAXI | na | 20 | 0 | 20 | 1 |
| 283a | Repple et al. (2017) | fMRI | Modified TAGG (with monetary reward/punishment) | Correlational | BPAQ, STAXI, Positive and Negative Affect Schedule (PANAS), emotional self rating (ESR) | na | 29 | 29 | 0 | 1 |
| 283b | Repple et al. (2017) | fMRI | Modified TAGG (with monetary reward/punishment) | Activation study | BPAQ, STAXI, Positive and Negative Affect Schedule (PANAS), emotional self rating (ESR) | na | 29 | 29 | 0 | 1 |
| 284 | Rijsdijsk et al. (2010) | MRI | VBM | Correlational | Strenghts and Difficulties Questionnaire, APSD callous-unemotional -scale | High conduct problems and core psychopathic features | 125 | 125 | 0 | 1 |
| 285 | Rilling et al. (2007) | fMRI | Prisoner's dilemma game with monetary rewards | Correlational | PPI, Levenson Primary and Secondary Psychopathy Scales | na | 30 | 15 | 15 | 16 |
| 286 | Rodman et al. (2016) | fMRI | Eriksen flanker task with a go/no go manipulation | Correlational | PCL-R, ESI | Inmates | 46 | 46 | 0 | 3 |
| 287a | Rosell et al. (2010) | PET | [11C]MDL100907 | Group comparison | OAS-Modified (OAS-M), BPAQ, BIS-11 | Patients with at least one DSM-IV personality disorder and IED with current physical aggression | 29 | 22 | 7 | 1 |
| 287b | Rosell et al. (2010) | PET | [11C]MDL100907 | Correlational | OAS-Modified (OAS-M), BPAQ, BIS-11 | Patients with at least one DSM-IV personality disorder and IED with current physical aggression | 29 | 22 | 7 | 2 |
| 287c | Rosell et al. (2010) | PET | [11C]MDL100907 | Group comparison | OAS-Modified (OAS-M), BPAQ, BIS-11 | At least one DSM-IV personality disorder and IED | 39 | 25 | 14 | 3 |
| 288 | Rosenthal-Von Der Pütten et al. (2014) | fMRI | Seeing violent human-human interaction (as well as human-robot and human-box interaction) + a low-level control visual search task | Activation study | na | na | 14 | 5 | 9 | 1 |
| 289 | Rubia et al. (2010) | fMRI | Meiran switch task | Group comparison | na | Noncomorbid CD | 34 | 34 | 0 | 1 |
| 290 | Rubia et al. (2009) | fMRI | Simon task including a congruent oddball | Group comparison | SDQ | Noncomorbid CD | 33 | 33 | 0 | 2 |
| 291 | Rubia et al. (2008) | fMRI | Visual tracking stop task (inhibiting motor response on stop signals) | Group comparison | SDQ | CD, ODD | 33 | 33 | 0 | 2 |
| 292 | Rubia et al. (2009a) | fMRI | Rewarded continous performance task | Group comparison | SDQ | Noncomorbid CD | 30 | 30 | 0 | 2 |
| 293a | Rylands et al. (2012) | PET | 11C-DASB | Group comparison | PPI-R, Impulsiveness-Venturesomeness-Empathy questionnaire (IVE-7), Expression Aggression Questionnaire (EXPAGG), BIS-11, STAXI, Novaco Anger Scale and Provocation Inventory (NAS-PI), BDHI | Violent offenders with ASPD, BPD, or both | 22 | 22 | 0 | 1 |

| | | | | | | | | | | |
|---|---|---|---|---|---|---|---|---|---|---|
| 293b | Rylands et al. (2012) | PET | 11C-DASB | Correlational | PPI-R, Impulsiveness-Venturesomeness-Empathy questionnaire (IVE-7), Expression Aggression Questionnaire (EXPAGG), BIS-11, STAXI, Novaco Anger Scale and Provocation Inventory (NAS-PI), BDHI | Violent offenders with ASPD, BPD, or both | 22 | 22 | 0 | 6 |
| 294 | Sadeh et al. (2013) | fMRI | Emotion-word Stroop task | Correlational | NEO-Five Factor Inventory (NEO-FFI) from which impulsive-antisociality (low agreeableness and conscientiousness) and fearless-dominance (low neuroticism, high extraversion and low agreeableness) dimensions were computed | na | 49 | 19 | 30 | 5 |
| 295 | Sajous-Turner et al. (2020) | MRI | VBM | Group comparison | PCL-R | Offenders convicted for homicide or have tried to commit homicide | 678 | 678 | 0 | 1 |
| 296a | Sakai et al. (2017) | fMRI | The altruism-antisocial game (AlAn's game) | Group comparison | Inventory of Callous Unemotional traits | Adolescents with conduct problems and low prosocial emotions | 45 | 45 | 0 | 1 |
| 296b | Sakai et al. (2017) | fMRI | The altruism-antisocial game (AlAn's game) | Group comparison | Inventory of Callous Unemotional traits | Adolescents with conduct problems and low prosocial emotions | 42 | 42 | 0 | 1 |
| 296c | Sakai et al. (2017) | fMRI | The altruism-antisocial game (AlAn's game) | Group comparison | Inventory of Callous Unemotional traits | Adolescents with conduct problems and high ICU score | 42 | 42 | 0 | 1 |
| 297 | Sala et al. (2011) | MRI | Manual tracing with Brains2 semiautomated software | Correlational | BDHI, BIS-11 | BPD | 15 | 4 | 11 | 2 |
| 298 | Sarkar et al. (2015) | MRI | Freesurfer | Group comparison | SDQ, APSD | Adolescents with CD | 40 | 40 | 0 | 1 |
| 299a | Schienle et al. (2017) | fMRI | Static or approaching pictures of neutral faces | Group comparison | PCL-R | Incarcerated violent offenders | 35 | 35 | 0 | 3 |
| 299b | Schienle et al. (2017) | fMRI | Static or approaching pictures of neutral faces | Correlational | PCL-R | Incarcerated violent offenders | 17 | 17 | 0 | 1 |
| 300a | Schiffer et al. (2017) | MRI | VBM | Group comparison | Screening Scale for Pedophilic Interest, 2nd version (SSPI-2) | Pedophilic sexual offenders | 118 | 118 | 0 | 1 |
| 300b | Schiffer et al. (2017) | MRI | VBM | Correlational | Screening Scale for Pedophilic Interest, 2nd version (SSPI-2) | Pedophilic sexual offenders | 58 | 58 | 0 | 1 |
| 300c | Schiffer et al. (2017) | MRI | VBM | Group comparison | Screening Scale for Pedophilic Interest, 2nd version (SSPI-2) | Pedophilic sexual offenders | 159 | 159 | 0 | 1 |
| 301a | Schiffer et al. (2013) | MRI | VBM | Group comparison | Assessment of life history of aggression interview | CD prior to age 15 | 52 | 52 | 0 | 2 |
| 301b | Schiffer et al. (2013) | MRI | VBM | Group comparison | Assessment of life history of aggression interview | SCZ patients with CD prior to age 15 | 50 | 50 | 0 | 2 |
| 301c | Schiffer et al. (2013) | MRI | VBM | Correlational | Assessment of life history of aggression interview | CD prior to age 15 | 52 | 52 | 0 | 1 |
| 301d | Schiffer et al. (2013) | MRI | VBM | Correlational | Assessment of life history of aggression interview | SCZ patients with CD prior to age 15 | 50 | 50 | 0 | 1 |
| 302a | Schiffer et al. (2011) | MRI | VBM | Group comparison | PCL-SV, Life History of Aggression assessment, BIS-11, perseverative errors on the Wisconsin Card Sorting Test + commission errors on a Go/noGo task | Violent offenders | 51 | 51 | 0 | 2 |
| 302b | Schiffer et al. (2011) | MRI | VBM | Correlational | PCL-SV, Life History of Aggression assessment, BIS-11, perseverative errors on the Wisconsin Card Sorting Test + commission errors on a Go/noGo task | Violent offenders | 51 | 51 | 0 | 4 |
| 303a | Schiffer et al. (2014) | fMRI | Stroop task | Group comparison | PCL:SV, BIS-11 | Incarcerated violent offenders | 44 | 44 | 0 | 3 |
| 303b | Schiffer et al. (2014) | fMRI | Stroop task | Correlational | PCL:SV, BIS-11 | Incarcerated violent offenders | 44 | 44 | 0 | 2 |
| 304a | Schiffer et al. (2017a) | fMRI | Reading the Mind in the Eyes task (mental state decoding from visual stimuli) | Group comparison | PCL:SV | Violent offenders with SCZ and CD | 31 | 31 | 0 | 1 |
| 304b | Schiffer et al. (2017a) | fMRI | Reading the Mind in the Eyes task (mental state decoding from visual stimuli) | Group comparison | PCL:SV | Violent offenders with SCZ and without CD | 34 | 34 | 0 | 1 |
| 304c | Schiffer et al. (2017a) | fMRI | Reading the Mind in the Eyes task (mental state decoding from visual stimuli) | Group comparison | PCL:SV | Violent offenders with SCZ and CD | 29 | 29 | 0 | 1 |
| 304d | Schiffer et al. (2017a) | fMRI | Reading the Mind in the Eyes task | Correlational | PCL:SV | Violent offenders with SCZ and CD | 47 | 47 | 0 | 2 |
| 305 | Schlüter et al. (2013) | PET | 6-[18F]-fluoro-L-DOPA (FDOPA) | Correlational | ratio of aggressive responses to all responses in point-subtraction aggression paradigm | na | 21 | 21 | 0 | 2 |
| 306 | Schneider et al. (2000) | fMRI | Conditioning task | Group comparison | PCL-R | Psychopathic patients with ASPD | 24 | 24 | 0 | 1 |
| 307 | Schoretsanitis et al. (2019) | MRI | VBM | Correlational | Modified OAS (MOAS) | SCZ spectrum disorders | 84 | 54 | 30 | 1 |
| 308 | Schultz et al. (2016) | fMRI | Pavlovian fear conditioning with electric shocks and two fractal images as conditional stimuli | Group comparison | PCL-R, Welsh Anxiety Scale (WAS) | Psychopathic prisoners | 50 | 50 | 0 | 1 |
| 309 | Schulz et al. (2013) | PET | Fluorodeoxyglucuse-F18 | Correlational | Zanarini Rating Scale for Borderline Personality Disorder (ZAN-BPD), BDHI | Adult women with BPD | 14 | 0 | 14 | 4 |
| 310a | Schwenck et al. (2017) | fMRI | Monetary gambling task | Group comparison | Diagnostic interview for mental disorders in children and adolescents (Kinder-DIPS), Parent rating scale for CD (FBB-SSV), ICU, CBCL | Adolescents with conduct problems | 43 | 43 | 0 | 2 |
| 310b | Schwenck et al. (2017) | fMRI | Monetary gambling task | Correlational | Diagnostic interview for mental disorders in children and adolescents (Kinder-DIPS), Parent rating scale for CD (FBB-SSV), ICU, CBCL | Adolescents with conduct problems | 43 | 43 | 0 | 1 |
| 311 | Seara-Cardoso et al. (2016) | fMRI | Assessing personal guilt on everyday moral scenarios which either end in harm-to-other or harm-to-self outcomes | Correlational | Self-Report Psychopathy Scale Short-Form (SRP-SF) | na | 28 | 28 | 0 | 1 |
| 312 | Seara-Cardoso et al. (2016a) | fMRI | Rating own affective state while observing images of emotional faces | Correlational | Self-Report Psychopathy Scale Short-Form (SRP-SF) | na | 30 | 30 | 0 | 1 |
| 313 | Seara-Cardoso et al. (2015) | fMRI | Observing pictures of other person's hand or foot in painful or nonpainful situations | Correlational | Self-Report Psychopathy Scale Short-Form (SRP-SF) | na | 46 | 46 | 0 | 2 |
| 314a | Sebastian et al. (2016) | MRI | VBM | Group comparison | ICU | Adolescents with conduct problems | 89 | 89 | 0 | 1 |
| 314b | Sebastian et al. (2016) | MRI | VBM | Group comparison | ICU | Conduct problems and high callous-unemotional traits | 60 | 60 | 0 | 1 |
| 314c | Sebastian et al. (2016) | MRI | VBM | Group comparison | ICU | Conduct problems and low callous-unemotional traits | 60 | 60 | 0 | 1 |
| 315a | Sebastian et al. (2014) | fMRI | Observing fearful or calm faces or eyes | Group comparison | ICU | Current conduct problems and low callous-unemotional traits | 34 | 34 | 0 | 3 |
| 315b | Sebastian et al. (2014) | fMRI | Observing fearful or calm faces or eyes | Group comparison | ICU | Current conduct problems and high callous-unemotional traits | 34 | 34 | 0 | 1 |
| 316a | Sebastian et al. (2012) | fMRI | Choosing correct ending to a cartoon concerning either affective theory of mind, cognitive theory of mind or simple physical causality | Group comparison | ICU, Child and Adolescent Symptom Inventory - conduct disorder (CASI-CD) scale | Children with conduct problems | 47 | 47 | 0 | 1 |
| 316b | Sebastian et al. (2012) | fMRI | Choosing correct ending to a cartoon concerning either affective theory of mind, cognitive theory of mind or simple physical causality | Correlational | ICU, Child and Adolescent Symptom Inventory - conduct disorder (CASI-CD) scale | Children with conduct problems | 47 | 47 | 0 | 1 |
| 317 | Sebastian et al. (2021) | fMRI | Emotional Simon task with stimuli of fearful, angry, calm and scrambled expressions | Group comparison | ICU, Child and Adolescent Symptom Inventory-4R (CASI-CD) | Children with conduct problems and low callous-unemotional traits | 35 | 35 | 0 | 5 |

| | | | | | | | | | | |
|---|---|---|---|---|---|---|---|---|---|---|
| 318 | Seidenwurm et al. (1997) | PET | fludeoxyglucose F18 (FDG) | Group comparison | na | Severely violent criminals with various psychiatric disorders | 16 | 6 | 10 | 1 |
| 319 | Sekine et al. (2006) | PET | Trans-1,2,3,5,6,10-beta-hexahydro-6-[4-(methylthio)phenyl]pyrrolo-[2,1-a]isoquinoline ([11C](+)McN-5652) | Correlational | BPAQ | Currently abstinent former methamphetamine abusers | 12 | 7 | 5 | 1 |
| 320 | Seleem et al. (2020) | MRI | Freesurfer | Group comparison | na | CD | 33 | 13 | 20 | 3 |
| 321 | Seo et al. (2016) | fMRI | Script guided imagery task with stress, alcohol cue and neutral-relaxing conditions | Correlational | Difficulties in emotion regulation scale (DERS) difficulties in impulse control (IMPULSE) scale | Alcohol dependent patients who had been 4 to 8 weeks abstinent | 37 | 29 | 8 | 1 |
| 322a | Seok & Cheong (2020) | fMRI | Observing anger eliciting or neutral film clips | Group comparison | BPAQ, LHA, STAXI-2 | IED | 30 | 30 | 0 | 1 |
| 322b | Seok & Cheong (2020) | fMRI | Observing anger eliciting or neutral film clips | Correlational | BPAQ, LHA, STAXI-2 | IED | 30 | 30 | 0 | 1 |
| 322c | Seok & Cheong (2020) | MRI | VBM | Group comparison | BPAQ, LHA, STAXI-2 | IED | 30 | 30 | 0 | 1 |
| 322d | Seok & Cheong (2020) | MRI | VBM | Correlational | BPAQ, LHA, STAXI-2 | IED | 15 | 15 | 0 | 1 |
| 323a | Sethi et al. (2018) | fMRI | Matching emotional or neutral faces | Group comparison | Self-Report Psychopathy-Short Form (SRP-SF), State-Trait Anxiety Inventory | Primarily psychopathic | 132 | 61 | 71 | 1 |
| 323b | Sethi et al. (2018) | fMRI | Matching emotional or neutral faces | Group comparison | Self-Report Psychopathy-Short Form (SRP-SF), State-Trait Anxiety Inventory | Secondarily psychopathic | 182 | 78 | 104 | 1 |
| 324a | Shane & Groat (2018) | fMRI | Increasing or decreasing emotional response or just watching when shown neutral or negatively valent pictures | Group comparison | PCL-R | Psychopathic criminals | 38 | 34 | 4 | 3 |
| 324b | Shane & Groat (2018) | fMRI | Increasing or decreasing emotional response or just watching when shown neutral or negatively valent pictures | Group comparison | PCL-R | Criminals with PCL-R score 21-29 | 52 | 42 | 10 | 3 |
| 324c | Shane & Groat (2018) | fMRI | Increasing or decreasing emotional response or just watching when shown neutral or negatively valent pictures | Group comparison | PCL-R | Psychopathic criminals | 44 | 36 | 8 | 2 |
| 324d | Shane & Groat (2018) | fMRI | Increasing or decreasing emotional response or just watching when shown neutral or negatively valent pictures | Group comparison | PCL-R | Criminals with high PCL-R score | 67 | 56 | 11 | 2 |
| 324e | Shane & Groat (2018) | fMRI | Increasing or decreasing emotional response or just watching when shown neutral or negatively valent pictures | Correlational | PCL-R | Criminals with high PCL-R score | 67 | 56 | 11 | 1 |
| 325 | Shao & Lee (2017) | fMRI | Lying or telling the truth about familiarity of shown faces and a visuo-spatial response control task | Group comparison | PPI-R | Healthy individuals with psychopathic traits | 52 | 26 | 26 | 7 |
| 326 | Shapiro et al. (2000) | SPECT | Technetium-99m-pertechnetate | Correlational | Cook-Medley Hostility Scale | Healthy individuals scoring in the highest quartile of hostility in a larger sample (n=48) | 10 | 10 | 0 | 1 |
| 327 | Sharp et al. (2011) | fMRI | Playing a monetary trust game with a mean, kind or neutral (unknown) player, who was shown to be the most mean/kind peer nominated by the subject | Group comparison | youth self report (YSR), CBCL, peer nomination instrument | Externalizing and aggressive boys | 20 | 20 | 0 | 1 |
| 328 | Sheng et al. (2010) | fMRI | Producing speech according to the cued intonation condition (happy/sad/neutral/question) + a rest condition | Correlational | PPI-R | na | 19 | | | 2 |
| 329 | Sitaram et al. (2014) | fMRI | Training to up-regulate with neurofeedback | Correlational | PCL:SV, Levenson Self-Report Psychopathy Scale (LSRP) | Psychopathic sexual offenders | 4 | | | 1 |
| 330a | Skibsted et al. (2017) | fMRI | PSAP | Correlational | BPAQ | na | 19 | 8 | 11 | 1 |
| 330b | Skibsted et al. (2017) | fMRI | PSAP | Activation study | BPAQ | na | 19 | 8 | 11 | 4 |
| 331a | Soderstrom et al. (2002) | SPECT | [99mTC]-d, l-HMPAO | Group comparison | PCL-R | Individuals charged with severe crime and with high PCL-R factor 1 scores | 32 | 29 | 3 | 1 |
| 331b | Soderstrom et al. (2002) | SPECT | [99mTC]-d, l-HMPAO | Group comparison | PCL-R | Individuals charged with severe crime and with high PCL-R factor 2 scores | 32 | 29 | 3 | 1 |
| 331c | Soderstrom et al. (2002) | SPECT | [99mTC]-d, l-HMPAO | Group comparison | PCL-R | Individuals charged with severe crime and with high PCL-R factor 1 scores (3-factor model) | 32 | 29 | 3 | 1 |
| 331d | Soderstrom et al. (2002) | SPECT | [99mTC]-d, l-HMPAO | Group comparison | PCL-R | Individuals charged with severe crime and with high PCL-R factor 2 scores (3-factor model) | 32 | 29 | 3 | 1 |
| 331e | Soderstrom et al. (2002) | SPECT | [99mTC]-d, l-HMPAO | Group comparison | PCL-R | Individuals charged with severe crime and with high PCL-R factor 3 scores (3-factor model) | 32 | 29 | 3 | 1 |
| 332 | Soderstrom et al. (2000) | SPECT | [99mTC]-d, l-HMPAO | Group comparison | na | Perpetrators of impulsive violent crimes | 32 | 28 | 4 | 4 |
| 333a | Soloff et al. (2014) | MRI | VBM | Correlational | international personality disorders examination (IPDE), the diagnostic interview for borderline patients (DIB), Columbia Suicide History Form and Lethality Rating Scale (LRS), LHA, BIS | BPD | 35 | 5 | 30 | 2 |
| 333b | Soloff et al. (2014) | MRI | VBM | Correlational | international personality disorders examination (IPDE), the diagnostic interview for borderline patients (DIB), Columbia Suicide History Form and Lethality Rating Scale (LRS), LHA, BIS | BPD | 16 | 5 | 11 | 3 |
| 334a | Soloff et al. (2017) | fMRI | Go No-Go task with emotional faces | Correlational | international personality disorders examination (IPDE), the diagnostic interview for borderline patients - revised (DIB-R), BIS-11, LHA | BPD | 31 | 0 | 31 | 2 |
| 334b | Soloff et al. (2017) | fMRI | Go No-Go task with emotional faces | Correlational | international personality disorders examination (IPDE), the diagnostic interview for borderline patients - revised (DIB-R), BIS-11, LHA | na | 25 | 0 | 25 | 2 |
| 335a | Soloff et al. (2014a) | PET | [18F]altanserin | Correlational | international personality disorders examination (IPDE), the diagnostic interview for borderline patients - revised (DIB-R), BIS-11, LHA | BPD and impulsivity | 33 | 13 | 20 | 1 |
| 335b | Soloff et al. (2014a) | PET | [18F]altanserin | Correlational | international personality disorders examination (IPDE), the diagnostic interview for borderline patients - revised (DIB-R), BIS-11, LHA | BPD and impulsivity | 32 | 0 | 32 | 2 |
| 336 | Soloff et al. (2010) | PET | [18F]altanserin | Correlational | BIS-11, LHA, Buss-Durkee Hostility Inventory (BDHI) | na | 21 | 10 | 11 | 2 |
| 337 | Sommer et al. (2010) | fMRI | Inferring mental state from a cartoon, where protagonist's intention is either fulfilled or unfulfilled | Group comparison | PCL-R | Psychopathic criminals | 28 | | | 1 |
| 338 | Spoont et al. (2010) | PET | H2(15)O | Group comparison | na | Individuals with histories of repetitive violent behavior that began by early adolescence and has continued | 16 | 16 | 0 | 6 |
| 339a | Sterzer et al. (2005) | fMRI | Emotional pictures | Group comparison | CBCL | Adolescents with CD | 27 | 27 | 0 | 1 |
| 339b | Sterzer et al. (2005) | fMRI | Emotional pictures | Correlational | CBCL | Adolescents with CD | 27 | 27 | 0 | 1 |
| 340a | Sterzer et al. (2007) | MRI | VBM | Group comparison | DCL-SSV symptom checklist, CBCL, Impulsiveness-Venturesomeness-Empathy Questionnaire (IVE) | Adolescents with CD | 24 | 24 | 0 | 1 |
| 340b | Sterzer et al. (2007) | MRI | VBM | Correlational | DCL-SSV symptom checklist, CBCL, Impulsiveness-Venturesomeness-Empathy Questionnaire (IVE) | Adolescents with CD | 24 | 24 | 0 | 1 |
| 341 | Stevens & Haney-Caron (2012) | MRI | VBM | Group comparison | K-SADS-PL | CD | 48 | 32 | 16 | 1 |
| 342a | Storvestre et al. (2019) | MRI | Freesurfer | Group comparison | na | SCZ patients with histories of severe violence | 28 | 26 | 2 | 2 |
| 342b | Storvestre et al. (2019) | MRI | Freesurfer | Group comparison | na | SCZ patients with histories of severe violence | 30 | 27 | 3 | 2 |

| | | | | | | | | | | |
|---|---|---|---|---|---|---|---|---|---|---|
| 343 | Strenziok et al. (2011) | fMRI | Videos of real scenes of aggression (e.g. fist fights, street brawls and stadium violence) with differing levels of aggression (low/mild/moderate) | Activation study | na | na | 22 | 22 | 0 | 1 |
| 344a | Strenziok et al. (2011a) | fMRI | Imagining scenes of neutral, pleasant or aggressive social interaction | Activation study | the aggression questionnaire (AQ) | na | 20 | 20 | 0 | 1 |
| 344b | Strenziok et al. (2011a) | fMRI | Imagining scenes of neutral, pleasant or aggressive social interaction | Correlational | the aggression questionnaire (AQ) | na | 20 | 20 | 0 | 1 |
| 345 | Sun et al. (2018) | fMRI | the GoStop task | Correlational | Chinese version of BIS-11 | CD | 53 | 53 | 0 | 1 |
| 346 | Suridjan et al. (2012) | PET | [11C]-(+)-PHNO | Correlational | NEO-PI-R | na | 11 | 7 | 4 | 1 |
| 347 | Sutherland & Fishbein (2017) | PET | H2(15)O | Correlational | Levenson psychopathy scale | Abstinent substance abusers | 28 | 14 | 14 | 1 |
| 348 | Swartz et al. (2020) | fMRI | Emotional face matching task | Activation study | na | na | 49 | 25 | 24 | 1 |
| 349 | Szabó et al. (2017) | fMRI | Implicit facial expression recognition task | Correlational | ICU | na | 41 | 16 | 25 | 1 |
| 350 | Takahashi et al. (2018) | PET | 11C-Cetrozole | Correlational | Japanese version of the Buss-Perry Aggression Questionnaire (BPAQ) | na | 10 | 0 | 10 | 1 |
| 351 | Tang et al. (2013) | MRI | VBM | Group comparison | personality diagnostic questionnaire-4+ (PDQ-4+) | Youth criminal offenders with ASPD | 67 | 67 | 0 | 1 |
| 352 | Tessitore et al. (2016) | MRI | Freesurfer | Group comparison | MIDI | PD and ICD, but no PD-related dementia | 30 | 25 | 5 | 2 |
| 353 | Thiel et al. (2014) | fMRI | Emotional pictures (aggression, disgust, neutral) | Activation study | na | na | 15 | 10 | 5 | 1 |
| 354a | Thornton et al. (2017) | fMRI | Dot probe task with images of fearful faces, neutral faces or negative and inanimate objects (e.g. gun, knife) | Group comparison | KSADS, ICU | CD or ODD | 49 | 30 | 19 | 4 |
| 354b | Thornton et al. (2017) | fMRI | Dot probe task with images of fearful faces, neutral faces or negative and inanimate objects (e.g. gun, knife) | Correlational | KSADS, ICU | Youths with either CD or ODD | 29 | 20 | 9 | 2 |
| 355 | Tiihonen et al. (1997) | SPECT | 2β-carbomethoxy-3β(4-iodophenyl)tropane ([123I]β-CIT) | Group comparison | na | violent offenders, all but one alcoholics, most with ASPD | 31 | 29 | 2 | 1 |
| 356 | Tiihonen et al. (2008) | MRI | VBM | Group comparison | PCL-R | Violent offenders with ASPD | 51 | 51 | 0 | 1 |
| 357a | Tikàsz et al. (2016) | fMRI | Emotional pictures | Group comparison | na | SCZ or schizoaffective disorder and a history of armed aggression | 39 | 39 | 0 | 2 |
| 357b | Tikàsz et al. (2016) | fMRI | Emotional pictures | Group comparison | na | SCZ or schizoaffective disorder and a history of armed aggression | 41 | 41 | 0 | 2 |
| 357c | Tikàsz et al. (2016) | fMRI | Emotional pictures | Group comparison | na | SCZ or schizoaffective disorder and a history of armed aggression | 60 | 60 | 0 | 2 |
| 358a | Tikàsz et al. (2018) | fMRI | Affective Go/NoGo task | Group comparison | na | SCZ and a history of violence | 46 | 46 | 0 | 1 |
| 358b | Tikàsz et al. (2018) | fMRI | Affective Go/NoGo task | Group comparison | na | SCZ and a history of violence | 47 | 47 | 0 | 1 |
| 359a | Tonnaer et al. (2017) | fMRI | Anger Articulated Thoughts during Simulated Situations (ATSS) paradigm (focusing on or regulating emotional state while listening to happy, neutral or anger situation descriptions) | Group comparison | PCL-R, RPQ, Aggression questionnaire (AQ), Anger-Single Target Implicit Association Test (anger-STIAT) | Violent offenders | 34 | 34 | 0 | 8 |
| 359b | Tonnaer et al. (2017) | fMRI | Anger Articulated Thoughts during Simulated Situations (ATSS) paradigm | Correlational | PCL-R, RPQ, Aggression questionnaire (AQ), Anger-Single Target Implicit Association Test | Violent offenders | 34 | 34 | 0 | 2 |
| 360 | van de Giessen et al. (2014) | PET | [11C]DASB | Group comparison | BPAQ, OAS-Modified (OAS-M), LHA, Dimensional assessment of pesronality pathology (DAPP) callousness -scale, BIS-11, Affective Lability Scale (ALS) | IED and at least one DSM-IV personality disorder | 59 | 43 | 16 | 2 |
| 361 | Van den Bos et al. (2014) | fMRI | Mini-ultimatum game | Group comparison | YPI, Youth self report, CBCL | Juvenile delinquets with severely antisocial behavior | 34 | 34 | 0 | 2 |
| 362 | Van den Stock et al. (2015) | fMRI | Focusing to either aggressor or victim in realistic video clips of aggressive two-person interactions | Activation study | Dutch version of the Aggression Questionnaire, Dutch version of the I7-questionnaire | na | 15 | 15 | 0 | 1 |
| 363 | van Lith et al. (2018) | fMRI | Classical fear learning task | Group comparison | Dutch version of the YPI, CBCL, youth self report (YSR) | Adolescents with ODD and/or CD | 41 | 41 | 0 | 1 |
| 364 | Veit et al. (2010) | fMRI | TAGG | Correlational | PCL:SV, Levenson self report scale (LSRS) | Criminals with elevated psychopathy scores | 10 | 10 | 0 | 2 |
| 365 | Verger et al. (2018) | PET | 18F-Fluorodesoxyglucose (18F-FDG) | Group comparison | na | PD patients without cognitive decline and with ICDs | 36 | 25 | 11 | 1 |
| 366 | Veroude et al. (2016) | fMRI | MID | Correlational | K-SADS-PL, ICU | ADHD and either ODD or CD | 328 | 185 | 143 | 1 |
| 367a | Vetter et al. (2020) | MRI | Freesurfer | Group comparison | CBCL | ADHD and CD or ODD | 62 | 62 | 0 | 1 |
| 367b | Vetter et al. (2020) | MRI | Freesurfer | Group comparison | CBCL | ADHD and CD or ODD | 56 | 56 | 0 | 1 |
| 368a | Viding et al. (2012) | fMRI | Fearful and calm faces presented preattentively | Group comparison | Child and Adolescent Symptom Inventory (CASI-4R), ICU | boys with CD and high callous unemotional traits (median split, median ICU score 44,5) | 46 | 46 | 0 | 2 |
| 368b | Viding et al. (2012) | fMRI | Fearful and calm faces presented preattentively | Correlational | Child and Adolescent Symptom Inventory (CASI-4R), ICU | CD and high callous unemotional traits | 30 | 30 | 0 | 2 |
| 369 | Vieira et al. (2014) | MRI | VBM | Correlational | Triarchic Psychopathy Measure (TriPM), PPI-Revised (PPI-R) | na | 35 | 15 | 20 | 4 |
| 370a | Vieira et al. (2017) | fMRI | Approaching or withdrawing (by zooming) emotional faces | Activation study | PPI - revised (PPI-R) | na | 23 | 11 | 12 | 1 |
| 370b | Vieira et al. (2017) | fMRI | Approaching or withdrawing (by zooming) emotional faces | Correlational | PPI - revised (PPI-R) | na | 23 | 11 | 12 | 1 |
| 371 | Vincent et al. (2018) | fMRI | Drug-related or neutral pictures | Correlational | PCL:YV | Adolescent offenders with cocaine or methamphetamine abuse with at least one-week abstinence | 50 | 50 | 0 | 1 |
| 372 | Volkow et al. (1995) | PET | 18F-deoxyglucose (FDG) | Group comparison | na | Violent offenders with either IED or ASPD | 16 | | | 1 |
| 373 | Volman et al. (2016) | fMRI | Responding to emotional faces by pulling towards of pushing away with a joystick | Group comparison | PCL-R | Violent psychopathic offenders | 34 | 34 | 0 | 1 |
| 374a | von Polier et al. (2020) | fMRI | Inferring other's emotional state from emotional (morphed) faces or judging own emotional response to them (+ control task of judging the width of faces) | Group comparison | K-SADS-PL, German version of ASPD, german version of the Bryant index of Empathy (BIE) | Early onset-type CD | 29 | 29 | 0 | 3 |
| 374b | von Polier et al. (2020) | fMRI | Inferring other's emotional state from emotional (morphed) faces or judging own emotional response to them (+ control task of judging the width of faces) | Correlational | K-SADS-PL, German version of ASPD, german version of the Bryant index of Empathy (BIE) | Early onset-type CD | 29 | 29 | 0 | 1 |
| 375 | Voon et al. (2014) | SPECT | [123I]FP-CIT (2β-carbomethoxy-3β-(4-iodophenyl)tropane) | Group comparison | na | PD, ICD | 30 | 18 | 12 | 1 |
| 376a | Völlm et al. (2007) | fMRI | Responding as quickly as possible to specific visual stimuli to earn (reward task) or not to lose (loss task) money | Group comparison | BIS | Patients with either ASPD or BPD and a history of impulsive behaviour, including offending | 22 | 22 | 0 | 4 |
| 376b | Völlm et al. (2007) | fMRI | Responding as quickly as possible to specific visual stimuli to earn (reward task) or not to lose (loss task) money | Correlational | BIS | Patients with either ASPD or BPD and a history of impulsive behaviour, including offending | 8 | 8 | 0 | 2 |
| 377 | Völlm et al. (2004) | fMRI | Go-NoGo task | Group comparison | BIS, Impulsiveness-Venturesomeness-Empathy inventory (IVE) | ASPD or BPD | 16 | 16 | 0 | 2 |
| 378a | Wallace et al. (2014) | MRI | Freesurfer | Group comparison | ICU | CD | 49 | 33 | 16 | 2 |
| 378b | Wallace et al. (2014) | MRI | Freesurfer | Correlational | ICU | Youths with CD | 16 | 11 | 5 | 1 |

| | | | | | | | | | | |
|---|---|---|---|---|---|---|---|---|---|---|
| 379 | Weber et al. (2006) | fMRI | Playing a mature rated first-person-shooter game Tactical Ops: Assault on Terror | Correlational | na | na | 13 | 13 | 0 | 2 |
| 380 | White et al. (2014a) | fMRI | Doors task (reinforcement learning task with physical threat images, contamination threat images or appetitive images) | Group comparison | KSADS, ICU | Disruptive behavior disorders | 30 | 21 | 9 | 4 |
| 381 | White et al. (2014) | fMRI | Social Fairness Game (Ultimatum game with a possibility of monetary punishment by spending money) | Correlational | na | na | 21 | 12 | 9 | 2 |
| 382a | White et al. (2012) | fMRI | Emotion-attention bars task (judging whether two bars flanking an emotional face are parallel) | Group comparison | APSD | Psychopathic youths with CD or ODD | 32 | 21 | 11 | 4 |
| 382b | White et al. (2012) | fMRI | Emotion-attention bars task (judging whether two bars flanking an emotional face are parallel) | Correlational | APSD | Psychopathic youths with CD or ODD | 32 | 21 | 11 | 1 |
| 383 | White et al. (2013) | fMRI | PAT with monetary rewards and punishments | Group comparison | APSD | CD or ODD | 38 | 27 | 11 | 4 |
| 384 | White et al. (2016) | fMRI | PAT with monetary rewards and punishments | Correlational | CBCL, ICU | na | 72 | 52 | 20 | 3 |
| 385a | White et al. (2016a) | fMRI | Social fairness game (ultimatum game with monetary punishments) | Group comparison | ICU, RPQ | CD or ODD and low callous-unemotional traits | 41 | | | 3 |
| 385b | White et al. (2016a) | fMRI | Social fairness game (ultimatum game with monetary punishments) | Group comparison | ICU, RPQ | CD or ODD and low callous-unemotional traits | 30 | 19 | 11 | 2 |
| 386 | White et al. (2018) | fMRI | Approaching or receding (zooming in/out) animate/inanimate, threatening/neutral pictures | Group comparison | ICU | Youth with disruptive behavior disorders | 58 | 36 | 22 | 14 |
| 387 | White et al. (2012a) | fMRI | Determining on which side of screen a probe "x" is when emotional faces are looking towards or away from it | Group comparison | APSD | Youths with either CD or ODD and psychopathic traits | 36 | 22 | 14 | 3 |
| 388a | Wiggins et al. (2016) | fMRI | Emotion labeling task | Correlational | The affective reactivity index | Disruptive mood dysregulation disorder | 25 | 15 | 10 | 3 |
| 388b | Wiggins et al. (2016) | fMRI | Emotion labeling task | Correlational | The affective reactivity index | Bipolar disorder | 24 | 16 | 8 | 1 |
| 389 | Witte et al. (2009) | PET | [carbonyl-11C]WAY-100635 | Group comparison | Questionnaire for Measuring Factors of Aggression (German, Hampel and Selg, 1975) | Healthy participants with high aggression scores | 33 | 17 | 16 | 1 |
| 390a | Woermann et al. (2000) | MRI | VBM | Group comparison | social dysfunction and aggression scale (SDAS-21) | Temporal lobe epilepsy and a history of episodic affective aggression | 48 | | | 1 |
| 390b | Woermann et al. (2000) | MRI | VBM | Correlational | social dysfunction and aggression scale (SDAS-21) | Temporal lobe epilepsy and a history of episodic affective aggression | 48 | | | 1 |
| 391 | Yang et al. (2010) | MRI | Volumetric segmentation, cortical pattern matching, surface-based mesh modeling | Group comparison | PCL-R | Psychopathic offenders | 43 | 36 | 7 | 3 |
| 392a | Yang et al. (2010a) | MRI | Hybrid discriminative/generative models | Group comparison | na | Detainees accused of homicide undergoing forensic psychiatric evaluation | 50 | 6 | 44 | 1 |
| 392b | Yang et al. (2010a) | MRI | Hybrid discriminative/generative models | Group comparison | na | Schizophrenic murderers | 41 | 6 | 35 | 1 |
| 393a | Yang et al. (2009) | MRI | Manual tracing | Group comparison | PCL-R | Community individuals with high psychopathy | 59 | 51 | 8 | 1 |
| 393b | Yang et al. (2009) | MRI | Manual tracing | Group comparison | PCL-R | Community individuals with high psychopathy | 59 | 51 | 8 | 2 |
| 394a | Yang et al. (2015) | MRI | Freesurfer | Correlational | a slightly extended version of the Childhood Psychopathy Scale (CPS) | na | 108 | 54 | 54 | 2 |
| 394b | Yang et al. (2015) | MRI | Freesurfer | Correlational | a slightly extended version of the Childhood Psychopathy Scale (CPS) | na | 108 | 54 | 54 | 1 |
| 395 | Yang et al. (2005) | MRI | Semiautomatic segmentation with CAMRA S200 ALLEGRO software | Correlational | PCL-R | Psychopathic individuals | 52 | 52 | 0 | 1 |
| 396 | Yang et al. (2007) | SPECT | [123I]-labeled 2-((2-((dimethylamino)methyl) phenyl)thio)-5-iodophenylamine ([123I]ADAM) | Correlational | Cook-Medley Hostility Scale (Ho) | na | 10 | 4 | 6 | 1 |
| 397 | Yoder et al. (2021) | fMRI | Matching emotional faces to probable reactions of perpetrator/agent or victim/beneficiary in morally laden social interactions | Correlational | PCL-R | Inmates | 107 | 0 | 107 | 3 |
| 398 | Yoder et al. (2015) | fMRI | Implicit and explicit moral judging of visual scenes of interpersonal harm or assistance | Correlational | PCL-R | Incarcerated criminals | 94 | 94 | 0 | 4 |
| 399 | Yoder et al. (2015a) | fMRI | Videos of mixed martial arts or capoeira as control | Correlational | PPI-R | na | 43 | 43 | 0 | 4 |
| 400 | Zhang et al. (2013) | MRI | Model-based integrated segmentation/registration tool - FIRST within the FSL | Group comparison | the Conflict Tactics Scale (CTS) | Alcoholics with history of domestic violence and various cluster B and C personality disorders | 38 | 38 | 0 | 1 |
| 401 | Zhang et al. (2019) | MRI | VBM | Group comparison | na | Adolescent violent offenders | 59 | 59 | 0 | 1 |
| 402a | Zhang et al. (2019a) | MRI | VBM | Correlational | BIS | CD | 60 | 60 | 0 | 1 |
| 402b | Zhang et al. (2019a) | MRI | VBM | Correlational | BIS | na | 60 | 60 | 0 | 2 |
| 403 | Zhang et al. (2018) | MRI | VBM | Group comparison | BIS | Adolescents with CD | 120 | 120 | 0 | 2 |
| 404 | Zhu et al. (2019) | MRI | VBM | Correlational | RPQ | na | 240 | 112 | 128 | 2 |
| 405 | Zhu et al. (2014) | fMRI | GoStop task | Group comparison | na | Children with pure ODD | 21 | 21 | 0 | 2 |
| 406 | Zijlmans et al. (2018) | fMRI | Judging the severity of moral violations immoral negative, non-moral negative and neutral pictures | Correlational | Youth Psychopathy Inventory - Short Version (YPI-SV) | Multi-problem young adults | 100 | 100 | 0 | 1 |

Abbreviations: AMII = Angry mood improvement inventory, APSD = Antisocial process screening device, ASPD = Antisocial personality disorder, ASRS = Adult ADHD Self-Report Scale, AST = Affective stroop task, BAQ = Brief Aggression Questionnaire, BDHI = Buss-Durkee Hostility Index, BIS = Barratt Impulsiveness Scale, BPAQ= Buss-Perry Aggression Questionnaire, BPD = Borderline personality disorder, CBCL=Child Behavior Checklist, CBG = Colorado Balloon Game, CD = Conduct disorder, CIDI(-SAM) = Composite International Diagnostic Interview(-Substance Abuse Module), CP = Cyberball paradigm, DISC-IV = the Diagnostic Interview Schedule for Children, DY-BOCS = Dimensional Yale-Brown Obsessive-Compulsive Scale, EIS = Eysenck Junior Impulsiveness Scale, ESI = Externalizing Spectrum Inventory, FAF = Aggressivity factors questionnaire, ICD = Impulse control disorder, ICU = Inventory of Callous-Unemotional Traits, IED = Intermittent explosive disorder, IRI = Interpersonal reactivity index, KSADS-PL = Kiddie Schedule for Affective Disorders and Schizophrenia Present and Lifetime Version, LHA = Life history of aggression, MDD = Major depressive disorder ,MID = Monetary incentive delay task ,MIDI = Minnesota impulsive disorders interview, MMPI2 = Minnesota Multiphasic Personality Inventory-2, MPQ = Multidimensional personality questionnaire, NEO-PI-R = The Revised NEO Personality Inventory, OAS = Overt Aggression Scale, OCD = Obsessive-compulsive disorder, ODD = Oppositional defiant disorder, PANSS = Positive and Negative Syndrome Scale, PAT = Passive avoidance Task, PBS = Pediatric Behavior Scale, short scale, PCL-R = Psychopathy Checklist – Revised, PCL-SV = Psychopathy checklist: screening version, PCL-YV = Psychopathy checklist: youth version, PD = Parkinson's disease, PPI = Psychopathic Personality Inventory, PSAP = Point Subtraction Aggression Paradigm, PTSD = Posttraumatic stress disorder, QUIP-RS = Questionnaire for Impulsive-Compulsive Disorder in Parkinson's Disease-Rating Scale, RPQ = Reactive and Proactive Aggression Questionnaire, SCL-90-R = Symptom Checklist-90-Revised, SCZ = Schizophrenia, SDQ = Strengths and difficulties questionnaire, SNAT = Social Network Aggression Task, SSAGA-II = Semi-Structured Assessment For The Genetics Of Alcoholism-II, SST= Stop-signal task, STAXI = State-Trait Anger Expression Inventory, STAXI-2 = revised State-Trait Anger Expression Inventory, TAGG = Taylor's aggression paradigm, WHOQOL-BREF = The World Health Organization Quality of Life-BREF, YPI = Youth psychopathic traits inventory

#### Supplementary references (included studies)


Abe, N., Greene, J. D., & Kiehl, K. A. (2018). Reduced engagement of the anterior cingulate cortex in the dishonest decision-making of incarcerated psychopaths. *Social Cognitive and Affective Neuroscience*, *13*(8), 797–807. https://doi.org/10.1093/scan/nsy050

Achterberg, M., van Duijvenvoorde, A. C. K., Bakermans-Kranenburg, M. J., & Crone, E. A. (2016). Control your anger! The neural basis of aggression regulation in response to negative social feedback. *Social Cognitive and Affective Neuroscience*, *11*(5), 712–720. https://doi.org/10.1093/scan/nsv154

Achterberg, M., van Duijvenvoorde, A. C. K., van der Meulen, M., Bakermans-Kranenburg, M. J., & Crone, E. A. (2018). Heritability of aggression following social evaluation in middle childhood: An fMRI study. *Human Brain Mapping*, *39*(7), 2828–2841. https://doi.org/10.1002/hbm.24043

Achterberg, M., van Duijvenvoorde, A. C. K., van IJzendoorn, M. H., Bakermans-Kranenburg, M. J., & Crone, E. A. (2020). Longitudinal changes in DLPFC activation during childhood are related to decreased aggression following social rejection. *Proceedings of the National Academy of Sciences of the United States of America*, *117*(15), 8602–8610. https://doi.org/10.1073/pnas.1915124117

Aggensteiner, P.-M., Holz, N. E., Böttinger, B. W., Baumeister, S., Hohmann, S., Werhahn, J. E., Naaijen, J., Ilbegi, S., Glennon, J. C., Hoekstra, P. J., Banaschewski, T., & Brandeis, D. (2020). The effects of callous-unemotional traits and aggression subtypes on amygdala activity in response to negative faces. *Psychological Medicine*. https://doi.org/10.1017/S0033291720002111

Aghajani, M., Klapwijk, E. T., Andershed, H., Fanti, K. A., van der Wee, N. J. A., Vermeiren, R. R. J. M., & Colins, O. F. (2021). Neural processing of socioemotional content in conduct-disordered juvenile offenders with limited prosocial emotions. *Progress in Neuro-Psychopharmacology and Biological Psychiatry*, *105*. https://doi.org/10.1016/j.pnpbp.2020.110045

Aharoni, E., Vincent, G. M., Harenski, C. L., Calhoun, V. D., Sinnott-Armstrong, W., Gazzaniga, M. S., & Kiehl, K. A. (2013). Neuroprediction of future rearrest. *Proceedings of the National Academy of Sciences of the United States of America*, *110*(15), 6223–6228. https://doi.org/10.1073/pnas.1219302110

Alia-Klein, N., Goldstein, R. Z., Kriplani, A., Logan, J., Tomasi, D., Williams, B., Telang, F., Shumay, E., Biegon, A., Craig, I. W., Volkow, N. D., & Fowler, J. S. (2008). Brain monoamine oxidase A activity predicts trait aggression. *Journal of Neuroscience*, *28*(19), 5099–5104. https://doi.org/10.1523/JNEUROSCI.0925-08.2008

Alia-Klein, N., Goldstein, R. Z., Tomasi, D., Woicik, P. A., Moeller, S. J., Williams, B., Craig, I. W., Telang, F., Biegon, A., Wang, G.-J., Fowler, J. S., & Volkow, N. D. (2009). Neural Mechanisms of Anger Regulation as a Function of Genetic Risk for Violence. *Emotion*, *9*(3), 385–396. https://doi.org/10.1037/a0015904

Alia-Klein, N., Wang, G.-J., Preston-Campbell, R. N., Moeller, S. J., Parvaz, M. A., Zhu, W., Jayne, M. C., Wong, C., Tomasi, D., Goldstein, R. Z., Fowler, J. S., & Volkow, N. D. (2014). Reactions to media violence: It's in the brain of the beholder. *PLoS ONE*, *9*(9). https://doi.org/10.1371/journal.pone.0107260

Alvarenga, P. G., do Rosário, M. C., Batistuzzo, M. C., Diniz, J. B., Shavitt, R. G., Duran, F. L. S., Dougherty, D. D., Bressan, R. A., Miguel, E. C., & Hoexter, M. Q. (2012). Obsessive-compulsive symptom dimensions correlate to specific gray matter volumes in treatment-naïve patients. *Journal of Psychiatric Research*, *46*(12), 1635–1642. https://doi.org/10.1016/j.jpsychires.2012.09.002

Amen, D. G., & Carmichael, B. (1997). Oppositional children similar to ocd on spect: Implications for treatment. *International Journal of Phytoremediation*, *21*(1), 1–6. https://doi.org/10.1300/J184v02n02_01

Amen, D. G., Hanks, C., Prunella, J. R., & Green, A. (2007). An analysis of regional cerebral blood flow in impulsive murderers using single photon emission computed tomography. *Journal of Neuropsychiatry and Clinical Neurosciences*, *19*(3), 304–309. https://doi.org/10.1176/jnp.2007.19.3.304

Amen, D. G., Stubblefield, M., Carmichael, B., & Thisted, R. (1996). Brain SPECT findings and aggressiveness. *Annals of Clinical Psychiatry*, *8*(3), 129–137. https://doi.org/10.3109/10401239609147750

Anderson, N. E., Maurer, J. M., Steele, V. R., & Kiehl, K. A. (2018). Psychopathic traits associated with abnormal hemodynamic activity in salience and default mode networks during auditory oddball task. *Cognitive, Affective and Behavioral Neuroscience*, *18*(3), 564–580. https://doi.org/10.3758/s13415-018-0588-2

Anderson, N. E., Steele, V. R., Maurer, J. M., Rao, V., Koenigs, M. R., Decety, J., Kosson, D. S., Calhoun, V. D., & Kiehl, K. A. (2017). Differentiating emotional processing and attention in psychopathy with functional neuroimaging. *Cognitive, Affective and Behavioral Neuroscience*, *17*(3), 491–515. https://doi.org/10.3758/s13415-016-0493-5

Baird, A. A., Silver, S. H., & Veague, H. B. (2010). Cognitive control reduces sensitivity to relational aggression among adolescent girls. *Social Neuroscience*, *5*(5), 519–532. https://doi.org/10.1080/17470911003747386

Barkataki, I., Kumari, V., Das, M., Sumich, A., Taylor, P., & Sharma, T. (2008). Neural correlates of deficient response inhibition in mentally disordered violent individuals. *Behavioral Sciences and the Law*, *26*(1), 51–64. https://doi.org/10.1002/bsl.787

Barkataki, I., Kumari, V., Das, M., Taylor, P., & Sharma, T. (2006). Volumetric structural brain abnormalities in men with schizophrenia or antisocial personality disorder. *Behavioural Brain Research*, *169*(2), 239–247. https://doi.org/10.1016/j.bbr.2006.01.009

Baskin-Sommers, A. R., Curtin, J. J., Larson, C. L., Stout, D., Kiehl, K. A., & Newman, J. P. (2012). Characterizing the anomalous cognition-emotion interactions in externalizing. *Biological Psychology*, *91*(1), 48–58. https://doi.org/10.1016/j.biopsycho.2012.05.001

Beames, J. R., Gilam, G., Schofield, T. P., Schira, M. M., & Denson, T. F. (2020). The impact of self-control training on neural responses following anger provocation. *Social Neuroscience*, *15*(5), 558–570. https://doi.org/10.1080/17470919.2020.1799860

Beaver, J. D., Lawrence, A. D., Passamonti, L., & Calder, A. J. (2008). Appetitive motivation predicts the neural response to facial signals of aggression. *Journal of Neuroscience*, *28*(11), 2719–2725. https://doi.org/10.1523/JNEUROSCI.0033-08.2008

Beckwith, T. J., Dietrich, K. N., Wright, J. P., Altaye, M., & Cecil, K. M. (2021). Criminal arrests associated with reduced regional brain volumes in an adult population with documented childhood lead exposure. *Environmental Research*, *201*. https://doi.org/10.1016/j.envres.2021.111559

Beckwith, T. J., Dietrich, K. N., Wright, J. P., Altaye, M., & Cecil, K. M. (2018). Reduced regional volumes associated with total psychopathy scores in an adult population with childhood lead exposure. *NeuroToxicology*, *67*, 1–26. https://doi.org/10.1016/j.neuro.2018.04.004

Benegal, V., Antony, G., Venkatasubramanian, G., & Jayakumar, P. N. (2007). Gray matter volume abnormalities and externalizing symptoms in subjects at high risk for alcohol dependence. *Addiction Biology*, *12*(1), 122–132. https://doi.org/10.1111/j.1369-1600.2006.00043.x

Bertsch, K., Grothe, M., Prehn, K., Vohs, K., Berger, C., Hauenstein, K., Keiper, P., Domes, G., Teipel, S., & Herpertz, S. C. (2013). Brain volumes differ between diagnostic groups of violent criminal offenders. *European Archives of Psychiatry and Clinical Neuroscience*, *263*(7), 593–606. https://doi.org/10.1007/s00406-013-0391-6

Besteher, B., Squarcina, L., Spalthoff, R., Bellani, M., Gaser, C., Brambilla, P., & Nenadić, I. (2017). Brain structural correlates of irritability: Findings in a large healthy cohort. *Human Brain Mapping*, *38*(12), 6230–6238. https://doi.org/10.1002/hbm.23824

Beyer, F., Münte, T. F., Göttlich, M., & Krämer, U. M. (2015). Orbitofrontal cortex reactivity to angry facial expression in a social interaction correlates with aggressive behavior. *Cerebral Cortex*, *25*(9), 3057–3063. https://doi.org/10.1093/cercor/bhu101

Beyer, F., Münte, T. F., & Krämer, U. M. (2014). Increased neural reactivity to socio-emotional stimuli links social exclusion and aggression. *Biological Psychology*, *96*(1), 102–110. https://doi.org/10.1016/j.biopsycho.2013.12.008

Birbaumer, N., Veit, R., Lotze, M., Erb, M., Hermann, C., Grodd, W., & Flor, H. (2005). Deficient fear conditioning in psychopathy: A functional magnetic resonance imaging study. *Archives of General Psychiatry*, *62*(7), 799–805. https://doi.org/10.1001/archpsyc.62.7.799

Biundo, R., Weis, L., Facchini, S., Formento-Dojot, P., Vallelunga, A., Pilleri, M., Weintraub, D., & Antonini, A. (2015). Patterns of cortical thickness associated with impulse control disorders in Parkinson's disease. *Movement Disorders*, *30*(5), 688–695. https://doi.org/10.1002/mds.26154

Bjork, J. M., Chen, G., & Hommer, D. W. (2012). Psychopathic tendencies and mesolimbic recruitment by cues for instrumental and passively obtained rewards. *Biological Psychology*, *89*(2), 408–415. https://doi.org/10.1016/j.biopsycho.2011.12.003

Bjork, J. M., Chen, G., Smith, A. R., & Hommer, D. W. (2010). Incentive-elicited mesolimbic activation and externalizing symptomatology in adolescents. *Journal of Child Psychology and Psychiatry and Allied Disciplines*, *51*(7), 827–837. https://doi.org/10.1111/j.1469-7610.2009.02201.x

Bobes, M. A., Ostrosky, F., Diaz, K., Romero, C., Borja, K., Santos, Y., & Valdés-Sosa, M. (2013). Linkage of functional and structural anomalies in the left amygdala of reactive-aggressive men. *Social Cognitive and Affective Neuroscience*, *8*(8), 928–936. https://doi.org/10.1093/scan/nss101

Boccardi, M., Bocchetta, M., Aronen, H. J., Repo-Tiihonen, E., Vaurio, O., Thompson, P. M., Tiihonen, J., & Frisoni, G. B. (2013). Atypical nucleus accumbens morphology in psychopathy: Another limbic piece in the puzzle. *International Journal of Law and Psychiatry*, *36*(2), 157–167. https://doi.org/10.1016/j.ijlp.2013.01.008

Boccardi, M., Frisoni, G. B., Hare, R. D., Cavedo, E., Najt, P., Pievani, M., Rasser, P. E., Laakso, M. P., Aronen, H. J., Repo-Tiihonen, E., Thompson, P. M., & Tiihonen, J. (2011). Cortex and amygdala morphology in psychopathy. *Psychiatry Research - Neuroimaging*, *193*(2), 85–92. https://doi.org/10.1016/j.pscychresns.2010.12.013

Boccardi, M., Ganzola, R., Rossi, R., Sabattoli, F., Laakso, M. P., Repo-Tiihonen, E., Vaurio, O., Könönen, M., Aronen, H. J., Thompson, P. M., Frisoni, G. B., & Tiihonen, J. (2010). Abnormal hippocampal shape in offenders with psychopathy. *Human Brain Mapping*, *31*(3), 438–447. https://doi.org/10.1002/hbm.20877

Boes, A. D., Bechara, A., Tranel, D., Anderson, S. W., Richman, L., & Nopoulos, P. (2009). Right ventromedial prefrontal cortex: A neuroanatomical correlate of impulse control in boys. *Social Cognitive and Affective Neuroscience*, *4*(1), 1–9. https://doi.org/10.1093/scan/nsn035

Boes, A. D., Tranel, D., Anderson, S. W., & Nopoulos, P. (2008). Right Anterior Cingulate: A Neuroanatomical Correlate of Aggression and Defiance in Boys. *Behavioral Neuroscience*, *122*(3), 677–684. https://doi.org/10.1037/0735-7044.122.3.677

Breitschuh, S., Schöne, M., Tozzi, L., Kaufmann, J., Strumpf, H., Fenker, D., Frodl, T., Bogerts, B., & Schiltz, K. (2018). Aggressiveness of martial artists correlates with reduced temporal pole grey matter concentration. *Psychiatry Research - Neuroimaging*, *281*, 24–30. https://doi.org/10.1016/j.pscychresns.2018.08.001

Brunnlieb, C., Münte, T. F., Krämer, U., Tempelmann, C., & Heldmann, M. (2013). Vasopressin modulates neural responses during human reactive aggression. *Social Neuroscience*, *8*(2), 148–164. https://doi.org/10.1080/17470919.2013.763654

Buades-Rotger, M., Beyer, F., & Krämer, U. M. (2017). Avoidant responses to interpersonal provocation are associated with increased amygdala and decreased mentalizing network activity. *eNeuro*, *4*(3). https://doi.org/10.1523/ENEURO.0337-16.2017

Buckholtz, J. W., Treadway, M. T., Cowan, R. L., Woodward, N. D., Benning, S. D., Li, R., Ansari, M. S., Baldwin, R. M., Schwartzman, A. N., Shelby, E. S., Kessler, R. M., & Zald, D. H. (2010). Mesolimbic dopamine reward system hypersensitivity in individuals with psychopathic traits. *Nature Neuroscience*, *13*(4), 419–421. https://doi.org/10.1038/nn.2510

Budhiraja, M., Pereira, J. B., Lindner, P., Westman, E., Jokinen, J., Savic, I., Tiihonen, J., & Hodgins, S. (2019). Cortical structure abnormalities in females with conduct disorder prior to age 15. *Psychiatry Research - Neuroimaging*, *289*, 37–44. https://doi.org/10.1016/j.pscychresns.2018.12.004

Budhiraja, M., Savic, I., Lindner, P., Jokinen, J., Tiihonen, J., & Hodgins, S. (2017). Brain structure abnormalities in young women who presented conduct disorder in childhood/adolescence. *Cognitive, Affective and Behavioral Neuroscience*, *17*(4), 869–885. https://doi.org/10.3758/s13415-017-0519-7

Byrd, A. L., Hawes, S. W., Burke, J. D., Loeber, R., & Pardini, D. A. (2018). Boys with conduct problems and callous-unemotional traits: Neural response to reward and punishment and associations with treatment response. *Developmental Cognitive Neuroscience*, *30*, 51–59. https://doi.org/10.1016/j.dcn.2017.12.004

Caldwell, B. M., Anderson, N. E., Harenski, K. A., Sitney, M. H., Caldwell, M. F., Van Rybroek, G. J., & Kiehl, K. A. (2019). The structural brain correlates of callous-unemotional traits in incarcerated male adolescents. *NeuroImage: Clinical*, *22*. https://doi.org/10.1016/j.nicl.2019.101703

Caldwell, B. M., Harenski, C. L., Harenski, K. A., Fede, S. J., Steele, V. R., Koenigs, M. R., & Kiehl, K. A. (2015). Abnormal frontostriatal activity in recently abstinent cocaine users during implicit moral processing. *Frontiers in Human Neuroscience*, *9*(OCTOBER). https://doi.org/10.3389/fnhum.2015.00565

Cardinale, E. M., Breeden, A. L., Robertson, E. L., Lozier, L. M., Vanmeter, J. W., & Marsh, A. A. (2018). Externalizing behavior severity in youths with callous-unemotional traits corresponds to patterns of amygdala activity and connectivity during judgments of causing fear. *Development and Psychopathology*, *30*(1), 191–201. https://doi.org/10.1017/S0954579417000566

Carlson, J. M., Greenberg, T., & Mujica-Parodi, L. R. (2010). Blind rage? Heightened anger is associated with altered amygdala responses to masked and unmasked fearful faces. *Psychiatry Research - Neuroimaging*, *182*(3), 281–283. https://doi.org/10.1016/j.pscychresns.2010.02.001

Carré, J. M., Murphy, K. R., & Hariri, A. R. (2013). What lies beneath the face of aggression? *Social Cognitive and Affective Neuroscience*, *8*(2), 224–229. https://doi.org/10.1093/scan/nsr096

Castellanos-Ryan, N., Struve, M., Whelan, R., Banaschewski, T., Barker, G. J., Bokde, A. L. W., Bromberg, U., Büchel, C., Flor, H., Fauth-Bühler, M., Garavan, H., & Conrod, P. J. (2014). Neural and cognitive correlates of the common and specific variance across externalizing problems in young adolescence. *American Journal of Psychiatry*, *171*(12), 1310–1319. https://doi.org/10.1176/appi.ajp.2014.13111499

Chang, C., Gau, S. S.-F., Huang, W.-S., Shiue, C.-Y., & Yeh, C.-B. (2017). Abnormal serotonin transporter availability in the brains of adults with conduct disorder. *Journal of the Formosan Medical Association*, *116*(6), 469–475. https://doi.org/10.1016/j.jfma.2016.07.012

Charpentier, J., Dzemidzic, M., West, J., Oberlin, B. G., Eiler, W. J. A., Saykin, A. J., & Kareken, D. A. (2016). Externalizing personality traits, empathy, and gray matter volume in healthy young drinkers. *Psychiatry Research - Neuroimaging*, *248*, 64–72. https://doi.org/10.1016/j.pscychresns.2016.01.006

Chen, B., Wu, X., Geniole, S. N., Ge, Q., Chen, Q., & Zhao, Y. (2021). Neural activity during provocation and aggressive responses in people from different social classes. *Current Psychology*. https://doi.org/10.1007/s12144-021-01925-y

Chester, D. S., & DeWall, C. N. (2016). The pleasure of revenge: Retaliatory aggression arises from a neural imbalance toward reward. *Social Cognitive and Affective Neuroscience*, *11*(7), 1173–1182. https://doi.org/10.1093/scan/nsv082

Chester, D. S., Lynam, D. R., Milich, R., & DeWall, C. N. (2018). Neural mechanisms of the rejection-aggression link. *Social Cognitive and Affective Neuroscience*, *13*(5), 501–512. https://doi.org/10.1093/scan/nsy025

Chester, D. S., Lynam, D. R., Milich, R., & DeWall, C. N. (2017). Physical aggressiveness and gray matter deficits in ventromedial prefrontal cortex. *Cortex*, *97*, 17–22. https://doi.org/10.1016/j.cortex.2017.09.024

Choe, D. E., Shaw, D. S., & Forbes, E. E. (2015). Maladaptive social information processing in childhood predicts young men's atypical amygdala reactivity to threat. *Journal of Child Psychology and Psychiatry and Allied Disciplines*, *56*(5), 549–557. https://doi.org/10.1111/jcpp.12316

Chumachenko, S. Y., Sakai, J. T., Dalwani, M. S., Mikulich-Gilbertson, S. K., Dunn, R., Tanabe, J., Young, S., McWilliams, S. K., Banich, M. T., & Crowley, T. J. (2015). Brain cortical thickness in male adolescents with serious substance use and conduct problems. *American Journal of Drug and Alcohol Abuse*, *41*(5), 414–424. https://doi.org/10.3109/00952990.2015.1058389

Coccaro, E. F., Cremers, H., Fanning, J., Nosal, E., Lee, R., Keedy, S., & Jacobson, K. C. (2018). Reduced frontal grey matter, life history of aggression, and underlying genetic influence. *Psychiatry Research - Neuroimaging*, *271*, 126–134. https://doi.org/10.1016/j.pscychresns.2017.11.007

Coccaro, E. F., Fitzgerald, D. A., Lee, R., McCloskey, M., & Phan, K. L. (2016). Frontolimbic Morphometric Abnormalities in Intermittent Explosive Disorder and Aggression. *Biological Psychiatry: Cognitive Neuroscience and Neuroimaging*, *1*(1), 32–38. https://doi.org/10.1016/j.bpsc.2015.09.006

Coccaro, E. F., Keedy, S. K., Gorka, S. M., King, A. C., Fanning, J. R., Lee, R. J., & Phan, K. L. (2016). Differential fMRI BOLD responses in amygdala in intermittent explosive disorder as a function of past Alcohol Use Disorder. *Psychiatry Research - Neuroimaging*, *257*, 5–10. https://doi.org/10.1016/j.pscychresns.2016.09.001

Coccaro, E. F., McCloskey, M. S., Fitzgerald, D. A., & Phan, K. L. (2007). Amygdala and Orbitofrontal Reactivity to Social Threat in Individuals with Impulsive Aggression. *Biological Psychiatry*, *62*(2), 168–178. https://doi.org/10.1016/j.biopsych.2006.08.024

Cohn, M. D., Popma, A., Van Den Brink, W., Pape, L. E., Kindt, M., Van Domburgh, L., Doreleijers, T. A. H., & Veltman, D. J. (2013). Fear conditioning, persistence of disruptive behavior and psychopathic traits: An fMRI study. *Translational Psychiatry*, *3*. https://doi.org/10.1038/tp.2013.89

Cohn, M. D., van Lith, K., Kindt, M., Pape, L. E., Doreleijers, A. H., van den Brink, W., Veltman, D. J., & Popma, A. (2016). Fear extinction, persistent disruptive behavior and psychopathic traits: FMRI in late adolescence. *Social Cognitive and Affective Neuroscience*, *11*(7), 1027–1035. https://doi.org/10.1093/scan/nsv067

Cohn, M. D., Veltman, D. J., Pape, L. E., Van Lith, K., Vermeiren, R. R. J. M., Van Den Brink, W., Doreleijers, T. A. H., & Popma, A. (2015). Incentive processing in persistent disruptive behavior and psychopathic traits: A functional magnetic resonance imaging study in adolescents. *Biological Psychiatry*, *78*(9), 615–624. https://doi.org/10.1016/j.biopsych.2014.08.017

Cohn, M. D., Viding, E., McCrory, E., Pape, L., van den Brink, W., Doreleijers, T. A. H., Veltman, D. J., & Popma, A. (2016). Regional grey matter volume and concentration in at-risk adolescents: Untangling associations with callous-unemotional traits and conduct disorder symptoms. *Psychiatry Research - Neuroimaging*, *254*, 180–187. https://doi.org/10.1016/j.pscychresns.2016.07.003

Contreras-rodríguez, O., Pujol, J., Batalla, I., Harrison, B. J., Bosque, J., Ibern-regàs, I., Hernández-ribas, R., Soriano-mas, C., Deus, J., López-solà, M., Menchón, J. M., & Cardoner, N. (2014). Disrupted neural processing of emotional faces in psychopathy. *Social Cognitive and Affective Neuroscience*, *9*(4), 505–512. https://doi.org/10.1093/scan/nst014

Contreras-Rodríguez, O., Pujol, J., Batalla, I., Harrison, B. J., Soriano-Mas, C., Deus, J., López-Solà, M., Macià, D., Pera, V., Hernández-Ribas, R., Menchón, J. M., & Cardoner, N. (2015). Functional connectivity bias in the prefrontal cortex of psychopaths. *Biological Psychiatry*, *78*(9), 647–655. https://doi.org/10.1016/j.biopsych.2014.03.007

Cope, L. M., Ermer, E., Gaudet, L. M., Steele, V. R., Eckhardt, A. L., Arbabshirani, M. R., Caldwell, M. F., Calhoun, V. D., & Kiehl, K. A. (2014). Abnormal brain structure in youth who commit homicide. *NeuroImage: Clinical*, *4*, 800–807. https://doi.org/10.1016/j.nicl.2014.05.002

Cope, L. M., Ermer, E., Nyalakanti, P. K., Calhoun, V. D., & Kiehl, K. A. (2014). Paralimbic gray matter reductions in incarcerated adolescent females with psychopathic traits. *Journal of Abnormal Child Psychology*, *42*(4), 659–668. https://doi.org/10.1007/s10802-013-9810-4

Cope, L. M., Shane, M. S., Segall, J. M., Nyalakanti, P. K., Stevens, M. C., Pearlson, G. D., Calhoun, V. D., & Kiehl, K. A. (2012). Examining the effect of psychopathic traits on gray matter volume in a community substance abuse sample. *Psychiatry Research - Neuroimaging*, *204*(2–3), 91–100. https://doi.org/10.1016/j.pscychresns.2012.10.004

Cope, L. M., Vincent, G. M., Jobelius, J. L., Nyalakanti, P. K., Calhoun, V. D., & Kiehl, K. A. (2014). Psychopathic traits modulate brain responses to drug cues in incarcerated offenders. *Frontiers in Human Neuroscience*, *8*(1 FEB). https://doi.org/10.3389/fnhum.2014.00087

Cropley, V. L., Tian, Y., Fernando, K., Mansour L., S., Pantelis, C., Cocchi, L., & Zalesky, A. (2021). Brain-Predicted Age Associates With Psychopathology Dimensions in Youths. *Biological Psychiatry: Cognitive Neuroscience and Neuroimaging*, *6*(4), 410–419. https://doi.org/10.1016/j.bpsc.2020.07.014

Crowley, T. J., Dalwani, M. S., Mikulich-Gilbertson, S. K., Du, Y. P., Lejuez, C. W., Raymond, K. M., & Banich, M. T. (2010). Risky decisions and their consequences: Neural processing by boys with antisocial substance disorder. *PLoS ONE*, *5*(9), 1–20. https://doi.org/10.1371/journal.pone.0012835

Crowley, T. J., Dalwani, M. S., Mikulich-Gilbertson, S. K., Young, S. E., Sakai, J. T., Raymond, K. M., McWilliams, S. K., Roark, M. J., & Banich, M. T. (2015). Adolescents neural processing of risky decisions: Effects of sex and behavioral disinhibition. *PLoS ONE*, *10*(7). https://doi.org/10.1371/journal.pone.0132322

da Cunha-Bang, S., Fisher, P. M., Hjordt, L. V., Holst, K., & Knudsen, G. M. (2019). Amygdala reactivity to fearful faces correlates positively with impulsive aggression. *Social Neuroscience*, *14*(2), 162–172. https://doi.org/10.1080/17470919.2017.1421262

da Cunha-Bang, S., Fisher, P. M., Hjordt, L. V., Perfalk, E., Skibsted, A. P., Bock, C., Baandrup, A. O., Deen, M., Thomsen, C., Sestoft, D. M., Sestoft, D. M., & Knudsen, G. M. (2017). Violent offenders respond to provocations with high amygdala and striatal reactivity. *Social Cognitive and Affective Neuroscience*, *12*(5), 802–810. https://doi.org/10.1093/scan/nsx006

da Cunha-Bang, S., McMahon, B., MacDonald Fisher, P., Steen Jensen, P., Svarer, C., & Knudsen, G. M. (2016). High trait aggression in men is associated with low 5-HT levels, as indexed by 5-HT<inf>4</inf> receptor binding. *Social Cognitive and Affective Neuroscience*, *11*(4), 548–555. https://doi.org/10.1093/scan/nsv140

Dalwani, M., Sakai, J. T., Mikulich-Gilbertson, S. K., Tanabe, J., Raymond, K., McWilliams, S. K., Thompson, L. L., Banich, M. T., & Crowley, T. J. (2011). Reduced cortical gray matter volume in male adolescents with substance and conduct problems. *Drug and Alcohol Dependence*, *118*(2–3), 295–305. https://doi.org/10.1016/j.drugalcdep.2011.04.006

Dalwani, M. S., Tregellas, J. R., Andrews-Hanna, J. R., Mikulich-Gilbertson, S. K., Raymond, K. M., Banich, M. T., Crowley, T. J., & Sakai, J. T. (2014). Default mode network activity in male adolescents with conduct and substance use disorder. *Drug and Alcohol Dependence*, *134*(1), 242–250. https://doi.org/10.1016/j.drugalcdep.2013.10.009

Dambacher, F., Sack, A. T., Lobbestael, J., Arntz, A., Brugman, S., & Schuhmann, T. (2014). Out of control: Evidence for anterior insula involvement in motor impulsivity and reactive aggression. *Social Cognitive and Affective Neuroscience*, *10*(4), 508–516. https://doi.org/10.1093/scan/nsu077

De Brito, S. A., Mechelli, A., Wilke, M., Laurens, K. R., Jones, A. P., Barker, G. J., Hodgins, S., & Viding, E. (2009). Size matters: Increased grey matter in boys with conduct problems and callousunemotional traits. *Brain*, *132*(4), 843–852. https://doi.org/10.1093/brain/awp011

de Oliveira-Souza, R., Hare, R. D., Bramati, I. E., Garrido, G. J., Azevedo Ignácio, F., Tovar-Moll, F., & Moll, J. (2008). Psychopathy as a disorder of the moral brain: Fronto-temporo-limbic grey matter reductions demonstrated by voxel-based morphometry. *NeuroImage*, *40*(3), 1202–1213. https://doi.org/10.1016/j.neuroimage.2007.12.054

Decety, J., Chen, C., Harenski, C., & Kiehl, K. A. (2013). An fMRI study of affective perspective taking in individuals with psychopathy: Imagining another in pain does not evoke empathy. *Frontiers in Human Neuroscience*, *SEP*. https://doi.org/10.3389/fnhum.2013.00489

Decety, J., Chen, C., Harenski, C. L., & Kiehl, K. A. (2015). Socioemotional processing of morally-laden behavior and their consequences on others in forensic psychopaths. *Human Brain Mapping*, *36*(6), 2015–2026. https://doi.org/10.1002/hbm.22752

Decety, J., Michalska, K. J., Akitsuki, Y., & Lahey, B. B. (2009). Atypical empathic responses in adolescents with aggressive conduct disorder: A functional MRI investigation. *Biological Psychology*, *80*(2), 203–211. https://doi.org/10.1016/j.biopsycho.2008.09.004

Decety, J., & Porges, E. C. (2011). Imagining being the agent of actions that carry different moral consequences: An fMRI study. *Neuropsychologia*, *49*(11), 2994–3001. https://doi.org/10.1016/j.neuropsychologia.2011.06.024

Decety, J., Skelly, L., Yoder, K. J., & Kiehl, K. A. (2014). Neural processing of dynamic emotional facial expressions in psychopaths. *Social Neuroscience*, *9*(1), 36–49. https://doi.org/10.1080/17470919.2013.866905

Decety, J., Skelly, L. R., & Kiehl, K. A. (2013). Brain response to empathy-eliciting scenarios involving pain in incarcerated individuals with psychopathy. *JAMA Psychiatry*, *70*(6), 638–645. https://doi.org/10.1001/jamapsychiatry.2013.27

Deeley, Q., Daly, E., Surguladze, S., Tunstall, N., Mezey, G., Beer, D., Ambikapathy, A., Robertson, D., Giampietro, V., Brammer, M. J., Phillips, M. L., & Murphy, D. G. (2006). Facial emotion processing in criminal psychopathy: Preliminary functional magnetic resonance imaging study. *British Journal of Psychiatry*, *189*(DEC.), 533–539. https://doi.org/10.1192/bjp.bp.106.021410

Delfin, C., Krona, H., Andiné, P., Ryding, E., Wallinius, M., & Hofvander, B. (2019). Prediction of recidivism in a long-term followup of forensic psychiatric patients: Incremental effects of neuroimaging data. *PLoS ONE*, *14*(5). https://doi.org/10.1371/journal.pone.0217127

Deming, P., Dargis, M., Haas, B. W., Brook, M., Decety, J., Harenski, C., Kiehl, K. A., Koenigs, M., & Kosson, D. S. (2020). Psychopathy is associated with fear-specific reductions in neural activity during affective perspective-taking. *NeuroImage*, *223*. https://doi.org/10.1016/j.neuroimage.2020.117342

Deming, P., Philippi, C. L., Wolf, R. C., Dargis, M., Kiehl, K. A., & Koenigs, M. (2018). Psychopathic traits linked to alterations in neural activity during personality judgments of self and others. *NeuroImage: Clinical*, *18*, 575–581. https://doi.org/10.1016/j.nicl.2018.02.029

Denson, T. F., Pedersen, W. C., Ronquillo, J., & Nandy, A. S. (2009). The angry brain: Neural correlates of anger, angry rumination, and aggressive personality. *Journal of Cognitive Neuroscience*, *21*(4), 737–744. https://doi.org/10.1162/jocn.2009.21051

Dolan, M. C., Deakin, J. F. W., Roberts, N., & Anderson, I. M. (2002). Quantitative frontal and temporal structural MRI studies in personality-disordered offenders and control subjects. *Psychiatry Research - Neuroimaging*, *116*(3), 133–149. https://doi.org/10.1016/S0925-4927(02)00085-9

Dolan, M. C., & Fullam, R. S. (2009). Psychopathy and Functional Magnetic Resonance Imaging Blood Oxygenation Level-Dependent Responses to Emotional Faces in Violent Patients with Schizophrenia. *Biological Psychiatry*, *66*(6), 570–577. https://doi.org/10.1016/j.biopsych.2009.03.019

Dong, D., Ming, Q., Wang, X., Yu, W., Jiang, Y., Wu, Q., Gao, Y., & Yao, S. (2017). Temporoparietal junction hypoactivity during pain-related empathy processing in adolescents with conduct disorder. *Frontiers in Psychology*, *7*(JAN). https://doi.org/10.3389/fpsyg.2016.02085

Dotterer, H. L., Hyde, L. W., Swartz, J. R., Hariri, A. R., & Williamson, D. E. (2017). Amygdala reactivity predicts adolescent antisocial behavior but not callous-unemotional traits. *Developmental Cognitive Neuroscience*, *24*, 84–92. https://doi.org/10.1016/j.dcn.2017.02.008

Dougherty, D. D., Bonab, A. A., Ottowitz, W. E., Livni, E., Alpert, N. M., Rauch, S. L., Fava, M., & Fischman, A. J. (2006). Decreased striatal D1 binding as measured using PET and [$^{11}$C]SCH 23,390 in patients with major depression with anger attacks. *Depression and Anxiety*, *23*(3), 175–177. https://doi.org/10.1002/da.20168

Emmerling, F., Schuhmann, T., Lobbestael, J., Arntz, A., Brugman, S., & Sack, A. T. (2016). The role of the insular cortex in retaliation. *PLoS ONE*, *11*(4). https://doi.org/10.1371/journal.pone.0152000

Ermer, E., Cope, L. M., Nyalakanti, P. K., Calhoun, V. D., & Kiehl, K. A. (2012). Aberrant Paralimbic gray matter in criminal Psychopathy. *Journal of Abnormal Psychology*, *121*(3), 649–658. https://doi.org/10.1037/a0026371

Ermer, E., Cope, L. M., Nyalakanti, P. K., Calhoun, V. D., & Kiehl, K. A. (2013). Aberrant paralimbic gray matter in incarcerated male adolescents with psychopathic traits. *Journal of the American Academy of Child and Adolescent Psychiatry*, *52*(1), 94-103.e3. https://doi.org/10.1016/j.jaac.2012.10.013

Ewbank, M. P., Passamonti, L., Hagan, C. C., Goodyer, I. M., Calder, A. J., & Fairchild, G. (2018). Psychopathic traits influence amygdala-anterior cingulate cortex connectivity during facial emotion processing. *Social Cognitive and Affective Neuroscience*, *13*(5), 525–534. https://doi.org/10.1093/scan/nsy019

Fahim, C., Fiori, M., Evans, A. C., & Pérusse, D. (2012). The Relationship between Social Defiance, Vindictiveness, Anger, and Brain Morphology in Eight-year-old Boys and Girls. *Social Development*, *21*(3), 592–609. https://doi.org/10.1111/j.1467-9507.2011.00644.x

Fahim, C., He, Y., Yoon, U., Chen, J., Evans, A., & Pérusse, D. (2011). Neuroanatomy of childhood disruptive behavior disorders. *Aggressive Behavior*, *37*(4), 326–337. https://doi.org/10.1002/ab.20396

Fairchild, G., Hagan, C. C., Passamonti, L., Walsh, N. D., Goodyer, I. M., & Calder, A. J. (2014). Atypical neural responses during face processing in female adolescents with conduct disorder. *Journal of the American Academy of Child and Adolescent Psychiatry*, *53*(6). https://doi.org/10.1016/j.jaac.2014.02.009

Fairchild, G., Hagan, C. C., Walsh, N. D., Passamonti, L., Calder, A. J., & Goodyer, I. M. (2013). Brain structure abnormalities in adolescent girls with conduct disorder. *Journal of Child Psychology and Psychiatry and Allied Disciplines*, *54*(1), 86–95. https://doi.org/10.1111/j.1469-7610.2012.02617.x

Fairchild, G., Passamonti, L., Hurford, G., Hagan, C. C., Von Dem Hagen, E. A. H., Van Goozen, S. H. M., Goodyer, I. M., & Calder, A. J. (2011). Brain structure abnormalities in early-onset and adolescent-onset conduct disorder. *American Journal of Psychiatry*, *168*(6), 624–633. https://doi.org/10.1176/appi.ajp.2010.10081184

Fairchild, G., Toschi, N., Hagan, C. C., Goodyer, I. M., Calder, A. J., & Passamonti, L. (2015). Cortical thickness, surface area, and folding alterations in male youths with conduct disorder and varying levels of callous-unemotional traits. *NeuroImage: Clinical*, *8*, 253–260. https://doi.org/10.1016/j.nicl.2015.04.018

Fede, S. J., Borg, J. S., Nyalakanti, P. K., Harenski, C. L., Cope, L. M., Sinnott-Armstrong, W., Koenigs, M., Calhoun, V. D., & Kiehl, K. A. (2016). Distinct neuronal patterns of positive and negative moral processing in psychopathy. *Cognitive, Affective and Behavioral Neuroscience*, *16*(6), 1074–1085. https://doi.org/10.3758/s13415-016-0454-z

Fehlbaum, L. V., Raschle, N. M., Menks, W. M., Prätzlich, M., Flemming, E., Wyss, L., Euler, F., Sheridan, M., Sterzer, P., & Stadler, C. (2018). Altered neuronal responses during an affective stroop task in adolescents with conduct disorder. *Frontiers in Psychology*, *9*(OCT). https://doi.org/10.3389/fpsyg.2018.01961

Fehr, T., Achtziger, A., Roth, G., & Struber, D. (2014). Neural correlates of the empathic perceptual processing of realistic social interaction scenarios displayed from a first-order perspective. *Brain Research*, *1583*(1), 141–158. https://doi.org/10.1016/j.brainres.2014.04.041

Finger, E. C., Marsh, A. A., Blair, K. S., Reid, M. E., Sims, C., Ng, P., Pine, D. S., & Blair, R. J. R. (2011). Disrupted reinforcement signaling in the orbitofrontal cortex and caudate in youths with conduct disorder or oppositional defiant disorder and a high level of psychopathic traits. *American Journal of Psychiatry*, *168*(2), 152–162. https://doi.org/10.1176/appi.ajp.2010.10010129

Finger, E. C., Marsh, A. A., Mitchell, D. G., Reid, M. E., Sims, C., Budhani, S., Kosson, D. S., Chen, G., Towbin, K. E., Leibenluft, E., Pine, D. S., & Blair, J. R. (2008). Abnormal ventromedial prefrontal cortex function in children with psychopathic traits during reversal learning. *Archives of General Psychiatry*, *65*(5), 586–594. https://doi.org/10.1001/archpsyc.65.5.586

Foell, J., Brislin, S. J., Strickland, C. M., Seo, D., Sabatinelli, D., & Patrick, C. J. (2016). Externalizing proneness and brain response during pre-cuing and viewing of emotional pictures. *Social Cognitive and Affective Neuroscience*, *11*(7), 1102–1110. https://doi.org/10.1093/scan/nsv080

Frankle, W. G., Lombardo, I., New, A. S., Goodman, M., Talbot, P. S., Huang, Y., Hwang, D.-R., Slifstein, M., Curry, S., Abi-Bargham, A., Laruelle, M., & Siever, L. J. (2005). Brain serotonin transporter distribution in subjects with impulsive aggressivity: A positron emission study with [$^{11}$C]McN 5652. *American Journal of Psychiatry*, *162*(5), 915–923. https://doi.org/10.1176/appi.ajp.162.5.915

Freeman, S. M., Clewett, D. V., Bennett, C. M., Kiehl, K. A., Gazzaniga, M. S., & Miller, M. B. (2015). The posteromedial region of the default mode network shows attenuated task-induced deactivation in psychopathic prisoners. *Neuropsychology*, *29*(3), 493–500. https://doi.org/10.1037/neu0000118

Fullam, R. S., McKie, S., & Dolan, M. C. (2009). Psychopathic traits and deception: Functional magnetic resonance imaging study. *British Journal of Psychiatry*, *194*(3), 229–235. https://doi.org/10.1192/bjp.bp.108.053199

Gan, G., Preston-Campbell, R. N., Moeller, S. J., Steinberg, J. L., Lane, S. D., Maloney, T., Parvaz, M. A., Goldstein, R. Z., & Alia-Klein, N. (2016). Reward vs. retaliation—the role of the mesocorticolimbic salience network in human reactive aggression. *Frontiers in Behavioral Neuroscience*, *10*(SEP). https://doi.org/10.3389/fnbeh.2016.00179

Gansler, D. A., Lee, A. K. W., Emerton, B. C., D'Amato, C., Bhadelia, R., Jerram, M., & Fulwiler, C. (2011). Prefrontal regional correlates of self-control in male psychiatric patients: Impulsivity facets and aggression. *Psychiatry Research - Neuroimaging*, *191*(1), 16–23. https://doi.org/10.1016/j.pscychresns.2010.09.003

Gansler, D. A., McLaughlin, N. C. R., Iguchi, L., Jerram, M., Moore, D. W., Bhadelia, R., & Fulwiler, C. (2009). A multivariate approach to aggression and the orbital frontal cortex in psychiatric patients. *Psychiatry Research - Neuroimaging*, *171*(3), 145–154. https://doi.org/10.1016/j.pscychresns.2008.03.007

Gao, Y., Jiang, Y., Ming, Q., Zhang, J., Ma, R., Wu, Q., Dong, D., Sun, X., He, J., Cao, W., Yuan, S., & Yao, S. (2021). Neuroanatomical changes associated with conduct disorder in boys: influence of childhood maltreatment. *European Child and Adolescent Psychiatry*. https://doi.org/10.1007/s00787-020-01697-z

Gard, A. M., Waller, R., Shaw, D. S., Forbes, E. E., Hariri, A. R., & Hyde, L. W. (2017). The Long Reach of Early Adversity: Parenting, Stress, and Neural Pathways to Antisocial Behavior in Adulthood. *Biological Psychiatry: Cognitive Neuroscience and Neuroimaging*, *2*(7), 582–590. https://doi.org/10.1016/j.bpsc.2017.06.005

George, D. T., Rawlings, R. R., Williams, W. A., Phillips, M. J., Fong, G., Kerich, M., Momenan, R., Umhau, J. C., & Hommer, D. (2004). A select group of perpetrators of domestic violence: Evidence of decreased metabolism in the right hypothalamus and reduced relationships between cortical/subcortical brain structures in position emission tomography. *Psychiatry Research - Neuroimaging*, *130*(1), 11–25. https://doi.org/10.1016/S0925-4927(03)00105-7

Gerra, G., Calbiani, B., Zaimovic, A., Sartori, R., Ugolotti, G., Ippolito, L., Delsignore, R., Rustichelli, P., & Fontanesi, B. (1998). Regional cerebral blood flow and comorbid diagnosis in abstinent opioid addicts. *Psychiatry Research - Neuroimaging*, *83*(2), 117–126. https://doi.org/10.1016/S0925-4927(98)00030-4

Geurts, D. E. M., von Borries, K., Volman, I., Bulten, B. H., Cools, R., & Verkes, R.-J. (2016). Neural connectivity during reward expectation dissociates psychopathic criminals from non-criminal individuals with high impulsive/antisocial psychopathic traits. *Social Cognitive and Affective Neuroscience*, *11*(8), 1326–1334. https://doi.org/10.1093/scan/nsw040

Glenn, A. L., Han, H., Yang, Y., Raine, A., & Schug, R. A. (2017). Associations between psychopathic traits and brain activity during instructed false responding. *Psychiatry Research - Neuroimaging*, *266*, 123–137. https://doi.org/10.1016/j.pscychresns.2017.06.008

Goldstein, R. Z., Alia-Klein, N., Leskovjan, A. C., Fowler, J. S., Wang, G.-J., Gur, R. C., Hitzemann, R., & Volkow, N. D. (2005). Anger and depression in cocaine addiction: Association with the orbitofrontal cortex. *Psychiatry Research - Neuroimaging*, *138*(1), 13–22. https://doi.org/10.1016/j.pscychresns.2004.10.002

Gopal, A., Clark, E., Allgair, A., D'Amato, C., Furman, M., Gansler, D. A., & Fulwiler, C. (2013). Dorsal/ventral parcellation of the amygdala: Relevance to impulsivity and aggression. *Psychiatry Research - Neuroimaging*, *211*(1), 24–30. https://doi.org/10.1016/j.pscychresns.2012.10.010

Gordon, H. L., Baird, A. A., & End, A. (2004). Functional differences among those high and low on a trait measure of psychopathy. *Biological Psychiatry*, *56*(7), 516–521. https://doi.org/10.1016/j.biopsych.2004.06.030

Gorka, A. X., Norman, R. E., Radtke, S. R., Carré, J. M., & Hariri, A. R. (2015). Anterior cingulate cortex gray matter volume mediates an association between 2D:4D ratio and trait aggression in women but not men. *Psychoneuroendocrinology*, *56*, 148–156. https://doi.org/10.1016/j.psyneuen.2015.03.004

Goyer, P. F., Andreason, P. J., Semple, W. E., Clayton, A. H., King, A. C., Compton-Toth, B. A., Schulz, S. C., & Cohen, R. M. (1994). Positron-emission tomography and personality disorders. *Neuropsychopharmacology*, *10*(1), 21–28. https://doi.org/10.1038/npp.1994.3

Gregory, S., Blair, R. J., ffytche, D., Simmons, A., Kumari, V., Hodgins, S., & Blackwood, N. (2015). Punishment and psychopathy: A case-control functional MRI investigation of reinforcement learning in violent antisocial personality disordered men. *The Lancet Psychiatry*, *2*(2), 153–160. https://doi.org/10.1016/S2215-0366(14)00071-6

Gregory, S., Ffytche, D., Simmons, A., Kumari, V., Howard, M., Hodgins, S., & Blackwood, N. (2012). The antisocial brain: Psychopathy matters: A structural mri investigation of antisocial male violent offenders. *Archives of General Psychiatry*, *69*(9), 962–972. https://doi.org/10.1001/archgenpsychiatry.2012.222

Han, T., Alders, G. L., Greening, S. G., Neufeld, R. W. J., & Mitchell, D. G. V. (2012). Do fearful eyes activate empathy-related brain regions in individuals with callous traits? *Social Cognitive and Affective Neuroscience*, *7*(8), 958–968. https://doi.org/10.1093/scan/nsr068

Harenski, C. L., Edwards, B. G., Harenski, K. A., & Kiehl, K. A. (2014). Neural correlates of moral and non-moral emotion in female psychopathy. *Frontiers in Human Neuroscience*, *8*(SEP), 1–10. https://doi.org/10.3389/fnhum.2014.00741

Harenski, C. L., Harenski, K. A., & Kiehl, K. A. (2014). Neural processing of moral violations among incarcerated adolescents with psychopathic traits. *Developmental Cognitive Neuroscience*, *10*, 181–189. https://doi.org/10.1016/j.dcn.2014.09.002

Harenski, C. L., Harenski, K. A., Shane, M. S., & Kiehl, K. A. (2010). Aberrant neural processing of moral violations in criminal psychopaths. *Journal of Abnormal Psychology*, *119*(4), 863–874. https://doi.org/10.1037/a0020979

Harenski, C. L., Thornton, D. M., Harenski, K. A., Decety, J., & Kiehl, K. A. (2012). Increased frontotemporal activation during pain observation in sexual sadism: Preliminary findings. *Archives of General Psychiatry*, *69*(3), 283–292. https://doi.org/10.1001/archgenpsychiatry.2011.1566

Heesink, L., Edward Gladwin, T., Terburg, D., van Honk, J., Kleber, R., & Geuze, E. (2017). Proximity alert! Distance related cuneus activation in military veterans with anger and aggression problems. *Psychiatry Research - Neuroimaging*, *266*, 114–122. https://doi.org/10.1016/j.pscychresns.2017.06.012

Heesink, L., Gladwin, T. E., Vink, M., van Honk, J., Kleber, R., & Geuze, E. (2018). Neural activity during the viewing of emotional pictures in veterans with pathological anger and aggression. *European Psychiatry*, *47*, 1–8. https://doi.org/10.1016/j.eurpsy.2017.09.002

Herpertz, S. C., Huebner, T., Marx, I., Vloet, T. D., Fink, G. R., Stoecker, T., Jon Shah, N., Konrad, K., & Herpertz-Dahlmann, B. (2008). Emotional processing in male adolescents with childhood-onset conduct disorder. *Journal of Child Psychology and Psychiatry and Allied Disciplines*, *49*(7), 781–791. https://doi.org/10.1111/j.1469-7610.2008.01905.x

Hirono, N., Mega, M. S., Dinov, I. D., Mishkin, F., & Cummings, J. L. (2000). Left frontotemporal hypoperfusion is associated with aggression in patients with dementia. *Archives of Neurology*, *57*(6), 861–866. https://doi.org/10.1001/archneur.57.6.861

Hofhansel, L., Weidler, C., Votinov, M., Clemens, B., Raine, A., & Habel, U. (2020). Morphology of the criminal brain: gray matter reductions are linked to antisocial behavior in offenders. *Brain Structure and Function*, *225*(7), 2017–2028. https://doi.org/10.1007/s00429-020-02106-6

Holz, N. E., Boecker, R., Hohm, E., Zohsel, K., Buchmann, A. F., Blomeyer, D., Jennen-Steinmetz, C., Baumeister, S., Hohmann, S., Wolf, I., Brandeis, D., & Laucht, M. (2015). The long-term impact of early life poverty on orbitofrontal cortex volume in adulthood: Results from a prospective study over 25 years. *Neuropsychopharmacology*, *40*(4), 996–1004. https://doi.org/10.1038/npp.2014.277

Hoptman, M. J., Volavka, J., Czobor, P., Gerig, G., Chakos, M., Blocher, J., Citrome, L. L., Sheitman, B., Lindenmayer, J.-P., Lieberman, J. A., Lieberman, J. A., & Bilder, R. M. (2006). Aggression and quantitative MRI measures of caudate in patients with chronic schizophrenia or schizoaffective disorder. *Journal of Neuropsychiatry and Clinical Neurosciences*, *18*(4), 509–515. https://doi.org/10.1176/jnp.2006.18.4.509

Hoptman, M. J., Volavka, J., Weiss, E. M., Czobor, P., Szeszko, P. R., Gerig, G., Chakos, M., Blocher, J., Citrome, L. L., Lindenmayer, J.-P., Lieberman, J. A., & Bilder, R. M. (2005). Quantitative MRI measures of orbitofrontal cortex in patients with chronic schizophrenia or schizoaffective disorder. *Psychiatry Research - Neuroimaging*, *140*(2), 133–145. https://doi.org/10.1016/j.pscychresns.2005.07.004

Hosking, J. G., Kastman, E. K., Dorfman, H. M., Samanez-Larkin, G. R., Baskin-Sommers, A., Kiehl, K. A., Newman, J. P., & Buckholtz, J. W. (2017). Disrupted Prefrontal Regulation of Striatal Subjective Value Signals in Psychopathy. *Neuron*, *95*(1), 221-231.e4. https://doi.org/10.1016/j.neuron.2017.06.030

Howner, K., Eskildsen, S. F., Fischer, H., Dierks, T., Wahlund, L.-O., Jonsson, T., Wiberg, M. K., & Kristiansson, M. (2012). Thinner cortex in the frontal lobes in mentally disordered offenders. *Psychiatry Research - Neuroimaging*, *203*(2–3), 126–131. https://doi.org/10.1016/j.pscychresns.2011.12.011

Huang, Y., Wu, T., Gao, Y., Luo, Y., Wu, Z., Fagan, S., Leung, S., & Li, X. (2019). The Impact of Callous-Unemotional Traits and Externalizing Tendencies on Neural Responsivity to Reward and Punishment in Healthy Adolescents. *Frontiers in Neuroscience*, *13*. https://doi.org/10.3389/fnins.2019.01319

Huber, C. G., Widmayer, S., Smieskova, R., Egloff, L., Riecher-Rössler, A., Stieglitz, R.-D., & Borgwardt, S. (2018). Voxel-Based Morphometry Correlates of an Agitated-Aggressive Syndrome in the At-Risk Mental State for Psychosis and First Episode Psychosis. *Scientific Reports*, *8*(1). https://doi.org/10.1038/s41598-018-33770-8

Huebner, T., Vloet, T. D., Marx, I., Konrad, K., Fink, G. R., Herpertz, S. C., & Herpertz-Dahlmann, B. (2008). Morphometric brain abnormalities in boys with conduct disorder. *Journal of the American Academy of Child and Adolescent Psychiatry*, *47*(5), 540–547. https://doi.org/10.1097/CHI.0b013e3181676545

Hwang, S., Meffert, H., Van Tieghem, M. R., Sinclair, S., Bookheimer, S. Y., Vaughan, B., & Blair, R. J. R. (2018). Dysfunctional social reinforcement processing in disruptive behavior disorders: An functional magnetic resonance imaging study. *Clinical Psychopharmacology and Neuroscience*, *16*(4), 449–460. https://doi.org/10.9758/cpn.2018.16.1.449

Hwang, S., Nolan, Z. T., White, S. F., Williams, W. C., Sinclair, S., & Blair, R. J. R. (2016). Dual neurocircuitry dysfunctions in disruptive behavior disorders: Emotional responding and response inhibition. *Psychological Medicine*, *46*(7), 1485–1496. https://doi.org/10.1017/S0033291716000118

Hyatt, C. J., Haney-Caron, E., & Stevens, M. C. (2012). Cortical thickness and folding deficits in conduct-disordered adolescents. *Biological Psychiatry*, *72*(3), 207–214. https://doi.org/10.1016/j.biopsych.2011.11.017

Hyde, L. W., Byrd, A. L., Votruba-Drzal, E., Hariri, A. R., & Manuck, S. B. (2014). Amygdala reactivity and negative emotionality: Divergent correlates of antisocial personality and psychopathy traits in a community sample. *Journal of Abnormal Psychology*, *123*(1), 214–224. https://doi.org/10.1037/a0035467

Ibrahim, K., Eilbott, J. A., Ventola, P., He, G., Pelphrey, K. A., McCarthy, G., & Sukhodolsky, D. G. (2019). Reduced Amygdala–Prefrontal Functional Connectivity in Children With Autism Spectrum Disorder and Co-occurring Disruptive Behavior. *Biological Psychiatry: Cognitive Neuroscience and Neuroimaging*, *4*(12), 1031–1041. https://doi.org/10.1016/j.bpsc.2019.01.009

Intrator, J., Hare, R., Stritzke, P., Brichtswein, K., Dorfman, D., Harpur, T., Bernstein, D., Handelsman, L., Schaefer, C., Keilp, J., Rosen, J., & Machac, J. (1997). A brain imaging (single photon emission computerized tomography) study of semantic and affective processing in psychopaths. *Biological Psychiatry*, *42*(2), 96–103. https://doi.org/10.1016/S0006-3223(96)00290-9

Jiang, W., Li, G., Liu, H., Shi, F., Wang, T., Shen, C., Shen, H., Lee, S.-W., Hu, D., Wang, W., Wang, W., & Shen, D. (2016). Reduced cortical thickness and increased surface area in antisocial personality disorder. *Neuroscience*, *337*, 143–152. https://doi.org/10.1016/j.neuroscience.2016.08.052

Jones, A. P., Laurens, K. R., Herba, C. M., Barker, G. J., & Viding, E. (2009). Amygdala hypoactivity to fearful faces in boys with conduct problems and callous-unemotional traits. *American Journal of Psychiatry*, *166*(1), 95–102. https://doi.org/10.1176/appi.ajp.2008.07071050

Jones, S., Joyal, C. C., Cisler, J. M., & Bai, S. (2017). Exploring Emotion Regulation in Juveniles Who Have Sexually Offended: An fMRI Study. *Journal of Child Sexual Abuse*, *26*(1), 40–57. https://doi.org/10.1080/10538712.2016.1259280

Joyal, C. C., Putkonen, A., Mancini-Marïe, A., Hodgins, S., Kononen, M., Boulay, L., Pihlajamaki, M., Soininen, H., Stip, E., Tiihonen, J., Tiihonen, J., & Aronen, H. J. (2007). Violent persons with schizophrenia and comorbid disorders: A functional magnetic resonance imaging study. *Schizophrenia Research*, *91*(1–3), 97–102. https://doi.org/10.1016/j.schres.2006.12.014

Juhász, C., Behen, M. E., Muzik, O., Chugani, D. C., & Chugani, H. T. (2001). Bilateral medial prefrontal and temporal neocortical hypometabolism in children with epilepsy and aggression. *Epilepsia*, *42*(8), 991–1001. https://doi.org/10.1046/j.1528-1157.2001.042008991.x

Kiehl, K. A., Smith, A. M., Hare, R. D., Mendrek, A., Forster, B. B., Brink, J., & Liddle, P. F. (2001). Limbic abnormalities in affective processing by criminal psychopaths as revealed by functional magnetic resonance imaging. *Biological Psychiatry*, *50*(9), 677–684. https://doi.org/10.1016/S0006-3223(01)01222-7

Kiehl, K. A., Smith, A. M., Mendrek, A., Forster, B. B., Hare, R. D., & Liddle, P. F. (2004). Temporal lobe abnormalities in semantic processing by criminal psychopaths as revealed by functional magnetic resonance imaging. *Psychiatry Research - Neuroimaging*, *130*(1), 27–42. https://doi.org/10.1016/S0925-4927(03)00106-9

Kim, M. Y., & James, L. R. (2015). Neurological evidence for the relationship between suppression and aggressive behavior: Implications for workplace aggression. *Applied Psychology*, *64*(2), 286–307. https://doi.org/10.1111/apps.12014

Kirino, E., Hayakawa, Y., Inami, R., Inoue, R., & Aoki, S. (2019). Simultaneous fMRI-EEG-DTI recording of MMN in patients with schizophrenia. *PLoS ONE*, *14*(5). https://doi.org/10.1371/journal.pone.0215023

Klapwijk, E. T., Aghajani, M., Colins, O. F., Marijnissen, G. M., Popma, A., Van Lang, N. D. J., Van Der Wee, N. J. A., & Vermeiren, R. R. J. M. (2016). Different brain responses during empathy in autism spectrum disorders versus conduct disorder and callous-unemotional traits. *Journal of Child Psychology and Psychiatry and Allied Disciplines*, *57*(6), 737–747. https://doi.org/10.1111/jcpp.12498

Klapwijk, E. T., Lelieveld, G.-J., Aghajani, M., Boon, A. E., van der Wee, N. J. A., Popma, A., Vermeiren, R. R. J. M., & Colins, O. F. (2016). Fairness decisions in response to emotions: A functional MRI study among criminal justice-involved boys with conduct disorder. *Social Cognitive and Affective Neuroscience*, *11*(4), 674–682. https://doi.org/10.1093/scan/nsv150

Klasen, M., Mathiak, K. A., Zvyagintsev, M., Sarkheil, P., Weber, R., & Mathiak, K. A. (2020). Selective reward responses to violent success events during video games. *Brain Structure and Function*, *225*(1), 57–69. https://doi.org/10.1007/s00429-019-01986-7

Kolla, N. J., Boileau, I., Karas, K., Watts, J. J., Rusjan, P., Houle, S., & Mizrahi, R. (2021). Lower amygdala fatty acid amide hydrolase in violent offenders with antisocial personality disorder: an [$^{11}$C]CURB positron emission tomography study. *Translational Psychiatry*, *11*(1). https://doi.org/10.1038/s41398-020-01144-2

Kolla, N. J., Gregory, S., Attard, S., Blackwood, N., & Hodgins, S. (2014). Disentangling possible effects of childhood physical abuse on gray matter changes in violent offenders with psychopathy. *Psychiatry Research - Neuroimaging*, *221*(2), 123–126. https://doi.org/10.1016/j.pscychresns.2013.11.008

Kolla, N. J., Matthews, B., Wilson, A. A., Houle, S., Michael Bagby, R., Links, P., Simpson, A. I., Hussain, A., & Meyer, J. H. (2015). Lower Monoamine Oxidase-A Total Distribution Volume in Impulsive and Violent Male Offenders with Antisocial Personality Disorder and High Psychopathic Traits: An $^{11C}$ Harmine Positron Emission Tomography Study. *Neuropsychopharmacology*, *40*(11), 2596–2603. https://doi.org/10.1038/npp.2015.106

Korponay, C., Pujara, M., Deming, P., Philippi, C., Decety, J., Kosson, D. S., Kieh, K. A., & Koenigs, M. (2017). Impulsive-antisocial psychopathic traits linked to increased volume and functional connectivity within prefrontal cortex. *Social Cognitive and Affective Neuroscience*, *12*(7), 1169–1178. https://doi.org/10.1093/scan/nsx042

Korponay, C., Pujara, M., Deming, P., Philippi, C., Decety, J., Kosson, D. S., Kiehl, K. A., & Koenigs, M. (2017). Impulsive-Antisocial Dimension of Psychopathy Linked to Enlargement and Abnormal Functional Connectivity of the Striatum. *Biological Psychiatry: Cognitive Neuroscience and Neuroimaging*, *2*(2), 149–157. https://doi.org/10.1016/j.bpsc.2016.07.004

Krämer, U. M., Jansma, H., Tempelmann, C., & Münte, T. F. (2007). Tit-for-tat: The neural basis of reactive aggression. *NeuroImage*, *38*(1), 203–211. https://doi.org/10.1016/j.neuroimage.2007.07.029

Kuikka, J. T., Tiihonen, J., Bergström, K. A., Karhu, J., Räsänen, P., & Eronen, M. (1998). Abnormal structure of human striatal dopamine re-uptake sites in habitually violent alcoholic offenders: A fractal analysis. *Neuroscience Letters*, *253*(3), 195–197. https://doi.org/10.1016/S0304-3940(98)00640-5

Kumari, V., Aasen, I., Taylor, P., ffytche, D. H., Das, M., Barkataki, I., Goswami, S., O'Connell, P., Howlett, M., Williams, S. C. R., Williams, S. C. R., & Sharma, T. (2006). Neural dysfunction and violence in schizophrenia: An fMRI investigation. *Schizophrenia Research*, *84*(1), 144–164. https://doi.org/10.1016/j.schres.2006.02.017

Kumari, V., Barkataki, I., Goswami, S., Flora, S., Das, M., & Taylor, P. (2009). Dysfunctional, but not functional, impulsivity is associated with a history of seriously violent behaviour and reduced orbitofrontal and hippocampal volumes in schizophrenia. *Psychiatry Research - Neuroimaging*, *173*(1), 39–44. https://doi.org/10.1016/j.pscychresns.2008.09.003

Kumari, V., Das, M., Taylor, P. J., Barkataki, I., Andrew, C., Sumich, A., Williams, S. C. R., & ffytche, D. H. (2009). Neural and behavioural responses to threat in men with a history of serious violence and schizophrenia or antisocial personality disorder. *Schizophrenia Research*, *110*(1–3), 47–58. https://doi.org/10.1016/j.schres.2009.01.009

Kumari, V., Gudjonsson, G. H., Raghuvanshi, S., Barkataki, I., Taylor, P., Sumich, A., Das, K., Kuipers, E., Ffytche, D. H., & Das, M. (2013). Reduced thalamic volume in men with antisocial personality disorder or schizophrenia and a history of serious violence and childhood abuse. *European Psychiatry*, *28*(4), 225–234. https://doi.org/10.1016/j.eurpsy.2012.03.002

Kumari, V., Uddin, S., Premkumar, P., Young, S., Gudjonsson, G. H., Raghuvanshi, S., Barkataki, I., Sumich, A., Taylor, P., & Das, M. (2014). Lower anterior cingulate volume in seriously violent men with antisocial personality disorder or schizophrenia and a history of childhood abuse. *Australian and New Zealand Journal of Psychiatry*, *48*(2), 153–161. https://doi.org/10.1177/0004867413512690

Kuroki, N., Kashiwagi, H., Ota, M., Ishikawa, M., Kunugi, H., Sato, N., Hirabayashi, N., & Ota, T. (2017). Brain structure differences among male schizophrenic patients with history of serious violent acts: An MRI voxel-based morphometric study. *BMC Psychiatry*, *17*(1). https://doi.org/10.1186/s12888-017-1263-9

Kuruoğlu, A. Ç., Arikan, Z., Vural, G., Karataş, M., Araç, M., & Işik, E. (1996). Single photon emission computerised tomography in chronic alcoholism: Antisocial personality disorder may be associated with decreased frontal perfusion. *British Journal of Psychiatry*, *169*(3), 348–354. https://doi.org/10.1192/bjp.169.3.348

Kärgel, C., Massau, C., Weiß, S., Walter, M., Borchardt, V., Krueger, T. H. C., Tenbergen, G., Kneer, J., Wittfoth, M., Pohl, A., Walter, H., & Schiffer, B. (2017). Evidence for superior neurobiological and behavioral inhibitory control abilities in non-offending as compared to offending pedophiles. *Human Brain Mapping*, *38*(2), 1092–1104. https://doi.org/10.1002/hbm.23443

Laakso, A., Wallius, E., Kajander, J., Bergman, J., Eskola, O., Solin, O., Ilonen, T., Salokangas, R. K. R., Syvälahti, E., & Hietala, J. (2003). Personality traits and striatal dopamine synthesis capacity in healthy subjects. *American Journal of Psychiatry*, *160*(5), 904–910. https://doi.org/10.1176/appi.ajp.160.5.904

Laakso, M. P., Vaurio, O., Koivisto, E., Savolainen, L., Eronen, M., Aronen, H. J., Hakola, P., Repo, E., Soininen, H., & Tiihonen, J. (2001). Psychopathy and the posterior hippocampus. *Behavioural Brain Research*, *118*(2), 187–193. https://doi.org/10.1016/S0166-4328(00)00324-7

Laakso, M. P., Gunning-Dixon, F., Vaurio, O., Repo-Tiihonen, E., Soininen, H., & Tiihonen, J. (2002). Prefrontal volumes in habitually violent subjects with antisocial personality disorder and type 2 alcoholism. *Psychiatry Research - Neuroimaging*, *114*(2), 95–102. https://doi.org/10.1016/S0925-4927(02)00005-7

Lam, B. Y. H., Yang, Y., Schug, R. A., Han, C., Liu, J., & Lee, T. M. C. (2017). Psychopathy moderates the relationship between orbitofrontal and striatal alterations and violence: The investigation of individuals accused of homicide. *Frontiers in Human Neuroscience*, *11*. https://doi.org/10.3389/fnhum.2017.00579

Larson, C. L., Baskin-Sommers, A. R., Stout, D. M., Balderston, N. L., Curtin, J. J., Schultz, D. H., Kiehl, K. A., & Newman, J. P. (2013). The interplay of attention and emotion: Top-down attention modulates amygdala activation in psychopathy. *Cognitive, Affective and Behavioral Neuroscience*, *13*(4), 757–770. https://doi.org/10.3758/s13415-013-0172-8

Lasko, E. N., Chester, D. S., Martelli, A. M., West, S. J., & Dewall, C. N. (2019). An investigation of the relationship between psychopathy and greater gray matter density in lateral prefrontal cortex. *Personality Neuroscience*, *2*. https://doi.org/10.1017/pen.2019.8

Lawrence, A. D., & Brooks, D. J. (2014). Ventral striatal dopamine synthesis capacity is associated with individual differences in behavioral disinhibition. *Frontiers in Behavioral Neuroscience*, *8*(MAR). https://doi.org/10.3389/fnbeh.2014.00086

Lee, T. M. C., Chan, S.-C., & Raine, A. (2009). Hyperresponsivity to threat stimuli in domestic violence offenders: A functional magnetic resonance imaging study. *Journal of Clinical Psychiatry*, *70*(1), 36–45. https://doi.org/10.4088/JCP.08m04143

Leutgeb, V., Leitner, M., Wabnegger, A., Klug, D., Scharmüller, W., Zussner, T., & Schienle, A. (2015). Brain abnormalities in high-risk violent offenders and their association with psychopathic traits and criminal recidivism. *Neuroscience*, *308*, 194–201. https://doi.org/10.1016/j.neuroscience.2015.09.011

Li, C.-S. R., Kosten, T. R., & Sinha, R. (2006). Antisocial personality and stress-induced brain activation in cocaine-dependent patients. *NeuroReport*, *17*(3), 243–247. https://doi.org/10.1097/01.wnr.0000199471.06487.a2

Liu, F., Shao, Y., Li, X., Liu, L., Zhao, R., Xie, B., & Qiao, Y. (2020). Volumetric Abnormalities in Violent Schizophrenia Patients on the General Psychiatric Ward. *Frontiers in Psychiatry*, *11*. https://doi.org/10.3389/fpsyt.2020.00788

Liu, J., Zubieta, J.-K., & Heitzeg, M. (2012). Sex differences in anterior cingulate cortex activation during impulse inhibition and behavioral correlates. *Psychiatry Research - Neuroimaging*, *201*(1), 54–62. https://doi.org/10.1016/j.pscychresns.2011.05.008

Lockwood, P. L., Sebastian, C. L., McCrory, E. J., Hyde, Z. H., Gu, X., De Brito, S. A., & Viding, E. (2013). Association of callous traits with reduced neural response to others' pain in children with conduct problems. *Current Biology*, *23*(10), 901–905. https://doi.org/10.1016/j.cub.2013.04.018

Lotze, M., Veit, R., Anders, S., & Birbaumer, N. (2007). Evidence for a different role of the ventral and dorsal medial prefrontal cortex for social reactive aggression: An interactive fMRI study. *NeuroImage*, *34*(1), 470–478. https://doi.org/10.1016/j.neuroimage.2006.09.028

Lozier, L. M., Cardinale, E. M., Van Meter, J. W., & Marsh, A. A. (2014). Mediation of the relationship between callous-unemotional traits and proactive aggression by amygdala response to fear among children with conduct problems. *JAMA Psychiatry*, *71*(6), 627–636. https://doi.org/10.1001/jamapsychiatry.2013.4540

Lundwall, R. A., Stephenson, K. G., Neeley-Tass, E. S., Cox, J. C., South, M., Bigler, E. D., Anderberg, E., Prigge, M. D., Hansen, B. D., Lainhart, J. E., Petrie, J. A., & Gabrielsen, T. P. (2017). Relationship between brain stem volume and aggression in children diagnosed with autism spectrum disorder. *Research in Autism Spectrum Disorders*, *34*, 44–51. https://doi.org/10.1016/j.rasd.2016.12.001

Ly, M., Motzkin, J. C., Philippi, C. L., Kirk, G. R., Newman, J. P., Kiehl, K. A., & Koenigs, M. (2012). Cortical thinning in psychopathy. *American Journal of Psychiatry*, *169*(7), 743–749. https://doi.org/10.1176/appi.ajp.2012.11111627

Mackey, S., Chaarani, B., Kan, K.-J., Spechler, P. A., Orr, C., Banaschewski, T., Barker, G., Bokde, A. L. W., Bromberg, U., Büchel, C., Althoff, R. R., & Garavan, H. (2017). Brain Regions Related to Impulsivity Mediate the Effects of Early Adversity on Antisocial Behavior. *Biological Psychiatry*, *82*(4), 275–282. https://doi.org/10.1016/j.biopsych.2015.12.027

Marín-Lahoz, J., Martínez-Horta, S., Sampedro, F., Pagonabarraga, J., Horta-Barba, A., Bejr-kasem, H., Botí, M. Á., Fernández-Bobadilla, R., Pascual-Sedano, B., Pérez-Pérez, J., Gómez-Ansón, B., & Kulisevsky, J. (2020). Measuring impulsivity in Parkinson's disease: a correlational and structural neuroimaging study using different tests. *European Journal of Neurology*, *27*(8), 1478–1486. https://doi.org/10.1111/ene.14235

Marín-Lahoz, J., Sampedro, F., Horta-Barba, A., Martínez-Horta, S., Aracil-Bolaños, I., Camacho, V., Bejr-kasem, H., Pascual-Sedano, B., Pérez-Pérez, J., Gironell, A., Carrió, I., & Kulisevsky, J. (2020). Preservation of brain metabolism in recently diagnosed Parkinson's impulse control disorders. *European Journal of Nuclear Medicine and Molecular Imaging*, *47*(9), 2165–2174. https://doi.org/10.1007/s00259-019-04664-2

Marín-Morales, A., Pérez-García, M., Catena-Martínez, A., & Verdejo-Román, J. (2021). Emotional Regulation in Male Batterers When Faced With Pictures of Intimate Partner Violence. Do They Have a Problem With Suppressing or Experiencing Emotions? *Journal of Interpersonal Violence*. https://doi.org/10.1177/0886260520985484

Marsh, A. A., & Cardinale, E. M. (2014). When psychopathy impairs moral judgments: Neural responses during judgments about causing fear. *Social Cognitive and Affective Neuroscience*, *9*(1), 3–11. https://doi.org/10.1093/scan/nss097

Marsh, A. A., Finger, E. C., Fowler, K. A., Adalio, C. J., Jurkowitz, I. T. N., Schechter, J. C., Pine, D. S., Decety, J., & Blair, R. J. R. (2013). Empathic responsiveness in amygdala and anterior cingulate cortex in youths with psychopathic traits. *Journal of Child Psychology and Psychiatry and Allied Disciplines*, *54*(8), 900–910. https://doi.org/10.1111/jcpp.12063

Marsh, A. A., Finger, E. C., Fowler, K. A., Jurkowitz, I. T. N., Schechter, J. C., Yu, H. H., Pine, D. S., & Blair, R. J. R. (2011). Reduced amygdala-orbitofrontal connectivity during moral judgments in youths with disruptive behavior disorders and psychopathic traits. *Psychiatry Research - Neuroimaging*, *194*(3), 279–286. https://doi.org/10.1016/j.pscychresns.2011.07.008

Martinelli, A., Kreifelts, B., Wildgruber, D., Bernhard, A., Ackermann, K., Freitag, C. M., & Schwenck, C. (2021). Aggression differentially modulates neural correlates of social intention attribution to benevolent, tickling and taunting laughter: An fMRI study in children and adolescents. *Social Neuroscience*, *16*(3), 303–316. https://doi.org/10.1080/17470919.2021.1908420

Martinez-Horta, S., Sampedro, F., Horta-Barba, A., Perez-Perez, J., Pagonabarraga, J., Gomez-Anson, B., & Kulisevsky, J. (2021). Structural brain correlates of irritability and aggression in early manifest Huntington's disease. *Brain Imaging and Behavior*, *15*(1), 107–113. https://doi.org/10.1007/s11682-019-00237-x

Mathews, V. P., Kronenberger, W. G., Wang, Y., Lurito, J. T., Lowe, M. J., & Dunn, D. W. (2005). Media violence exposure and frontal lobe activation measured by functional magnetic resonance imaging in aggressive and nonaggressive adolescents. *Journal of Computer Assisted Tomography*, *29*(3), 287–292. https://doi.org/10.1097/01.rct.0000162822.46958.33

Matthies, S., Rsch, N., Weber, M., Lieb, K., Philipsen, A., Tuescher, O., Ebert, D., Hennig, J., & Van Elst, L. T. (2012). Small amygdala high aggression? the role of the amygdala in modulating aggression in healthy subjects. *World Journal of Biological Psychiatry*, *13*(1), 75–81. https://doi.org/10.3109/15622975.2010.541282

Maurer, J. M., Steele, V. R., Vincent, G. M., Rao, V., Calhoun, V. D., & Kiehl, K. A. (2019). Adolescent Psychopathic Traits Negatively Relate to Hemodynamic Activity within the Basal Ganglia during Error-Related Processing. *Journal of Abnormal Child Psychology*, *47*(12), 1917–1929. https://doi.org/10.1007/s10802-019-00560-3

McCloskey, M. S., Phan, K. L., Angstadt, M., Fettich, K. C., Keedy, S., & Coccaro, E. F. (2016). Amygdala hyperactivation to angry faces in intermittent explosive disorder. *Journal of Psychiatric Research*, *79*, 34–41. https://doi.org/10.1016/j.jpsychires.2016.04.006

Meffert, H., Gazzola, V., Den Boer, J. A., Bartels, A. A. J., & Keysers, C. (2013). Reduced spontaneous but relatively normal deliberate vicarious representations in psychopathy. *Brain*, *136*(8), 2550–2562. https://doi.org/10.1093/brain/awt190

Meldrum, R. C., Trucco, E. M., Cope, L. M., Zucker, R. A., & Heitzeg, M. M. (2018). Brain activity, low self-control, and delinquency: An fMRI study of at-risk adolescents. *Journal of Criminal Justice*, *56*, 107–117. https://doi.org/10.1016/j.jcrimjus.2017.07.007

Menks, W. M., Fehlbaum, L. V., Borbás, R., Sterzer, P., Stadler, C., & Raschle, N. M. (2021). Eye gaze patterns and functional brain responses during emotional face processing in adolescents with conduct disorder. *NeuroImage: Clinical*, *29*. https://doi.org/10.1016/j.nicl.2020.102519

Mercedes Perez-Rodriguez, M., Hazlett, E. A., Rich, E. L., Ripoll, L. H., Weiner, D. M., Spence, N., Goodman, M., Koenigsberg, H. W., Siever, L. J., & New, A. S. (2012). Striatal activity in borderline personality disorder with comorbid intermittent explosive disorder: Sex differences. *Journal of Psychiatric Research*, *46*(6), 797–804. https://doi.org/10.1016/j.jpsychires.2012.02.014

Michalska, K. J., Decety, J., Zeffiro, T. A., & Lahey, B. B. (2015). Association of regional gray matter volumes in the brain with disruptive behavior disorders in male and female children. *NeuroImage: Clinical*, *7*, 252–257. https://doi.org/10.1016/j.nicl.2014.12.012

Michalska, K. J., Zeffiro, T. A., & Decety, J. (2016). Brain response to viewing others being harmed in children with conduct disorder symptoms. *Journal of Child Psychology and Psychiatry and Allied Disciplines*, *57*(4), 510–519. https://doi.org/10.1111/jcpp.12474

Miedl, S. F., Wegerer, M., Kerschbaum, H., Blechert, J., & Wilhelm, F. H. (2018). Neural activity during traumatic film viewing is linked to endogenous estradiol and hormonal contraception. *Psychoneuroendocrinology*, *87*, 20–26. https://doi.org/10.1016/j.psyneuen.2017.10.006

Mier, D., Haddad, L., Diers, K., Dressing, H., Meyer-Lindenberg, A., & Kirsch, P. (2014). Reduced embodied simulation in psychopathy. *World Journal of Biological Psychiatry*, *15*(6), 479–487. https://doi.org/10.3109/15622975.2014.902541

Miskovich, T. A., Anderson, N. E., Harenski, C. L., Harenski, K. A., Baskin-Sommers, A. R., Larson, C. L., Newman, J. P., Hanson, J. L., Stout, D. M., Koenigs, M., Kosson, D. S., & Kiehl, K. A. (2018). Abnormal cortical gyrification in criminal psychopathy. *NeuroImage: Clinical*, *19*, 876–882. https://doi.org/10.1016/j.nicl.2018.06.007

Moeller, S. J., Froböse, M. I., Konova, A. B., Misyrlis, M., Parvaz, M. A., Goldstein, R. Z., & Alia-Klein, N. (2014). Common and distinct neural correlates of inhibitory dysregulation: Stroop fMRI study of cocaine addiction and intermittent explosive disorder. *Journal of Psychiatric Research*, *58*, 55–62. https://doi.org/10.1016/j.jpsychires.2014.07.016

Mohammadi, B., Szycik, G. R., te Wildt, B., Heldmann, M., Samii, A., & Münte, T. F. (2020). Structural brain changes in young males addicted to video-gaming. *Brain and Cognition*, *139*. https://doi.org/10.1016/j.bandc.2020.105518

Molenberghs, P., Bosworth, R., Nott, Z., Louis, W. R., Smith, J. R., Amiot, C. E., Vohs, K. D., & Decety, J. (2014). The influence of group membership and individual differences in psychopathy and perspective taking on neural responses when punishing and rewarding others. *Human Brain Mapping*, *35*(10), 4989–4999. https://doi.org/10.1002/hbm.22527

Molenberghs, P., Gapp, J., Wang, B., Louis, W. R., & Decety, J. (2016). Increased Moral Sensitivity for Outgroup Perpetrators Harming Ingroup Members. *Cerebral Cortex*, *26*(1), 225–233. https://doi.org/10.1093/cercor/bhu195

Montag, C., Weber, B., Trautner, P., Newport, B., Markett, S., Walter, N. T., Felten, A., & Reuter, M. (2012). Does excessive play of violent first-person-shooter-video-games dampen brain activity in response to emotional stimuli? *Biological Psychology*, *89*(1), 107–111. https://doi.org/10.1016/j.biopsycho.2011.09.014

Morandotti, N., Dima, D., Jogia, J., Frangou, S., Sala, M., Vidovich, G. Z., Lazzaretti, M., Gambini, F., Marraffini, E., d'Allio, G., Caverzasi, E., & Brambilla, P. (2013). Childhood abuse is associated with structural impairment in the ventrolateral prefrontal cortex and aggressiveness in patients with borderline personality disorder. *Psychiatry Research - Neuroimaging*, *213*(1), 18–23. https://doi.org/10.1016/j.pscychresns.2013.02.002

Murray, L., Shaw, D. S., Forbes, E. E., & Hyde, L. W. (2017). Reward-Related Neural Correlates of Antisocial Behavior and Callous–Unemotional Traits in Young Men. *Biological Psychiatry: Cognitive Neuroscience and Neuroimaging*, *2*(4), 346–354. https://doi.org/10.1016/j.bpsc.2017.01.009

Müller, J. L., Gänßbauer, S., Sommer, M., Döhnel, K., Weber, T., Schmidt-Wilcke, T., & Hajak, G. (2008). Gray matter changes in right superior temporal gyrus in criminal psychopaths. Evidence from voxel-based morphometry. *Psychiatry Research - Neuroimaging*, *163*(3), 213–222. https://doi.org/10.1016/j.pscychresns.2007.08.010

Müller, J. L., Sommer, M., Wagner, V., Lange, K., Taschler, H., Röder, C. H., Schuierer, G., Klein, H. E., & Hajak, G. (2003). Abnormalities in emotion processing within cortical and subcortical regions in criminal psychopaths: Evidence from a functional magnetic resonance imaging study using pictures with emotional content. *Biological Psychiatry*, *54*(2), 152–162. https://doi.org/10.1016/S0006-3223(02)01749-3

Müller, J. L., Sommer, M., Döhnel, K., Weber, T., Schmidt-Wilcke, T., & Hajak, G. (2008). Disturbed Prefrontal and Temporal Brain Function During Emotion and Cognition Interaction in Criminal Psychopathy. *Behavioral Sciences and the Law*, *26*, 131–150. 10.1002/bsl.796

Naaijen, J., Mulder, L. M., Ilbegi, S., de Bruijn, S., Kleine-Deters, R., Dietrich, A., Hoekstra, P. J., Marsman, J.-B. C., Aggensteiner, P. M., Holz, N. E., Zwiers, M. P., & Buitelaar, J. K. (2020). Specific cortical and subcortical alterations for reactive and proactive aggression in children and adolescents with disruptive behavior. *NeuroImage: Clinical*, *27*. https://doi.org/10.1016/j.nicl.2020.102344

Nakano, S., Asada, T., Yamashita, F., Kitamura, N., Matsuda, H., Hirai, S., & Yamada, T. (2006). Relationship between antisocial behavior and regional cerebral blood flow in frontotemporal dementia. *NeuroImage*, *32*(1), 301–306. https://doi.org/10.1016/j.neuroimage.2006.02.040

Narayan, V. M., Narr, K. L., Kumari, V., Woods, R. P., Thompson, P. M., Toga, A. W., & Sharma, T. (2007). Regional cortical thinning in subjects with violent antisocial personality disorder or schizophrenia. *American Journal of Psychiatry*, *164*(9), 1418–1427. https://doi.org/10.1176/appi.ajp.2007.06101631

Navalpotro-Gomez, I., Dacosta-Aguayo, R., Molinet-Dronda, F., Martin-Bastida, A., Botas-Peñin, A., Jimenez-Urbieta, H., Delgado-Alvarado, M., Gago, B., Quiroga-Varela, A., & Rodriguez-Oroz, M. C. (2019). Nigrostriatal dopamine transporter availability, and its metabolic and clinical correlates in Parkinson's disease patients with impulse control disorders. *European Journal of Nuclear Medicine and Molecular Imaging*, *46*(10), 2065–2076. https://doi.org/10.1007/s00259-019-04396-3

New, A. S., Hazlett, E. A., Buchsbaum, M. S., Goodman, M., Mitelman, S. A., Newmark, R., Trisdorfer, R., Haznedar, M. M., Koenigsberg, H. W., Flory, J., Flory, J., & Siever, L. J. (2007). Amygdala-prefrontal disconnection in borderline personality disorder. *Neuropsychopharmacology*, *32*(7), 1629–1640. https://doi.org/10.1038/sj.npp.1301283

New, A. S., Hazlett, E. A., Newmark, R. E., Zhang, J., Triebwasser, J., Meyerson, D., Lazarus, S., Trisdorfer, R., Goldstein, K. E., Goodman, M., Siever, L. J., & Buchsbaum, M. S. (2009). Laboratory Induced Aggression: A Positron Emission Tomography Study of Aggressive Individuals with Borderline Personality Disorder. *Biological Psychiatry*, *66*(12), 1107–1114. https://doi.org/10.1016/j.biopsych.2009.07.015

Oberlin, B. G., Dzemidzic, M., Bragulat, V., Lehigh, C. A., Talavage, T., O'Connor, S. J., & Kareken, D. A. (2012). Limbic responses to reward cues correlate with antisocial trait density in heavy drinkers. *NeuroImage*, *60*(1), 644–652. https://doi.org/10.1016/j.neuroimage.2011.12.043

Oder, W., Goldenberg, G., Spatt, J., Podreka, I., Binder, H., & Deecke, L. (1992). Behavioural and psychosocial sequelae of severe closed head injury and regional cerebral blood flow: A SPECT study. *Journal of Neurology Neurosurgery and Psychiatry*, *55*(6), 475–480. https://doi.org/10.1136/jnnp.55.6.475

Oquendo, M. A., Krunic, A., Parsey, R. V., Milak, M., Malone, K. M., Anderson, A., Van Heertum, R. L., & Mann, J. J. (2005). Positron emission tomography of regional brain metabolic responses to a serotonergic challenge in major depressive disorder with and without borderline personality disorder. *Neuropsychopharmacology*, *30*(6), 1163–1172. https://doi.org/10.1038/sj.npp.1300689

Osumi, T., Nakao, T., Kasuya, Y., Shinoda, J., Yamada, J., & Ohira, H. (2012). Amygdala dysfunction attenuates frustration-induced aggression in psychopathic individuals in a non-criminal population. *Journal of Affective Disorders*, *142*(1–3), 331–338. https://doi.org/10.1016/j.jad.2012.05.012

Overgaauw, S., Jansen, M., Korbee, N. J., & de Bruijn, E. R. A. (2019). Neural mechanisms involved in social conformity and psychopathic traits: Prediction errors, reward processing and saliency. *Frontiers in Behavioral Neuroscience*, *13*. https://doi.org/10.3389/fnbeh.2019.00160

Pagliaccio, D., Wiggins, J. L., Adleman, N. E., Curhan, A., Zhang, S., Towbin, K. E., Brotman, M. A., Pine, D. S., & Leibenluft, E. (2017). Behavioral and Neural Sustained Attention Deficits in Disruptive Mood Dysregulation Disorder and Attention-Deficit/Hyperactivity Disorder. *Journal of the American Academy of Child and Adolescent Psychiatry*, *56*(5), 426–435. https://doi.org/10.1016/j.jaac.2017.02.008

Pardini, D. A., & Phillips, M. (2010). Neural responses to emotional and neutral facial expressions in chronically violent men. *Journal of Psychiatry and Neuroscience*, *35*(6), 390–398. https://doi.org/10.1503/jpn.100037

Pardini, D. A., Raine, A., Erickson, K., & Loeber, R. (2014). Lower amygdala volume in men is associated with childhood aggression, early psychopathic traits, and future violence. *Biological Psychiatry*, *75*(1), 73–80. https://doi.org/10.1016/j.biopsych.2013.04.003

Parsey, R. V., Oquendo, M. A., Simpson, N. R., Ogden, R. T., Van Heertum, R., Arango, V., & Mann, J. J. (2002). Effects of sex, age, and aggressive traits in man on brain serotonin 5-HT<inf>1A</inf> receptor binding potential measured by PET using [C-11]WAY-100635. *Brain Research*, *954*(2), 173–182. https://doi.org/10.1016/S0006-8993(02)03243-2

Passamonti, L., Fairchild, G., Goodyer, I. M., Hurford, G., Hagan, C. C., Rowe, J. B., & Calder, A. J. (2010). Neural abnormalities in early-onset and adolescence-onset conduct disorder. *Archives of General Psychiatry*, *67*(7), 729–738. https://doi.org/10.1001/archgenpsychiatry.2010.75

Pawliczek, C. M., Derntl, B., Kellermann, T., Gur, R. C., Schneider, F., & Habel, U. (2013). Anger under Control: Neural Correlates of Frustration as a Function of Trait Aggression. *PLoS ONE*, *8*(10). https://doi.org/10.1371/journal.pone.0078503

Pawliczek, C. M., Derntl, B., Kellermann, T., Kohn, N., Gur, R. C., & Habel, U. (2013). Inhibitory control and trait aggression: Neural and behavioral insights using the emotional stop signal task. *NeuroImage*, *79*, 264–274. https://doi.org/10.1016/j.neuroimage.2013.04.104

Payer, D. E., Baicy, K., Lieberman, M. D., & London, E. D. (2012). Overlapping neural substrates between intentional and incidental down-regulation of negative emotions. *Emotion (Washington, D.C.)*, *12*(2), 229–235. https://doi.org/10.1037/a0027421

Payer, D. E., Lieberman, M. D., & London, E. D. (2011). Neural correlates of affect processing and aggression in methamphetamine dependence. *Archives of General Psychiatry*, *68*(3), 271–282. https://doi.org/10.1001/archgenpsychiatry.2010.154

Paz-Alonso, P. M., Navalpotro-Gomez, I., Boddy, P., Dacosta-Aguayo, R., Delgado-Alvarado, M., Quiroga-Varela, A., Jimenez-Urbieta, H., Carreiras, M., & Rodriguez-Oroz, M. C. (2020). Functional inhibitory control dynamics in impulse control disorders in Parkinson's disease. *Movement Disorders*, *35*(2), 316–325. https://doi.org/10.1002/mds.27885

Pellicano, C., Niccolini, F., Wu, K., O'Sullivan, S. S., Lawrence, A. D., Lees, A. J., Piccini, P., & Politis, M. (2015). Morphometric changes in the reward system of Parkinson's disease patients with impulse control disorders. *Journal of Neurology*, *262*(12), 2653–2661. https://doi.org/10.1007/s00415-015-7892-3

Pera-Guardiola, V., Contreras-Rodríguez, O., Batalla, I., Kosson, D., Menchón, J. M., Pifarré, J., Bosque, J., Cardoner, N., & Soriano-Mas, C. (2016). Brain structural correlates of emotion recognition in psychopaths. *PLoS ONE*, *11*(5). https://doi.org/10.1371/journal.pone.0149807

Perino, M. T., Guassi Moreira, J. F., & Telzer, E. H. (2019). Links between adolescent bullying and neural activation to viewing social exclusion. *Cognitive, Affective and Behavioral Neuroscience*, *19*(6), 1467–1478. https://doi.org/10.3758/s13415-019-00739-7

Pietrini, P., Guazzelli, M., Basso, G., Jaffe, K., & Grafman, J. (2000). Neural correlates of imaginal aggressive behavior assessed by positron emission tomography in healthy subjects. *American Journal of Psychiatry*, *157*(11), 1772–1781. https://doi.org/10.1176/appi.ajp.157.11.1772

Porges, E. C., & Decety, J. (2013). Violence as a source of pleasure or displeasure is associated with specific functional connectivity with the nucleus accumbens. *Frontiers in Human Neuroscience*, *JUL*. https://doi.org/10.3389/fnhum.2013.00447

Prehn, K., Schlagenhauf, F., Schulze, L., Berger, C., Vohs, K., Fleischer, M., Hauenstein, K., Keiper, P., Domes, G., & Herpertz, S. C. (2013). Neural correlates of risk taking in violent criminal offenders characterized by emotional hypo- and hyper-reactivity. *Social Neuroscience*, *8*(2), 136–147. https://doi.org/10.1080/17470919.2012.686923

Prehn, K., Schulze, L., Rossmann, S., Berger, C., Vohs, K., Fleischer, M., Hauenstein, K., Keiper, P., Domes, G., & Herpertz, S. C. (2013). Effects of emotional stimuli on working memory processes in male criminal offenders with borderline and antisocial personality disorder. *World Journal of Biological Psychiatry*, *14*(1), 71–78. https://doi.org/10.3109/15622975.2011.584906

Premi, E., Pilotto, A., Garibotto, V., Bigni, B., Turrone, R., Alberici, A., Cottini, E., Poli, L., Bianchi, M., Formenti, A., Borroni, B., & Padovani, A. (2016). Impulse control disorder in PD: A lateralized monoaminergic frontostriatal disconnection syndrome? *Parkinsonism and Related Disorders*, *30*, 62–66. https://doi.org/10.1016/j.parkreldis.2016.05.028

Pujara, M., Motzkin, J. C., Newman, J. P., Kiehl, K. A., & Koenigs, M. (2013). Neural correlates of reward and loss sensitivity in psychopathy. *Social Cognitive and Affective Neuroscience*, *9*(6), 794–801. https://doi.org/10.1093/scan/nst054

Pujol, J., Batalla, I., Contreras-Rodríguez, O., Harrison, B. J., Pera, V., Hernández-Ribas, R., Real, E., Bosa, L., Soriano-Mas, C., Deus, J., Menchón, J. M., & Cardoner, N. (2012). Breakdown in the brain network subserving moral judgment in criminal psychopathy. *Social Cognitive and Affective Neuroscience*, *7*(8), 917–923. https://doi.org/10.1093/scan/nsr075

Puri, B. K., Counsell, S. J., Saeed, N., Bustos, M. G., Treasaden, I. H., & Bydder, G. M. (2008). Regional grey matter volumetric changes in forensic schizophrenia patients: An MRI study comparing the brain structure of patients who have seriously and violently offended with that of patients who have not. *BMC Psychiatry*, *8*(SUPPL. 1). https://doi.org/10.1186/1471-244X-8-S1-S6

Qiao, Y., Mei, Y., Du, X., Xie, B., & Shao, Y. (2016). Reduced brain activation in violent adolescents during response inhibition. *Scientific Reports*, *6*. https://doi.org/10.1038/srep21318

Qiao, Y., Xie, B., & Du, X. (2012). Abnormal response to emotional stimulus in male adolescents with violent behavior in China. *European Child and Adolescent Psychiatry*, *21*(4), 193–198. https://doi.org/10.1007/s00787-012-0252-2

Quan, F., Zhu, W., Dong, Y., Qiu, J., Gong, X., Xiao, M., Zheng, Y., Zhao, Y., Chen, X., & Xia, L.-X. (2019). Brain structure links trait hostile attribution bias and attitudes toward violence. *Neuropsychologia*, *125*, 42–50. https://doi.org/10.1016/j.neuropsychologia.2019.01.015

Raine, A., Buchsbaum, M., & LaCasse, L. (1997). Brain abnormalities in murderers indicated by positron emission tomography. *Biological Psychiatry*, *42*(6), 495–508. https://doi.org/10.1016/S0006-3223(96)00362-9

Raine, A., Buchsbaum, M. S., Stanley, J., Lottenberg, S., Abel, L., & Stoddard, J. (1994). Selective reductions in prefrontal glucose metabolism in murderers. *Biological Psychiatry*, *36*(6), 365–373. https://doi.org/10.1016/0006-3223(94)91211-4

Raine, A., Lencz, T., Bihrle, S., LaCasse, L., & Colletti, P. (2000). Reduced prefrontal gray matter volume and reduced autonomic activity in antisocial personality disorder. *Archives of General Psychiatry*, *57*(2), 119–127. https://doi.org/10.1001/archpsyc.57.2.119

Raine, A., Meloy, J. R., Bihrle, S., Stoddard, J., LaCasse, L., & Buchsbaum, M. S. (1998). Reduced prefrontal and increased subcortical brain functioning assessed using positron emission tomography in predatory and affective murderers. *Behavioral Sciences and the Law*, *16*(3), 319–332. https://doi.org/10.1002/(SICI)1099-0798(199822)16:3<319::AID-BSL311>3.0.CO;2-G

Raine, A., Yang, Y., Narr, K. L., & Toga, A. W. (2011). Sex differences in orbitofrontal gray as a partial explanation for sex differences in antisocial personality. *Molecular Psychiatry*, *16*(2), 227–236. https://doi.org/10.1038/mp.2009.136

Rao, H., Mamikonyan, E., Detre, J. A., Siderowf, A. D., Stern, M. B., Potenza, M. N., & Weintraub, D. (2010). Decreased ventral striatal activity with impulse control disorders in Parkinson's Disease. *Movement Disorders*, *25*(11), 1660–1669. https://doi.org/10.1002/mds.23147

Raschle, N. M., Fehlbaum, L. V., Menks, W. M., Martinelli, A., Prätzlich, M., Bernhard, A., Ackermann, K., Freitag, C., De Brito, S., Fairchild, G., Fairchild, G., & Stadler, C. (2019). Atypical Dorsolateral Prefrontal Activity in Female Adolescents With Conduct Disorder During Effortful Emotion Regulation. *Biological Psychiatry: Cognitive Neuroscience and Neuroimaging*, *4*(11), 984–994. https://doi.org/10.1016/j.bpsc.2019.05.003

Raschle, N. M., Menks, W. M., Fehlbaum, L. V., Steppan, M., Smaragdi, A., Gonzalez-Madruga, K., Rogers, J., Clanton, R., Kohls, G., Martinelli, A., De Brito, S. A., & Stadler, C. (2018). Callous-unemotional traits and brain structure: Sex-specific effects in anterior insula of typically-developing youths. *NeuroImage: Clinical*, *17*, 856–864. https://doi.org/10.1016/j.nicl.2017.12.015

Regenbogen, C., Herrmann, M., & Fehr, T. (2010). The neural processing of voluntary completed, real and virtual violent and nonviolent computer game scenarios displaying predefined actions in gamers and nongamers. *Social Neuroscience*, *5*(2), 221–240. https://doi.org/10.1080/17470910903315989

Reniers, R. L. E. P., Corcoran, R., Völlm, B. A., Mashru, A., Howard, R., & Liddle, P. F. (2012). Moral decision-making, ToM, empathy and the default mode network. *Biological Psychology*, *90*(3), 202–210. https://doi.org/10.1016/j.biopsycho.2012.03.009

Repple, J., Habel, U., Wagels, L., Pawliczek, C. M., Schneider, F., & Kohn, N. (2018). Sex differences in the neural correlates of aggression. *Brain Structure and Function*, *223*(9), 4115–4124. https://doi.org/10.1007/s00429-018-1739-5

Repple, J., Pawliczek, C. M., Voss, B., Siegel, S., Schneider, F., Kohn, N., & Habel, U. (2017). From provocation to aggression: The neural network. *BMC Neuroscience*, *18*(1). https://doi.org/10.1186/s12868-017-0390-z

Rijsdijsk, F. V., Viding, E., De Brito, S., Forgiarini, M., Mechelli, A., Jones, A. P., & McCrory, E. (2010). Heritable variations in gray matter concentration as a potential endophenotype for psychopathic traits. *Archives of General Psychiatry*, *67*(4), 406–413. https://doi.org/10.1001/archgenpsychiatry.2010.20

Rilling, J. K., Glenn, A. L., Jairam, M. R., Pagnoni, G., Goldsmith, D. R., Elfenbein, H. A., & Lilienfeld, S. O. (2007). Neural Correlates of Social Cooperation and Non-Cooperation as a Function of Psychopathy. *Biological Psychiatry*, *61*(11), 1260–1271. https://doi.org/10.1016/j.biopsych.2006.07.021

Rodman, A. M., Kastman, E. K., Dorfman, H. M., Baskin-Sommers, A. R., Kiehl, K. A., Newman, J. P., & Buckholtz, J. W. (2016). Selective mapping of psychopathy and externalizing to dissociable circuits for inhibitory self-control. *Clinical Psychological Science*, *4*(3), 559–571. https://doi.org/10.1177/2167702616631495

Rosell, D. R., Thompson, J. L., Slifstein, M., Xu, X., Frankle, W. G., New, A. S., Goodman, M., Weinstein, S. R., Laruelle, M., Abi-Dargham, A., Abi-Dargham, A., & Siever, L. J. (2010). Increased Serotonin 2A Receptor Availability in the Orbitofrontal Cortex of Physically Aggressive Personality Disordered Patients. *Biological Psychiatry*, *67*(12), 1154–1162. https://doi.org/10.1016/j.biopsych.2010.03.013

Rosenthal-Von Der Pütten, A. M., Schulte, F. P., Eimler, S. C., Sobieraj, S., Hoffmann, L., Maderwald, S., Brand, M., & Krämer, N. C. (2014). Investigations on empathy towards humans and robots using fMRI. *Computers in Human Behavior*, *33*, 201–212. https://doi.org/10.1016/j.chb.2014.01.004

Rubia, K., Halari, R., Cubillo, A., Mohammad, A.-M., Scott, S., & Brammer, M. (2010). Disorder-specific inferior prefrontal hypofunction in boys with pure attention-deficit/hyperactivity disorder compared to boys with pure conduct disorder during cognitive flexibility. *Human Brain Mapping*, *31*(12), 1823–1833. https://doi.org/10.1002/hbm.20975

Rubia, K., Halari, R., Smith, A. B., Mohammad, M., Scott, S., & Brammer, M. J. (2009). Shared and disorder-specific prefrontal abnormalities in boys with pure attention-deficit/hyperactivity disorder compared to boys with pure CD during interference inhibition and attention allocation. *Journal of Child Psychology and Psychiatry and Allied Disciplines*, *50*(6), 669–678. https://doi.org/10.1111/j.1469-7610.2008.02022.x

Rubia, K., Halari, R., Smith, A. B., Mohammed, M., Scott, S., Giampietro, V., Taylor, E., & Brammer, M. J. (2008). Dissociated functional brain abnormalities of inhibition in boys with pure conduct disorder and in boys with pure attention deficit hyperactivity disorder. *American Journal of Psychiatry*, *165*(7), 889–897. https://doi.org/10.1176/appi.ajp.2008.07071084

Rubia, K., Smith, A. B., Halari, R., Matsukura, F., Mohammad, M., Taylor, E., & Brammer, M. J. (2009). Disorder-specific dissociation of orbitofrontal dysfunction in boys with pure conduct disorder during reward and ventrolateral prefrontal dysfunction in boys with pure ADHD during sustained attention. *American Journal of Psychiatry*, *166*(1), 83–94. https://doi.org/10.1176/appi.ajp.2008.08020212

Rylands, A. J., Hinz, R., Jones, M., Holmes, S. E., Feldmann, M., Brown, G., McMahon, A. W., & Talbot, P. S. (2012). Pre- and postsynaptic serotonergic differences in males with extreme levels of impulsive aggression without callous unemotional traits: A positron emission tomography study using $^{11}$C-DASB and $^{11}$C-MDL100907. *Biological Psychiatry*, *72*(12), 1004–1011. https://doi.org/10.1016/j.biopsych.2012.06.024

Sadeh, N., Spielberg, J. M., Heller, W., Herrington, J. D., Engels, A. S., Warren, S. L., Crocker, L. D., Sutton, B. P., & Miller, G. A. (2013). Emotion disrupts neural activity during selective attention in psychopathy. *Social Cognitive and Affective Neuroscience*, *8*(3), 235–246. https://doi.org/10.1093/scan/nsr092

Sajous-Turner, A., Anderson, N. E., Widdows, M., Nyalakanti, P., Harenski, K., Harenski, C., Koenigs, M., Decety, J., & Kiehl, K. A. (2020). Aberrant brain gray matter in murderers. *Brain Imaging and Behavior*, *14*(5), 2050–2061. https://doi.org/10.1007/s11682-019-00155-y

Sakai, J. T., Dalwani, M. S., Mikulich-Gilbertson, S. K., Raymond, K., McWilliams, S., Tanabe, J., Rojas, D., Regner, M., Banich, M. T., & Crowley, T. J. (2017). Imaging decision about whether to benefit self by harming others: Adolescents with conduct and substance problems, with or without callous-unemotionality, or developing typically. *Psychiatry Research - Neuroimaging*, *263*, 103–112. https://doi.org/10.1016/j.pscychresns.2017.03.004

Sala, M., Caverzasi, E., Lazzaretti, M., Morandotti, N., De Vidovich, G., Marraffini, E., Gambini, F., Isola, M., De Bona, M., Rambaldelli, G., d'Allio, G., Barale, F., Zappoli, F., & Brambilla, P. (2011). Dorsolateral prefrontal cortex and hippocampus sustain impulsivity and aggressiveness in borderline personality disorder. *Journal of Affective Disorders*, *131*, 417–421.

Sarkar, S., Daly, E., Feng, Y., Ecker, C., Craig, M. C., Harding, D., Deeley, Q., & Murphy, D. G. M. (2015). Reduced cortical surface area in adolescents with conduct disorder. *European Child and Adolescent Psychiatry*, *24*(8), 909–917. https://doi.org/10.1007/s00787-014-0639-3

Schienle, A., Wabnegger, A., Leitner, M., & Leutgeb, V. (2017). Neuronal correlates of personal space intrusion in violent offenders. *Brain Imaging and Behavior*, *11*(2), 454–460. https://doi.org/10.1007/s11682-016-9526-5

Schiffer, B., Amelung, T., Pohl, A., Kaergel, C., Tenbergen, G., Gerwinn, H., Mohnke, S., Massau, C., Matthias, W., Weiß, S., Schiltz, K., & Walter, H. (2017). Gray matter anomalies in pedophiles with and without a history of child sexual offending. *Translational Psychiatry*, *7*(5). https://doi.org/10.1038/tp.2017.96

Schiffer, B., Leygraf, N., Müller, B. W., Scherbaum, N., Forsting, M., Wiltfang, J., Gizewski, E. R., & Hodgins, S. (2013). Structural brain alterations associated with schizophrenia preceded by conduct disorder: A common and distinct subtype of schizophrenia? *Schizophrenia Bulletin*, *39*(5), 1115–1128. https://doi.org/10.1093/schbul/sbs115

Schiffer, B., Müller, B. W., Scherbaum, N., Hodgins, S., Forsting, M., Wiltfang, J., Gizewski, E. R., & Leygraf, N. (2011). Disentangling structural brain alterations associated with violent behavior from those associated with substance use disorders. *Archives of General Psychiatry*, *68*(10), 1039–1049. https://doi.org/10.1001/archgenpsychiatry.2011.61

Schiffer, B., Pawliczek, C., Mu̇̈ller, B., Forsting, M., Gizewski, E., Leygraf, N., & Hodgins, S. (2014). Neural mechanisms underlying cognitive control of men with lifelong antisocial behavior. *Psychiatry Research - Neuroimaging*, *222*(1–2), 43–51. https://doi.org/10.1016/j.pscychresns.2014.01.008

Schiffer, B., Pawliczek, C., Müller, B. W., Wiltfang, J., Brüne, M., Forsting, M., Gizewski, E. R., Leygraf, N., & Hodgins, S. (2017). Neural Mechanisms Underlying Affective Theory of Mind in Violent Antisocial Personality Disorder and/or Schizophrenia. *Schizophrenia Bulletin*, *43*(6), 1229–1239. https://doi.org/10.1093/schbul/sbx012

Schlüter, T., Winz, O., Henkel, K., Prinz, S., Rademacher, L., Schmaljohann, J., Dautzenberg, K., Cumming, P., Kumakura, Y., Rex, S., Gründer, G., & Vernaleken, I. (2013). The impact of dopamine on aggression: An [$^{18}$F]-FDOPA PET study in healthy males. *Journal of Neuroscience*, *33*(43), 16889–16896. https://doi.org/10.1523/JNEUROSCI.1398-13.2013

Schneider, F., Habel, U., Kessler, C., Posse, S., Grodd, W., & Müller-Gärtner, H.-W. (2000). Functional imaging of conditioned aversive emotional responses in antisocial personality disorder. *Neuropsychobiology*, *42*(4), 192–201. https://doi.org/10.1159/000026693

Schoretsanitis, G., Stegmayer, K., Razavi, N., Federspiel, A., Müller, T. J., Horn, H., Wiest, R., Strik, W., & Walther, S. (2019). Inferior frontal gyrus gray matter volume is associated with aggressive behavior in schizophrenia spectrum disorders. *Psychiatry Research - Neuroimaging*, *290*, 14–21. https://doi.org/10.1016/j.pscychresns.2019.06.003

Schultz, D. H., Balderston, N. L., Baskin-Sommers, A. R., Larson, C. L., & Helmstetter, F. J. (2016). Psychopaths show enhanced amygdala activation during fear conditioning. *Frontiers in Psychology*, *7*(MAR). https://doi.org/10.3389/fpsyg.2016.00348

Schulz, S. C., Camchong, J., Romine, A., Schlesinger, A., Kuskowski, M., Pardo, J. V., Cullen, K. R., & Lim, K. O. (2013). An exploratory study of the relationship of symptom domains and diagnostic severity to PET scan imaging in borderline personality disorder. *Psychiatry Research: Neuroimaging*, *214*, 161–168. https://doi.org/10.1016/j.pscychresns.2013.05.007

Schwenck, C., Ciaramidaro, A., Selivanova, M., Tournay, J., Freitag, C. M., & Siniatchkin, M. (2017). Neural correlates of affective empathy and reinforcement learning in boys with conduct problems: fMRI evidence from a gambling task. *Behavioural Brain Research*, *320*, 75–84. https://doi.org/10.1016/j.bbr.2016.11.037

Seara-Cardoso, A., Sebastian, C. L., McCrory, E., Foulkes, L., Buon, M., Roiser, J. P., & Viding, E. (2016). Anticipation of guilt for everyday moral transgressions: The role of the anterior insula and the influence of interpersonal psychopathic traits. *Scientific Reports*, *6*. https://doi.org/10.1038/srep36273

Seara-Cardoso, A., Sebastian, C. L., Viding, E., & Roiser, J. P. (2016). Affective resonance in response to others' emotional faces varies with affective ratings and psychopathic traits in amygdala and anterior insula. *Social Neuroscience*, *11*(2), 140–152. https://doi.org/10.1080/17470919.2015.1044672

Seara-Cardoso, A., Viding, E., Lickley, R. A., & Sebastian, C. L. (2015). Neural responses to others' pain vary with psychopathic traits in healthy adult males. *Cognitive, Affective and Behavioral Neuroscience*, *15*(3), 578–588. https://doi.org/10.3758/s13415-015-0346-7

Sebastian, C. L., De Brito, S. A., McCrory, E. J., Hyde, Z. H., Lockwood, P. L., Cecil, C. A. M., & Viding, E. (2016). Grey Matter Volumes in Children with Conduct Problems and Varying Levels of Callous-Unemotional Traits. *Journal of Abnormal Child Psychology*, *44*(4), 639–649. https://doi.org/10.1007/s10802-015-0073-0

Sebastian, C. L., McCrory, E. J., Dadds, M. R., Cecil, C. A. M., Lockwood, P. L., Hyde, Z. H., De Brito, S. A., & Viding, E. (2014). Neural responses to fearful eyes in children with conduct problems and varying levels of callous-unemotional traits. *Psychological Medicine*, *44*(1), 99–109. https://doi.org/10.1017/S0033291713000482

Sebastian, C. L., McCrory, E. J. P., Cecil, C. A. M., Lockwood, P. L., De Brito, S. A., Fontaine, N. M. G., & Viding, E. (2012). Neural responses to affective and cognitive theory of mind in children with conduct problems and varying levels of callous-unemotional traits. *Archives of General Psychiatry*, *69*(8), 814–822. https://doi.org/10.1001/archgenpsychiatry.2011.2070

Sebastian, C. L., Stafford, J., McCrory, E. J., Sethi, A., De Brito, S. A., Lockwood, P. L., & Viding, E. (2021). Modulation of Amygdala Response by Cognitive Conflict in Adolescents with Conduct Problems and Varying Levels of CU Traits. *Research on Child and Adolescent Psychopathology*, *49*(8), 1043–1054. https://doi.org/10.1007/s10802-021-00787-z

Seidenwurm, D., Pounds, T. R., Globus, A., & Valk, P. E. (1997). Abnormal temporal lobe metabolism in violent subjects: Correlation of imaging and neuropsychiatric findings. *American Journal of Neuroradiology*, *18*(4), 625–631.

Sekine, Y., Ouchi, Y., Takei, N., Yoshikawa, E., Nakamura, K., Futatsubashi, M., Okada, H., Minabe, Y., Suzuki, K., Iwata, Y., Iyo, M., & Mori, N. (2006). Brain serotonin transporter density and aggression in abstinent methamphetamine abusers. *Archives of General Psychiatry*, *63*(1), 90–100. https://doi.org/10.1001/archpsyc.63.1.90

Seleem, M. A., El-Shafey, R., Shahin, L. T., Abdel-Aziz, L. E., Elkonaisy, N. M., Marey, Y. K., Rizkallah, M., & Baghdadi, M. (2020). Volumetric brain abnormalities in adolescents with conduct disorder with and without attention deficit-hyperactivity disorder: a case control study. *Middle East Current Psychiatry*, *27*(1). https://doi.org/10.1186/s43045-020-00025-0

Seo, D., Lacadie, C. M., & Sinha, R. (2016). Neural Correlates and Connectivity Underlying Stress-Related Impulse Control Difficulties in Alcoholism. *Alcoholism: Clinical and Experimental Research*, *40*(9), 1884–1894. https://doi.org/10.1111/acer.13166

Seok, J.-W., & Cheong, C. (2020). Gray Matter Deficits and Dysfunction in the Insula Among Individuals With Intermittent Explosive Disorder. *Frontiers in Psychiatry*, *11*. https://doi.org/10.3389/fpsyt.2020.00439

Sethi, A., McCrory, E., Puetz, V., Hoffmann, F., Knodt, A. R., Radtke, S. R., Brigidi, B. D., Hariri, A. R., & Viding, E. (2018). Primary and Secondary Variants of Psychopathy in a Volunteer Sample Are Associated With Different Neurocognitive Mechanisms. *Biological Psychiatry: Cognitive Neuroscience and Neuroimaging*, *3*(12), 1013–1021. https://doi.org/10.1016/j.bpsc.2018.04.002

Shane, M. S., & Groat, L. L. (2018). Capacity for upregulation of emotional processing in psychopathy: All you have to do is ask. *Social Cognitive and Affective Neuroscience*, *13*(11), 1163–1176. https://doi.org/10.1093/scan/nsy088

Shao, R., & Lee, T. M. C. (2017). Are individuals with higher psychopathic traits better learners at lying? Behavioural and neural evidence. *Translational Psychiatry*, *7*(7). https://doi.org/10.1038/tp.2017.147

Shapiro, P. A., Sloan, R. P., Bagiella, E., Kuhl, J. P., Anjilvel, S., & Mann, J. J. (2000). Cerebral activation, hostility, and cardiovascular control during mental stress. *Journal of Psychosomatic Research*, *48*(4–5), 485–491. https://doi.org/10.1016/S0022-3999(00)00100-8

Sharp, C., Burton, P. C., & Ha, C. (2011). "Better the devil you know": A preliminary study of the differential modulating effects of reputation on reward processing for boys with and without externalizing behavior problems. *European Child and Adolescent Psychiatry*, *20*(11–12), 581–592. https://doi.org/10.1007/s00787-011-0225-x

Sheng, T., Gheytanchi, A., & Aziz-Zadeh, L. (2010). Default network deactivations are correlated with psychopathic personality traits. *PLoS ONE*, *5*(9), 1–7. https://doi.org/10.1371/journal.pone.0012611

Sitaram, R., Caria, A., Veit, R., Gaber, T., Ruiz, S., & Birbaumer, N. (2014). Volitional control of the anterior insula in criminal psychopaths using real-time fMRI neurofeedback: A pilot study. *Frontiers in Behavioral Neuroscience*, *8*(OCT), 1–13. https://doi.org/10.3389/fnbeh.2014.00344

Skibsted, A. P., Cunha-Bang, S. D., Carré, J. M., Hansen, A. E., Beliveau, V., Knudsen, G. M., & Fisher, P. M. (2017). Aggression-related brain function assessed with the Point Subtraction Aggression Paradigm in fMRI. *Aggressive Behavior*, *43*(6), 601–610. https://doi.org/10.1002/ab.21718

Soderstrom, H., Hultin, L., Tullberg, M., Wikkelso, C., Ekholm, S., & Forsman, A. (2002). Reduced frontotemporal perfusion in psychopathic personality. *Psychiatry Research - Neuroimaging*, *114*(2), 81–94. https://doi.org/10.1016/S0925-4927(02)00006-9

Soderstrom, H., Tullberg, M., Wikkelsö, C., Ekholm, S., & Forsman, A. (2000). Reduced regional cerebral blood flow in non-psychotic violent offenders. *Psychiatry Research - Neuroimaging*, *98*(1), 29–41. https://doi.org/10.1016/S0925-4927(99)00049-9

Soloff, P., White, R., & Diwadkar, V. A. (2014). Impulsivity, aggression and brain structure in high and low lethality suicide attempters with borderline personality disorder. *Psychiatry Research - Neuroimaging*, *222*(3). https://doi.org/10.1016/j.pscychresns.2014.02.006

Soloff, P. H., Abraham, K., Burgess, A., Ramaseshan, K., Chowdury, A., & Diwadkar, V. A. (2017). Impulsivity and aggression mediate regional brain responses in Borderline Personality Disorder: An fMRI study. *Psychiatry Research - Neuroimaging*, *260*, 76–85. https://doi.org/10.1016/j.pscychresns.2016.12.009

Soloff, P. H., Chiappetta, L., Mason, N. S., Becker, C., & Price, J. C. (2014). Effects of serotonin-2A receptor binding and gender on personality traits and suicidal behavior in borderline personality disorder. *Psychiatry Research - Neuroimaging*, *222*(3), 140–148. https://doi.org/10.1016/j.pscychresns.2014.03.008

Soloff, P. H., Price, J. C., Mason, N. S., Becker, C., & Meltzer, C. C. (2010). Gender, personality, and serotonin-2A receptor binding in healthy subjects. *Psychiatry Research - Neuroimaging*, *181*(1), 77–84. https://doi.org/10.1016/j.pscychresns.2009.08.007

Sommer, M., Sodian, B., Döhnel, K., Schwerdtner, J., Meinhardt, J., & Hajak, G. (2010). In psychopathic patients emotion attribution modulates activity in outcome-related brain areas. *Psychiatry Research - Neuroimaging*, *182*(2), 88–95. https://doi.org/10.1016/j.pscychresns.2010.01.007

Spoont, M. R., Kuskowski, M., & Pardo, J. V. (2010). Autobiographical memories of anger in violent and non-violent individuals: A script-driven imagery study. *Psychiatry Research - Neuroimaging*, *183*(3), 225–229. https://doi.org/10.1016/j.pscychresns.2010.06.004

Sterzer, P., Stadler, C., Krebs, A., Kleinschmidt, A., & Poustka, F. (2005). Abnormal neural responses to emotional visual stimuli in adolescents with conduct disorder. *Biological Psychiatry*, *57*(1), 7–15. https://doi.org/10.1016/j.biopsych.2004.10.008

Sterzer, P., Stadler, C., Poustka, F., & Kleinschmidt, A. (2007). A structural neural deficit in adolescents with conduct disorder and its association with lack of empathy. *NeuroImage*, *37*(1), 335–342. https://doi.org/10.1016/j.neuroimage.2007.04.043

Stevens, M. C., & Haney-Caron, E. (2012). Comparison of brain volume abnormalities between ADHD and conduct disorder in adolescence. *Journal of Psychiatry and Neuroscience*, *37*(6), 389–398. https://doi.org/10.1503/jpn.110148

Storvestre, G. B., Valnes, L. M., Jensen, A., Nerland, S., Tesli, N., Hymer, K.-E., Rosaeg, C., Server, A., Ringen, P. A., Jacobsen, M., Melle, I., & Haukvik, U. K. (2019). A preliminary study of cortical morphology in schizophrenia patients with a history of violence. *Psychiatry Research - Neuroimaging*, *288*, 29–36. https://doi.org/10.1016/j.pscychresns.2019.04.013

Strenziok, M., Krueger, F., Deshpande, G., Lenroot, R. K., Van der meer, E., & Grafman, J. (2011). Fronto-parietal regulation of media violence exposure in adolescents: A multi-method study. *Social Cognitive and Affective Neuroscience*, *6*(5), 537–547. https://doi.org/10.1093/scan/nsq079

Strenziok, M., Krueger, F., Heinecke, A., Lenroot, R. K., Knutson, K. M., van der Meer, E., & Grafman, J. (2011). Developmental effects of aggressive behavior in male adolescents assessed with structural and functional brain imaging. *Social Cognitive and Affective Neuroscience*, *6*(1), 2–11. https://doi.org/10.1093/scan/nsp036

Sun, X., Ma, R., Jiang, Y., Gao, Y., Ming, Q., Wu, Q., Dong, D., Wang, X., & Yao, S. (2018). MAOA genotype influences neural response during an inhibitory task in adolescents with conduct disorder. *European Child and Adolescent Psychiatry*, *27*(9), 1159–1169. https://doi.org/10.1007/s00787-018-1170-8

Suridjan, I., Boileau, I., Bagby, M., Rusjan, P. M., Wilson, A. A., Houle, S., & Mizrahi, R. (2012). Dopamine response to psychosocial stress in humans and its relationship to individual differences in personality traits. *Journal of Psychiatric Research*, *46*(7), 890–897. https://doi.org/10.1016/j.jpsychires.2012.03.009

Sutherland, M. T., & Fishbein, D. H. (2017). Higher trait psychopathy is associated with increased risky decision-making and less coincident insula and striatal activity. *Frontiers in Behavioral Neuroscience*, *11*. https://doi.org/10.3389/fnbeh.2017.00245

Swartz, J. R., Carranza, A. F., & Knodt, A. R. (2020). Amygdala activity to angry and fearful faces relates to bullying and victimization in adolescents. *Social Cognitive and Affective Neuroscience*, *14*(10), 1027–1035. https://doi.org/10.1093/scan/nsz084

Szabó, E., Kocsel, N., Édes, A., Pap, D., Galambos, A., Zsombók, T., Szabó, Á., Kozák, L. R., Bagdy, G., Juhász, G., Juhász, G., & Kökönyei, G. (2017). Callous-unemotional traits and neural responses to emotional faces in a community sample of young adults. *Personality and Individual Differences*, *111*, 312–317. https://doi.org/10.1016/j.paid.2017.02.026

Takahashi, K., Hosoya, T., Onoe, K., Takashima, T., Tanaka, M., Ishii, A., Nakatomi, Y., Tazawa, S., Takahashi, K., Doi, H., Wada, Y., & Watanabe, Y. (2018). Association between aromatase in human brains and personality traits. *Scientific Reports*, *8*(1). https://doi.org/10.1038/s41598-018-35065-4

Tang, Y., Jiang, W., Liao, J., Wang, W., & Luo, A. (2013). Identifying Individuals with Antisocial Personality Disorder Using Resting-State fMRI. *PLoS ONE*, *8*(4). https://doi.org/10.1371/journal.pone.0060652

Tessitore, A., Santangelo, G., De Micco, R., Vitale, C., Giordano, A., Raimo, S., Corbo, D., Amboni, M., Barone, P., & Tedeschi, G. (2016). Cortical thickness changes in patients with Parkinson's disease and impulse control disorders. *Parkinsonism and Related Disorders*, *24*, 119–125. https://doi.org/10.1016/j.parkreldis.2015.10.013

Thiel, A., Thiel, J., Oddo, S., Langnickel, R., Brand, M., Markowitsch, H. J., & Stirn, A. (2014). Obsessive-compulsive disorder patients with washing symptoms show a specific brain network when confronted with aggressive, sexual, and disgusting stimuli. *Neuropsychoanalysis*, *16*(2), 83–96. https://doi.org/10.1080/15294145.2014.976649

Thornton, L. C., Penner, E. A., Nolan, Z. T., Adalio, C. J., Sinclair, S., Meffert, H., Hwang, S., Blair, R. J. R., & White, S. F. (2017). The processing of animacy information is disrupted as a function of callous-unemotional traits in youth with disruptive behavior disorders. *NeuroImage: Clinical*, *16*, 498–506. https://doi.org/10.1016/j.nicl.2017.08.024

Tiihonen, J., Kuikka, J. T., Bergström, K. A., Karhu, J., Viinamäki, H., Lehtonen, J., Hallikainen, T., Yang, J., & Hakola, P. (1997). Single-photon emission tomography imaging of monoamine transporters in impulsive violent behaviour. *European Journal of Nuclear Medicine*, *24*(10), 1253–1260. https://doi.org/10.1007/s002590050149

Tiihonen, J., Rossi, R., Laakso, M. P., Hodgins, S., Testa, C., Perez, J., Repo-Tiihonen, E., Vaurio, O., Soininen, H., Aronen, H. J., Thompson, P. M., & Frisoni, G. B. (2008). Brain anatomy of persistent violent offenders: More rather than less. *Psychiatry Research - Neuroimaging*, *163*(3), 201–212. https://doi.org/10.1016/j.pscychresns.2007.08.012

Tikàsz, A., Potvin, S., Lungu, O., Joyal, C. C., Hodgins, S., Mendrek, A., & Dumais, A. (2016). Anterior cingulate hyperactivations during negative emotion processing among men with schizophrenia and a history of violent behavior. *Neuropsychiatric Disease and Treatment*, *12*, 1397–1410. https://doi.org/10.2147/NDT.S107545

Tikàsz, A., Potvin, S., Richard-Devantoy, S., Lipp, O., Hodgins, S., Lalonde, P., Lungu, O., & Dumais, A. (2018). Reduced dorsolateral prefrontal cortex activation during affective Go/NoGo in violent schizophrenia patients: An fMRI study. *Schizophrenia Research*, *197*, 249–252. https://doi.org/10.1016/j.schres.2017.11.011

Tonnaer, F., Siep, N., Van Zutphen, L., Arntz, A., & Cima, M. (2017). Anger provocation in violent offenders leads to emotion dysregulation. *Scientific Reports*, *7*(1). https://doi.org/10.1038/s41598-017-03870-y

van de Giessen, E., Rosell, D. R., Thompson, J. L., Xu, X., Girgis, R. R., Ehrlich, Y., Slifstein, M., Abi-Dargham, A., & Siever, L. J. (2014). Serotonin transporter availability in impulsive aggressive personality disordered patients: A PET study with [$^{11}$C]DASB. *Journal of Psychiatric Research*, *58*, 147–154. https://doi.org/10.1016/j.jpsychires.2014.07.025

Van den Bos, W., Vahl, P., Güroğlu, B., Van Nunspeet, F., Colins, O., Markus, M., Rombouts, S. A. R. B., Van der Wee, N., Vermeiren, R., & Crone, E. A. (2014). Neural correlates of social decision-making in severely antisocial adolescents. *Social Cognitive and Affective Neuroscience*, *9*(12), 2059–2066. https://doi.org/10.1093/scan/nsu003

Van den Stock, J., Hortensius, R., Sinke, C., Goebel, R., & de Gelder, B. (2015). Personality traits predict brain activation and connectivity when witnessing a violent conflict. *Scientific Reports*, *5*. https://doi.org/10.1038/srep13779

van Lith, K., Veltman, D. J., Cohn, M. D., Pape, L. E., van den Akker-Nijdam, M. E., van Loon, A. W. G., Bet, P., van Wingen, G. A., van den Brink, W., Doreleijers, T., Doreleijers, T., & Popma, A. (2018). Effects of Methylphenidate During Fear Learning in Antisocial Adolescents: A Randomized Controlled fMRI Trial. *Journal of the American Academy of Child and Adolescent Psychiatry*, *57*(12), 934–943. https://doi.org/10.1016/j.jaac.2018.06.026

Veit, R., Lotze, M., Sewing, S., Missenhardt, H., Gaber, T., & Birbaumer, N. (2010). Aberrant social and cerebral responding in a competitive reaction time paradigm in criminal psychopaths. *NeuroImage*, *49*(4), 3365–3372. https://doi.org/10.1016/j.neuroimage.2009.11.040

Verger, A., Klesse, E., Chawki, M. B., Witjas, T., Azulay, J.-P., Eusebio, A., & Guedj, E. (2018). Brain PET substrate of impulse control disorders in Parkinson's disease: A metabolic connectivity study. *Human Brain Mapping*, *39*(8), 3178–3186. https://doi.org/10.1002/hbm.24068

Veroude, K., von Rhein, D., Chauvin, R. J. M., van Dongen, E. V., Mennes, M. J. J., Franke, B., Heslenfeld, D. J., Oosterlaan, J., Hartman, C. A., Hoekstra, P. J., Glennon, J. C., & Buitelaar, J. K. (2016). The link between callous-unemotional traits and neural mechanisms of reward processing: An fMRI study. *Psychiatry Research - Neuroimaging*, *255*, 75–80. https://doi.org/10.1016/j.pscychresns.2016.08.005

Vetter, N. C., Backhausen, L. L., Buse, J., Roessner, V., & Smolka, M. N. (2020). Altered brain morphology in boys with attention deficit hyperactivity disorder with and without comorbid conduct disorder/oppositional defiant disorder. *Human Brain Mapping*, *41*(4), 973–983. https://doi.org/10.1002/hbm.24853

Viding, E., Sebastian, C. L., Dadds, M. R., Lockwood, P. L., Cecil, C. A. M., De Brito, S. A., & McCrory, E. J. (2012). Amygdala response to preattentive masked fear in children with conduct problems: The role of callous-unemotional traits. *American Journal of Psychiatry*, *169*(10), 1109–1116. https://doi.org/10.1176/appi.ajp.2012.12020191

Vieira, J. B., Ferreira-Santos, F., Almeida, P. R., Barbosa, F., Marques-Teixeira, J., & Marsh, A. A. (2014). Psychopathic traits are associated with cortical and subcortical volume alterations in healthy individuals. *Social Cognitive and Affective Neuroscience*, *10*(12), 1693–1704. https://doi.org/10.1093/scan/nsv062

Vieira, J. B., Tavares, T. P., Marsh, A. A., & Mitchell, D. G. V. (2017). Emotion and personal space: Neural correlates of approach-avoidance tendencies to different facial expressions as a function of coldhearted psychopathic traits. *Human Brain Mapping*, *38*(3), 1492–1506. https://doi.org/10.1002/hbm.23467

Vincent, G. M., Cope, L. M., King, J., Nyalakanti, P., & Kiehl, K. A. (2018). Callous-Unemotional Traits Modulate Brain Drug Craving Response in High-Risk Young Offenders. *Journal of Abnormal Child Psychology*, *46*(5), 993–1009. https://doi.org/10.1007/s10802-017-0364-8

Volkow, N. D., Tancredib, L. R., Grant, C., Gillespie, H., Valentine, A., Mullani, N., Wang, G.-J., & Hollister, L. (1995). Brain glucose metabolism in violent psychiatric patients: a preliminary study. *Psychiatry Research: Neuroimaging*, *61*(4), 243–253. https://doi.org/10.1016/0925-4927(95)02671-J

Volman, I., von Borries, A. K. L., Bulten, B. H., Verkes, R. J., Toni, I., & Roelofs, K. (2016). Testosterone modulates altered prefrontal control of emotional actions in psychopathic offenders. *eNeuro*, *3*(1), 52–60. https://doi.org/10.1523/ENEURO.0107-15.2016

von Polier, G. G., Greimel, E., Konrad, K., Großheinrich, N., Kohls, G., Vloet, T. D., Herpertz-Dahlmann, B., & Schulte-Rüther, M. (2020). Neural Correlates of Empathy in Boys With Early Onset Conduct Disorder. *Frontiers in Psychiatry*, *11*. https://doi.org/10.3389/fpsyt.2020.00178

Voon, V., Rizos, A., Chakravartty, R., Mulholland, N., Robinson, S., Howell, N. A., Harrison, N., Vivian, G., & Chaudhuri, K. R. (2014). Impulse control disorders in Parkinson's disease: Decreased striatal dopamine transporter levels. *Journal of Neurology, Neurosurgery and Psychiatry*, *85*(2), 148–152. https://doi.org/10.1136/jnnp-2013-305395

Völlm, B., Richardson, P., McKie, S., Elliott, R., Dolan, M., & Deakin, B. (2007). Neuronal correlates of reward and loss in Cluster B personality disorders: A functional magnetic resonance imaging study. *Psychiatry Research - Neuroimaging*, *156*(2), 151–167. https://doi.org/10.1016/j.pscychresns.2007.04.008

Völlm, B., Richardson, P., Stirling, J., Elliott, R., Dolan, M., Chaudhry, I., Del Ben, C., Mckie, S., Anderson, I., & Deakin, B. (2004). Neurobiological substrates of antisocial and borderline personality disorder: Preliminary results of a functional fMRI study. *Criminal Behaviour and Mental Health*, *14*(1), 39–54. https://doi.org/10.1002/cbm.559

Wallace, G. L., White, S. F., Robustelli, B., Sinclair, S., Hwang, S., Martin, A., & Blair, R. J. R. (2014). Cortical and subcortical abnormalities in youths with conduct disorder and elevated callous-unemotional traits. *Journal of the American Academy of Child and Adolescent Psychiatry*, *53*(4). https://doi.org/10.1016/j.jaac.2013.12.008

Weber, R., Ritterfeld, U., & Mathiak, K. (2006). Does playing violent video games induce aggression? Empirical evidence of a functional magnetic resonance imaging study. *Media Psychology*, *8*(1), 39–60. https://doi.org/10.1207/S1532785XMEP0801_4

White, S. F., Brislin, S. J., Sinclair, S., & Blair, J. R. (2014). Punishing unfairness: Rewarding or the organization of a reactively aggressive response? *Human Brain Mapping*, *35*(5), 2137–2147. https://doi.org/10.1002/hbm.22316

White, S. F., Fowler, K. A., Sinclair, S., Schechter, J. C., Majestic, C. M., Pine, D. S., & Blair, R. J. (2014). Disrupted expected value signaling in youth with disruptive behavior disorders to environmental reinforcers. *Journal of the American Academy of Child and Adolescent Psychiatry*, *53*(5), 579-588.e9. https://doi.org/10.1016/j.jaac.2013.12.023

White, S. F., Marsh, A. A., Fowler, K. A., Schechter, J. C., Adalio, C., Pope, K., Sinclair, S., Pine, D. S., & Blair, R. J. R. (2012). Reduced amygdala response in youths with disruptive behavior disorders and psychopathic traits: Decreased emotional response versus increased top-down attention to nonemotional features. *American Journal of Psychiatry*, *169*(7), 750–758. https://doi.org/10.1176/appi.ajp.2012.11081270

White, S. F., Pope, K., Sinclair, S., Fowler, K. A., Brislin, S. J., Williams, W. C., Pine, D. S., & Blair, R. J. R. (2013). Disrupted expected value and prediction error signaling in Youths with disruptive behavior disorders during a passive avoidance task. *American Journal of Psychiatry*, *170*(3), 315–323. https://doi.org/10.1176/appi.ajp.2012.12060840

White, S. F., Thornton, L. C., Leshin, J., Clanton, R., Sinclair, S., Coker-Appiah, D., Meffert, H., Hwang, S., & Blair, J. R. (2018). Looming Threats and Animacy: Reduced Responsiveness in Youth with Disrupted Behavior Disorders. *Journal of Abnormal Child Psychology*, *46*(4), 741–754. https://doi.org/10.1007/s10802-017-0335-0

White, S. F., Tyler, P. M., Erway, A. K., Botkin, M. L., Kolli, V., Meffert, H., Pope, K., & Blair, J. R. (2016). Dysfunctional representation of expected value is associated with reinforcement-based decision-making deficits in adolescents with conduct problems. *Journal of Child Psychology and Psychiatry and Allied Disciplines*, *57*(8), 938–946. https://doi.org/10.1111/jcpp.12557

White, S. F., Van Tieghem, M., Brislin, S. J., Sypher, I., Sinclair, S., Pine, D. S., Hwang, S., & Blair, R. J. R. (2016). Neural correlates of the propensity for retaliatory behavior in youths with disruptive behavior disorders. *American Journal of Psychiatry*, *173*(3), 282–290. https://doi.org/10.1176/appi.ajp.2015.15020250

White, S. F., Williams, W. C., Brislin, S. J., Sinclair, S., Blair, K. S., Fowler, K. A., Pine, D. S., Pope, K., & Blair, R. J. (2012). Reduced activity within the dorsal endogenous orienting of attention network to fearful expressions in youth with disruptive behavior disorders and psychopathic traits. *Development and Psychopathology*, *24*(3), 1105–1116. https://doi.org/10.1017/S0954579412000569

Wiggins, J. L., Brotman, M. A., Adleman, N. E., Kim, P., Oakes, A. H., Reynolds, R. C., Chen, G., Pine, D. S., & Leibenluft, E. (2016). Neural correlates of irritability in disruptive mood dysregulation and bipolar disorders. *American Journal of Psychiatry*, *173*(7), 722–730. https://doi.org/10.1176/appi.ajp.2015.15060833

Witte, A. V., Flöel, A., Stein, P., Savli, M., Mien, L.-K., Wadsak, W., Spindelegger, C., Moser, U., Fink, M., Hahn, A., Kasper, S., & Lanzenberger, R. (2009). Aggression is related to frontal serotonin-1A receptor distribution as revealed by PET in healthy subjects. *Human Brain Mapping*, *30*(8), 2558–2570. https://doi.org/10.1002/hbm.20687

Woermann, F. G., Van Elst, L. T., Koepp, M. J., Free, S. L., Thompson, P. J., Trimble, M. R., & Duncan, J. S. (2000). Reduction of frontal neocortical grey matter associated with affective aggression in patients with temporal lobe epilepsy: An objective voxel by voxel analysis of automatically segmented MRI. *Journal of Neurology Neurosurgery and Psychiatry*, *68*(2), 162–169. https://doi.org/10.1136/jnnp.68.2.162

Yang, Y., Raine, A., Colletti, P., Toga, A. W., & Narr, K. L. (2010). Morphological alterations in the prefrontal cortex and the amygdala in unsuccessful psychopaths. *Journal of Abnormal Psychology*, *119*(3), 546–554. https://doi.org/10.1037/a0019611

Yang, Y., Raine, A., Han, C.-B., Schug, R. A., Toga, A. W., & Narr, K. L. (2010). Reduced hippocampal and parahippocampal volumes in murderers with schizophrenia. *Psychiatry Research - Neuroimaging*, *182*(1), 9–13. https://doi.org/10.1016/j.pscychresns.2009.10.013

Yang, Y., Raine, A., Narr, K. L., Colletti, P., & Toga, A. W. (2009). Localization of deformations within the amygdala in individuals with psychopathy. *Archives of General Psychiatry*, *66*(9), 986–994. https://doi.org/10.1001/archgenpsychiatry.2009.110

Yang, Y., Wang, P., Baker, L. A., Narr, K. L., Joshi, S. H., Hafzalla, G., Raine, A., & Thompson, P. M. (2015). Thicker temporal cortex associates with a developmental trajectory for psychopathic traits in adolescents. *PLoS ONE*, *10*(5). https://doi.org/10.1371/journal.pone.0127025

Yang, Y. K., Yao, W. J., Yeh, T. L., Lee, I. H., Chen, K. C., & Lu, R. B. (2007). Association between serotonin transporter availability and hostility scores in healthy volunteers-A single photon emission computed tomography study with [$^{123}$I] ADAM. *Psychiatry Research - Neuroimaging*, *154*(3), 281–284. https://doi.org/10.1016/j.pscychresns.2006.11.010

Yang, Y., Raine, A., Lencz, T., Bihrle, S., Lacasse, L., & Colletti, P. (2005). *Volume Reduction in Prefrontal Gray Matter in Unsuccessful Criminal Psychopaths*. https://doi.org/10.1016/j.biopsych.2005.01.021

Yoder, K. J., Harenski, C., Kiehl, K. A., & Decety, J. (2015). Neural networks underlying implicit and explicit moral evaluations in psychopathy. *Translational Psychiatry*, *5*(8). https://doi.org/10.1038/tp.2015.117

Yoder, K. J., Harenski, C., Kiehl, K. A., & Decety, J. (2021). Neural responses to morally laden interactions in female inmates with psychopathy. *NeuroImage: Clinical*, *30*. https://doi.org/10.1016/j.nicl.2021.102645

Yoder, K. J., Porges, E. C., & Decety, J. (2015). Amygdala subnuclei connectivity in response to violence reveals unique influences of individual differences in psychopathic traits in a nonforensic sample. *Human Brain Mapping*, *36*(4), 1417–1428. https://doi.org/10.1002/hbm.22712

Zhang, J., Cao, W., Wang, M., Wang, N., Yao, S., & Huang, B. (2019). Multivoxel pattern analysis of structural MRI in children and adolescents with conduct disorder. *Brain Imaging and Behavior*, *13*(5), 1273–1280. https://doi.org/10.1007/s11682-018-9953-6

Zhang, J., Liu, W., Zhang, J., Wu, Q., Gao, Y., Jiang, Y., Gao, J., Yao, S., & Huang, B. (2018). Distinguishing adolescents with conduct disorder from typically developing youngsters based on pattern classification of brain structural MRI. *Frontiers in Human Neuroscience*, *12*. https://doi.org/10.3389/fnhum.2018.00152

Zhang, L., Kerich, M., Schwandt, M. L., Rawlings, R. R., McKellar, J. D., Momenan, R., Hommer, D. W., & George, D. T. (2013). Smaller right amygdala in Caucasian alcohol-dependent male patients with a history of intimate partner violence: A volumetric imaging study. *Addiction Biology*, *18*(3), 537–547. https://doi.org/10.1111/j.1369-1600.2011.00381.x

Zhang, Y.-D., Zhou, J.-S., Lu, F.-M., & Wang, X.-P. (2019). Reduced gray matter volume in male adolescent violent offenders. *PeerJ*, *2019*(9). https://doi.org/10.7717/peerj.7349

Zhu, W., Zhou, X., & Xia, L.-X. (2019). Brain structures and functional connectivity associated with individual differences in trait proactive aggression. *Scientific Reports*, *9*(1). https://doi.org/10.1038/s41598-019-44115-4

Zhu, Y., Ying, K., Wang, J., Su, L., Chen, J., Lin, F., Cai, D., Zhou, M., Wu, D., Guo, C., Guo, C., & Wang, S. (2014). Differences in functional activity between boys with pure oppositional defiant disorder and controls during a response inhibition task: A preliminary study. *Brain Imaging and Behavior*, *8*(4), 588–597. https://doi.org/10.1007/s11682-013-9275-7

Zijlmans, J., Marhe, R., Bevaart, F., Luijks, M. A., van Duin, L., Tiemeier, H., & Popma, A. (2018). Neural correlates of moral evaluation and psychopathic traits in male multi-problem young adults. *Frontiers in Psychiatry*, *9*(JUN). https://doi.org/10.3389/fpsyt.2018.00248
